# Large-Scale Restructuring of the Caspase-1 Gene Cluster Region in Mammals

**DOI:** 10.1101/2025.06.19.660630

**Authors:** David Stevens, Tasman Daish, Frank Grützner

**Affiliations:** School of Biological Sciences, The University of Adelaide, Adelaide, 5005 SA, Australia

**Keywords:** Inflammasome, Phylogenetics, Mammalian gene evolution, Comparative Immunology/Evolution, Gene Rearrangement, Inflammation

## Abstract

The inflammatory response is an important component of the host immune defence mechanism against pathogens. Activation of the NLRP3 inflammasome has been shown to have a wide range of triggers including fungal, bacterial and viral components, cellular stress and environmental irritants. The NLRP3 inflammasome has been well characterised in mouse and humans but limited information is available from other mammalian species. In order to gain a better understanding of the evolution of genes involved in the NLRP3 inflammasome pathway, we examined them in mammalian species representing the three major groups (eutheria, metatheria and prototheria) and in chicken as an outgroup. Our results show that the inflammasome pathway machinery is generally well conserved though chicken appears to lack several key components and monotremes have a duplication of *Syk*. Analysis of the proinflammatory caspase cluster and neighboring genes revealed massive reorganisation as well as the previously described multiple duplications that occurred during mammalian evolution. Our data suggest that *Caspase-1* moved to a new chromosomal region in early mammalian evolution. This was followed by expansion of the cluster and accumulation of additional genes regulating inflammatory responses such as *Card16*, *Card17*, *Card18* and the *Birc* genes. The expansion of key gene families flanking *Caspase-1* may have led to an expansion of inflammasome pathways and a more regulated immune system in humans through the CARD genes.

## INTRODUCTION

Inflammasomes are multiprotein complexes, which generally consist of a sensor and an adaptor serving as a scaffold that promotes Caspase-1 autocatalytic cleavage and processing (Martinon et al. 2002). Inflammasomes are known to play essential roles in the host response to pathogen-associated molecular patterns (PAMPs) and damage-associated molecular patterns (DAMPs). There have been several inflammasomes described to date, four of which are formed by Nod-like Receptor (NLR) family members: NLR family, pyrin domain containing 1 (NLRP1) (Martinon et al. 2002), NLRP3 (Agostini et al. 2004), NLRP6 (Leng et al. 2020) and NLRC4 (Mariathasan et al. 2004) as well as the HIN-200 family member AIM2 (Bürckstümmer et al. 2009; Fernandes-Alnemri et al. 2009; Hornung et al. 2009) and MFEV (Pyrin) (Akbaba et al. 2021; Xu et al. 2014). The NLRP3 inflammasome has been shown to form in response to a variety of stimuli including PAMPs such as lipopolysaccharide (LPS) and DAMPs such as ATP (Dostert et al. 2008; Gao et al.; Gross et al. 2009; Kanneganti et al. 2006; Martinon et al. 2006; Rodrigues et al. 2020; Zeng et al. 2022). Here we investigate the evolution of genes involved in pathways relating to gram negative bacteria (Kanneganti et al. 2006) and fungi (Gross et al. 2009).

Caspases are a class of cysteine proteases which cleave proteins with caspase-specific target motifs following an aspartate residue (Cerretti et al. 1992). Generally, caspases are ubiquitously expressed and exist as inactive zymogens (termed pro-caspases) which contain a prodomain and sites involved in cleavage and dimerization. Caspases are considered initiators of apoptosis (caspases-2,-8,-9 and-10), effectors of apoptosis (caspases-3,-6 and-7) or proinflammatory proteins (caspases-1,-4,-5,-11,-12 and-13) (Degterev et al. 2003; Eckhart et al. 2008). The proinflammatory caspase subfamily is also referred to as the *Caspase-1* subfamily as all the members are in close proximity in a cluster on the same chromosome. Gene duplication has shaped diversity of the caspase genes where *Caspase-11* and *Caspase-12* are likely to be the result of duplications of *Caspase-1* and *Caspase-5* a duplication of *Caspase-4* (Eckhart et al. 2008). While the main function of the proinflammatory caspases is to promote the inflammatory response, there is evidence that certain members are also capable of effecting apoptosis (Wang et al. 1998). *Caspase-12* appears to be unique in that it acts as a negative regulator of the pathway (Saleh et al. 2006).

Knowledge of the molecular and biochemical mechanisms underlying the inflammasome response is increasing but information on the evolution of the pathway is still limited to human and mouse systems. The Caspase-1 subfamily has undergone gene expansion during evolution (Eckhart et al. 2008), associated with increased complexity of the inflammatory response. *Tumour Necrosis Factor* (*TNF*) (Glenney and Wiens 2007; Roca et al. 2008) and the *Toll-like Receptors* (*TLR*s) (Alvarez-Pellitero 2008) are present in multiple species outside of mammals but knowledge of the other inflammasome pathway genes in other species is very limited. NLR family members have been examined extensively in rodents and primates (Hughes 2006; Pétrilli and Martinon 2011; Tian et al. 2009) with further work expanding this to include dog, cow and chicken (Hughes 2006; Tian et al. 2009). Platypus (using the OANA5 assembly) and opossum reproductive NLRPs have been previously examined using genbank sequences (Duenez-Guzman and Haig 2014). Their combined works show species specific expansions for several of the NLR members as well as deletion of certain members in some lineages. These analyses do not include representatives of marsupial and monotreme lineages.

Given the important role of the inflammasome pathway in the initiation of the inflammatory response, we studied eight mammalian species (covering the three major mammalian lineages: eutheria, metatheria and prototheria) and chicken to develop a better understanding of how genes in this system have evolved in mammals. The recently released echidna genome and the improved platypus genome assemblies provides a unique opportunity to assess the evolution of genes involved in the NLRP3 inflammasome pathway (Zhou et al. 2025; Zhou et al. 2021). Here we show that the majority of the genes involved in the NLRP3 inflammasome pathway are conserved through amniote evolution and provide evidence for an expansion of key gene families which may have led to the development of the non-canonical inflammasome pathways providing alternate pathways to induce a response as well as an increase the regulatory elements through the CARD genes.

## MATERIALS AND METHODS

### Bioinformatic analysis; sequence queries, alignments and synteny analysis

A combination of BLAST and synteny analysis facilitated determination of the conservation status of the different inflammasome pathway components in multiple mammalian species and chicken (Table I). BLAST searches were performed against each species individually using default settings and human gene sequences as a reference (Online Resource 1). All sequences and non-monotreme genomes used were sourced from NCBI (http://www.ncbi.nlm.nih.gov/). For the platypus (mOrnAna1) and echidna (mTacAcu1) assemblies the flanking sequence of the BLAST results were extracted and GENSCAN (Burge and Karlin 1997; Burge and Karlin 1998) used to predict putative genes in that region followed by tBLATn against the human RNA Refseq library to determine likely orthologues. Putative platypus and echidna sequences can be found in Online Resource 2. Synteny analysis was performed using eutherian chromosome alignments as references. Predicted protein sequences from examined species were aligned in MEGAX (Kumar et al. 2018) using MUSCLE (Edgar 2004) and Neighbour-joining phylogenetic trees were generated using the p-distance method with 1000 bootstrap repeats and pairwise deletion. The ExPASy Prosite tool (de Castro et al. 2006; Sigrist et al. 2002; Sigrist et al. 2013; Sigrist et al. 2005) was used to predict protein domains and active sites. The accession numbers of the sequences used for this analysis can be found in Online Resource 1.

**Table 1.**
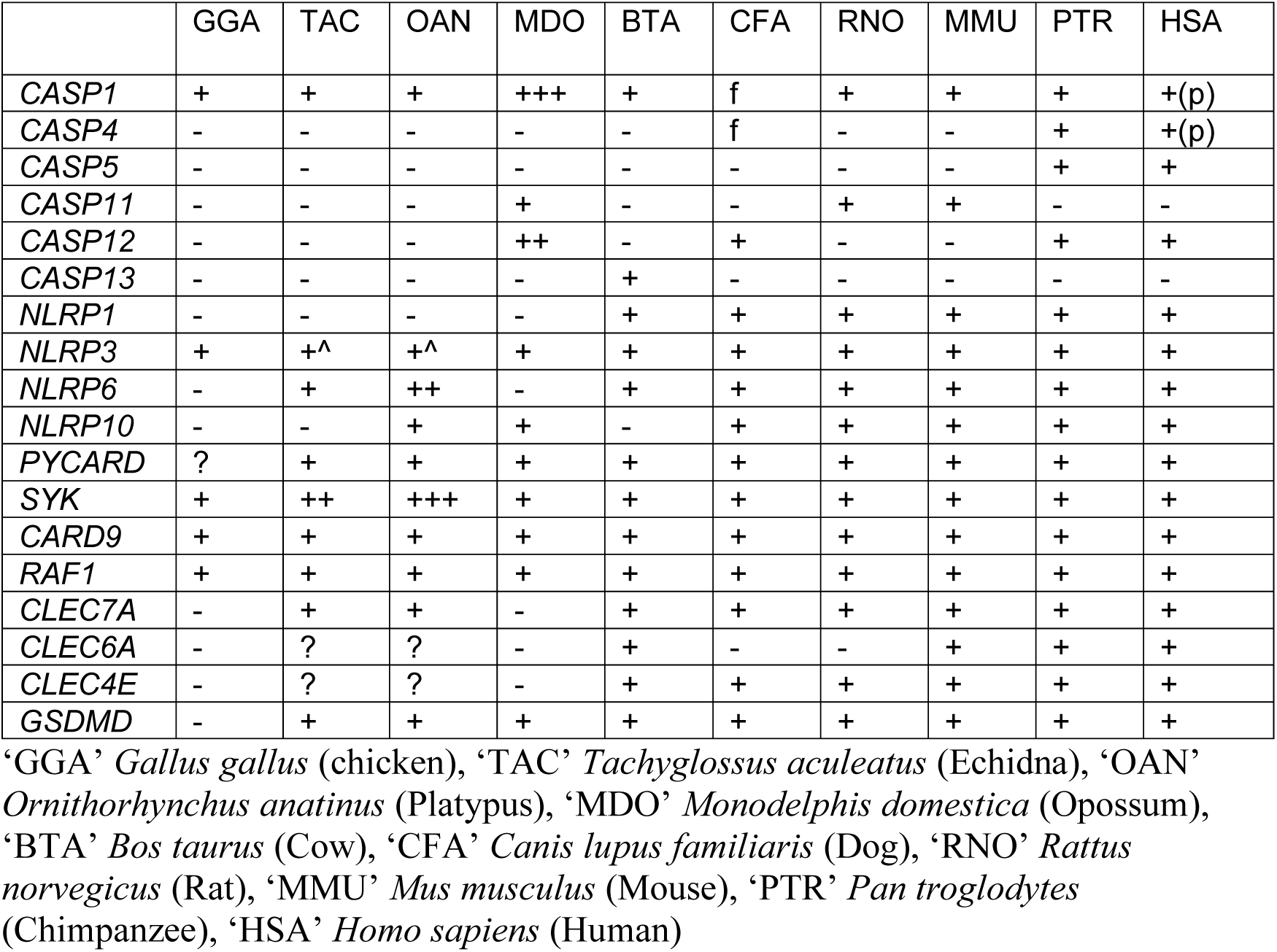
Inflammasome pathway genes in amniotes. +: present,-: absent, f: fusion described previously (Eckhart et al. 2008). p: pseudogene, ?: undetermined, and ^: needs to be confirmed through experimental analysis. Other genes which are conserved have not been included in this figure.

### DNA preparation and polymerase chain reactions (PCRs)

Genomic DNA was prepared from tissue using a standard phenol:chloroform protocol. Total RNA was prepared from frozen tissue using TRIzol (Invitrogen, Carlsbad, CA, USA) according to the manufacturer’s instructions. cDNA was produced by reverse transcription using a SuperScript III First-Strand Synthesis Kit (Invitrogen, Carlsbad, CA, USA). PCR (94°C 4 min, 25-35 cycles of 94°C 30 sec, 55-60°C 1 min, 72°C 1 min, with a final 72°C 4 min) was used with annealing temperature and number of cycles determined by the primers used (Online Resource 3). For degenerate primers the number of cycles was increased to 45. PCR products were purified and prepared using a standard dye terminator protocol then sequenced by the Australian Genome Research Facility (AGRF). Results were analysed using Chromas (http://www.technelysium.com.au/chromas.html).

## RESULTS

### Evolution of inflammasome pathway components in vertebrates

Generally, the inflammasome pathway is evolutionarily conserved except lineage specific duplications for example in caspase genes. Here we embarked on a more comprehensive analysis which confirmed conservation of many genes in mammals but also revealing unexpected changes in several components of this pathway (Table I).

### Dectin family receptors

*Dectin*-1 (*Clec7a*), *Dectin-2* (*Clec6a*), *Dectin-3* (*Clec4d*) and *Mincle* (*Clec4e*) belong to a group of immunoreceptor tyrosine-based activation motif (ITAM)-containing or ITAM-coupled C-type lectins (Goodridge et al. 2011). These receptors have been shown to be essential in humans and mice for the activation of Syk-dependent and independent pathways in response to fungi (Gross et al. 2006; Sato et al. 2006; Steele et al. 2005; Zhu et al. 2013). The four genes are located within the natural killer cluster in two different clusters. The Dectin-1 cluster consists of *Dectin-1*, *Clec1*, *Clec2* (*Clec1b*), *DNGR1* (*Clec9a*), *Micl* (*Clec12a*), *Mah* (*Clec12b*) and *Lox1* (*Olr1*) (Online Resource 4a). The Dectin-2 cluster consists of *Dectin-2*, *DCIR* (*Clec4a*), *DCAR* (*Clec4b1*), *BDCA-2* (*Clec4c*), *Mincle* (*Clec4e*) and *Dectin-3* (Online Resource Figure 4b). Some species-specific variations are known; *Clec4c* is only found in humans and chimpanzees, while *Clec4b1/2* is specific to rodents (Malamud and Brown 2024). We examined this family in monotremes and marsupials to better understand the evolution of these genes in mammals.

Our analysis shows that in monotremes the Dectin family genes are split across two chromosomes where the Dectin-1 subfamily co-locates with the natural killer cluster (Chr17 in platypus and scaffold_11 in echidna) while sequences with high similarity to the Dectin-2 subfamily are located on ChrX4 in platypus and scaffold_55 in echidna.

For both species, the Dectin-1 cluster contains eight genes while the second has only three. These genes have been labelled as *Clec-like* with 1-8 covering those on platypus Chr17 and echidna scaffold_11 and 9-11 for the genes located on platypus ChrX4 and echidna scaffold_55. In most eutherians examined the Dectin-1 subfamily contains 7 genes with 8 genes found in cow (Online Resource 4a). The genes also show similar organisation on chromosomes apart from human and chimpanzee which appear to have flipped *Clec1b* and *Clec12b*. Our data suggests that monotreme *Clec-like* genes 1 and 3 are most similar to *Clec12b* while *Clec-like* 2 appears similar to *Clec12a*, *Clec1b* and *Clec12b* (Online Resource 5). This is similar to what is observed in cow. The remaining genes showed similarity to human *Clec1b* (*Clec-like* 4), *Clec9a* (*Clec-like* 5), *Clec1a* (*Clec-like* 6), *Clec7a* (*Clec-like* 7) and *Olr1* (*Clec-like* 8) reflecting the arrangement seem in the eutherians.

The second cluster in monotremes is flanked by *Necap* and *C3ar1* which are consistently found flanking the Dectin-2 subfamily in eutherians supporting initial BLAST results suggesting the three genes belong to this subfamily. In platypus, *Clec-like* 9 showed highest similarity to human *Clec4d* with lower coverage (Online Resource 5a). It also showed similarity to *Clec6a* and *Clec4a*. However, in echidna *Clec-like* 9 showed similarity to human *Clec6a*, *Clec4c* and *Clec4d* (Online Resource 5b). In platypus, *Clec-like* 10 showed similarity to human *Clec4a*, *Clec4d* and *Clec4c*. Echidna *Clec-like* 10 and platypus *Clec-like* 11 appears most similar to human *Clec4e* while echidna *Clec-like* 11 appears most similar to *Clec4d*.

Phylogenetic analysis shows some unexpected groupings such as chimpanzee *Clec12a* grouping with the *Clec7a* genes, cow *Clec6a* grouping with *Clec9* and very low bootstrap values (Online Resource 6). Interestingly, in opossum we identified only three Dectin family genes; *Clec1a*, *Clec1b* and *Clec4e* suggesting marsupials may have lost most of this gene family.

### Duplication of spleen tyrosine kinase (Syk) in monotremes

Next we investigated spleen tyrosine kinase (*Syk*) which plays an essential role in inflammasome activation in response to the fungi *C. albicans* and *S. cerevisiae* (Gross et al. 2009).

*Syk* was identified in chicken and is found in all therian mammals (human, chimpanzee, mouse, rat, dog, cow and opossum) examined. In the OANA5 platypus assembly *Syk* was absent with only a partial sequence present which appeared more similar to *Zap70*, a member of the same tyrosine kinase family. We re-examined this result with the new assemblies to determine if this result was accurate. The echidna genome contains a copy of *Zap70* on scaffold_1 flanked upstream by *Adamts10* and *Myo1f* as in therians however *Tmem131* was found on scaffold_33 (Fig 1, Online Resource 7b,d).

**Fig 1.**
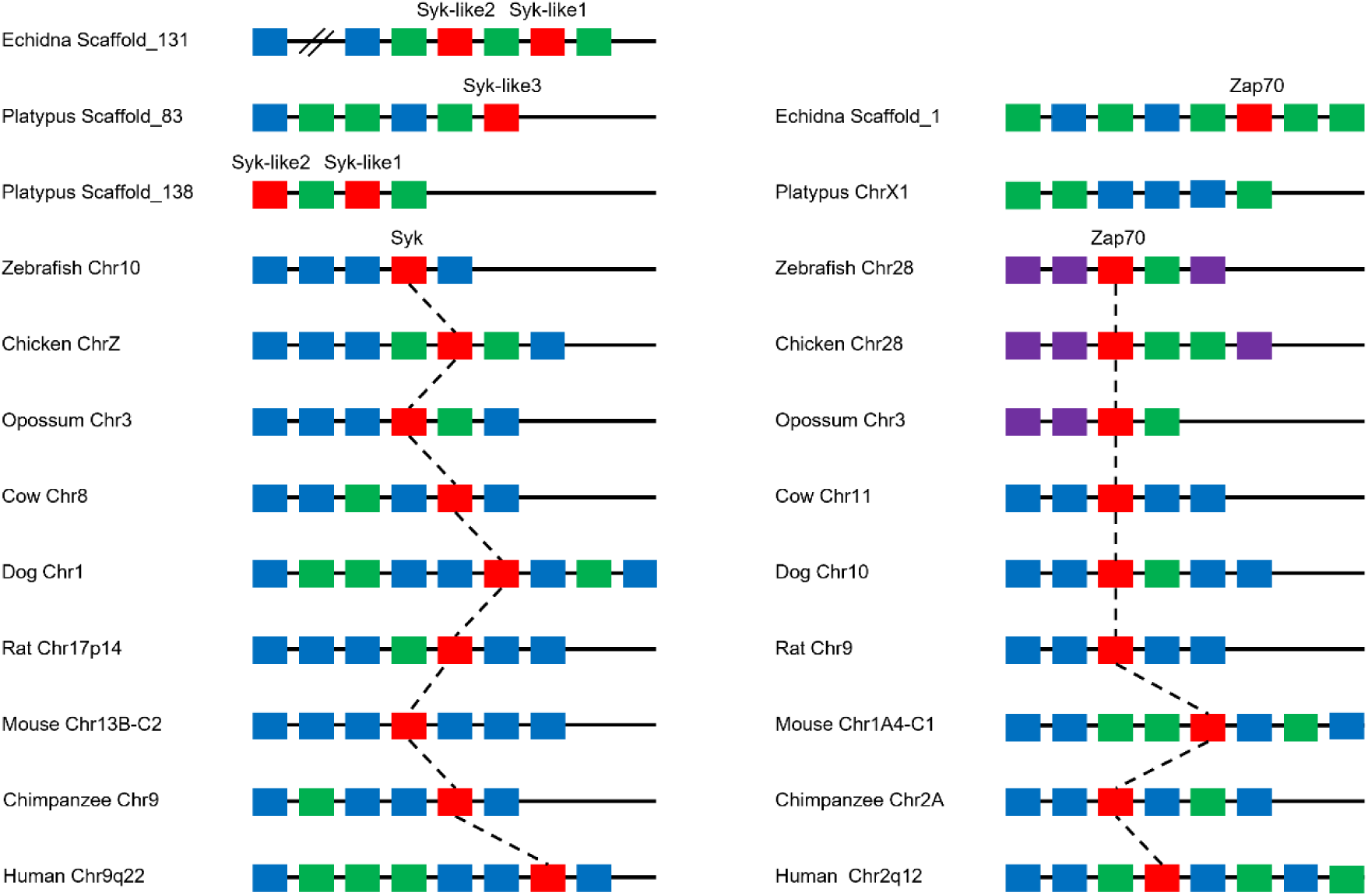
Chromosomal arrangements of *Syk* and *Zap70* in multiple mammalian species with chicken and zebrafish as non-mammalian outgroups. Red boxes represent the gene of interest *Syk* or *Zap70*, blue and purple boxes represent syntenic genes while green represents those that are species specific

Surprisingly two copies of *Syk* were identified on echidna scaffold_131 (Online Resource 7a). *Syk-like1* is flanked downstream by *Auh* and *Nfil3* which is consistent with *Syk* in therians (Online Resource 7a,c). *Syk-like2* is located upstream of *Syk-like1* with an unknown gene in between. Of the two genes commonly found upstream of *Syk* in therians, *Diras2* was identified on scaffold_17 while *Gadd45g* appears to be absent. In platypus three potential *Syk* were identified (Online Resource 7a). As with echidna, two of the genes (*Syk-like1* and *Syk-like2*) are located next to each other (scaffold_138) while the third (*Syk-like3*), was found on scaffold_83, and only contained the 5’ end of the gene. Interestingly the flanking genes *Auh* and *Nfil3* were also identified on scaffold_83 along with *Syk-like3*. *Diras2* is located on chr18 while *Ror2* and *Gadd45g* are absent. No putative *Zap70* homologue was found in the platypus genome assembly while the flanking genes *Myo1f* and *Adamts10* were found on chrX1 and *Tmem131* on chr18.

### NLRP3 is conserved in amniotes

NLRP3 functions as the inflammasome scaffold through NACHT domain interactions (Mariathasan et al. 2006). The LRR sequence of the NLRs is thought to function as a PAMP sensor and the interaction of these PAMPs with the LRR region is thought to trigger dimerization and inflammasome formation (Kanneganti et al. 2007). Like the caspases, the NLRP genes appear to be divided in to two roles: reproduction and inflammation (Tian et al. 2009). While a previous study has examined this family in the platypus and opossum (Duenez-Guzman and Haig 2014) we decided to re-examine the conservation of the inflammatory associated NLRPs in the new monotreme assemblies to further elucidate the evolution of this subset of genes (Fig 2).

**Fig 2.**
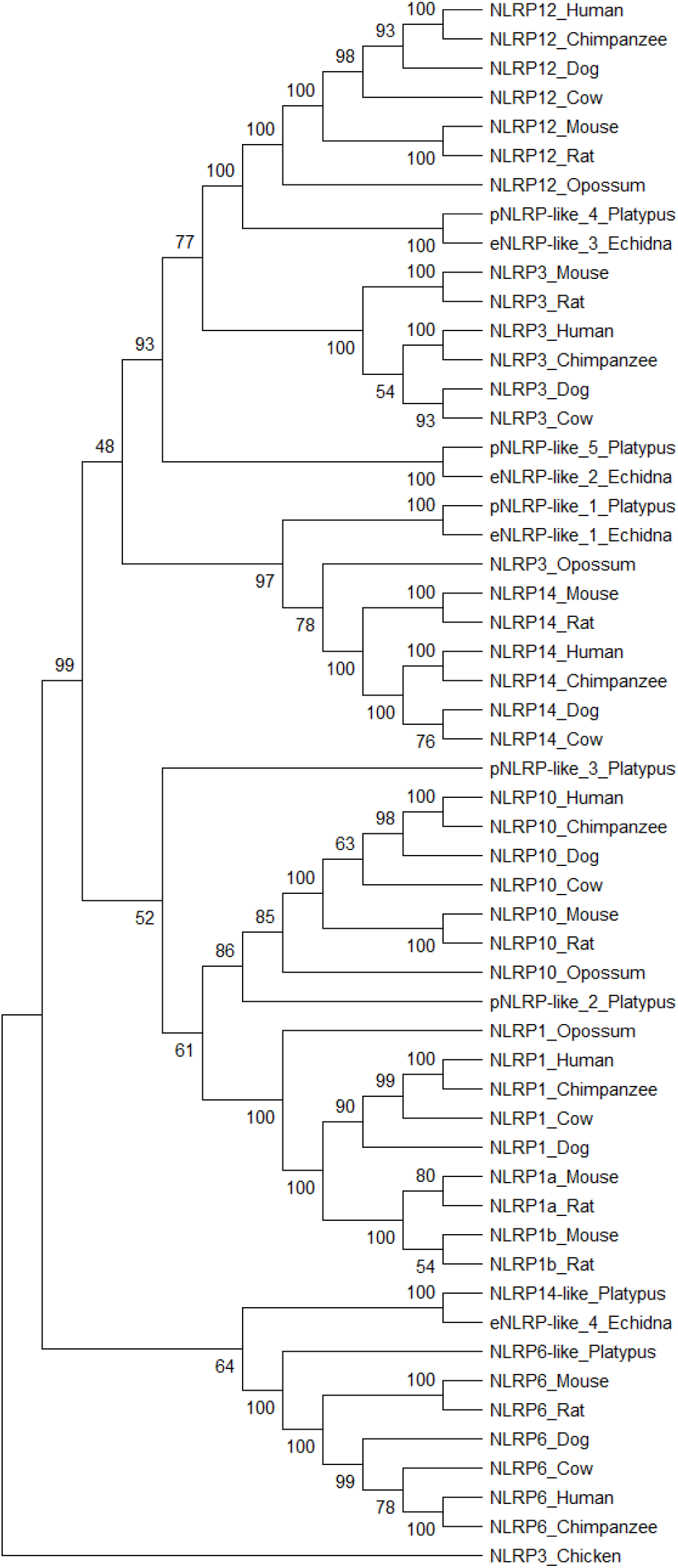
Phylogenetic analysis of NLRP and NLRP-like genes associated with the inflammatory response. Predicted protein sequences from multiple mammalian species and chicken as an outgroup were aligned in MEGAX using MUSCLE and Neighbour-joining phylogenetic trees were generated using the p-distance method with 1000 bootstrap repeats and pairwise deletion

Seven *NLRP-like* genes were identified in the platypus assembly and four in echidna. The putative genes on platypus chr1 (*pNLRP-like 1*) and echidna scaffold_42 (*eNLRP-like 1*) are homologous based on phylogenetic analysis (Figure 2). However, reciprocal BLAST suggests *pNLRP-like 1* is most similar to human *NLRP12* while *eNLRP-like 1* is most similar to human *NLRP3* (Online Resource 5). Interestingly in the phylogenetic tree these genes cluster with Opossum *NLRP3* which is located in proximity of the eutherian *NLRP14* homologues.

The two genes on platypus chr2 (*pNLRP-like 2* and *pNLRP-like 3*) show highest similarity to human *NLRP12* and *NLRP3* respectively (Online Resource 5a).

Phylogenetic analysis suggests that *pNLRP-like 2* is *NLRP10* while *pNLRP-like 3* is located on a separate node prior to an apparent split that led to the rise of *NLRP1* and *NLRP10* (Fig 2). Synteny supports *pNLRP-like 2* as being the ortholog of *NLRP10*. To determine whether pNLRP-like 3 is *NLRP1* a search was performed for *Mis12*, a gene commonly found to flank *NLRP1* in therians. *Mis12* was found on chr17 where no *NLRP-like* genes were identified suggesting *pNLRP-like 3* is not *NLRP1*. Interestingly, there appears to be no homologous genes in echidna although whether this is the result of a loss in echidna, a gain in platypus or the result of assembly quality in echidna is currently unclear.

The two *NLRP-like* genes on platypus chr10; *pNLRP-like 4* and *pNLRP-like 5* show high similarity to human *NLRP12* and *NLRP3* respectively (Online Resource 5a). Based on phylogenetic analysis the homologous genes in echidna are *eNLRP-like 3* (highest similarity was human *NLRP12*) and *eNLRP-like 2* (highest similarity was human *NLRP3*) which are located on echidna scaffold_45 (Fig2, Online Resource 5b). While the *pNLRP-like 4* and *eNLRP-like 3* indeed cluster on the phylogenetic tree with therian *NLRP12*, *pNLRP-like 5* and *eNLRP-like 2* form their own cluster separate from the *NLRP3* and *NLRP12* clusters. The presence of *Myadm* on platypus chr10 and echidna scaffold_45 further supports this data as the gene is consistently found flanking *NLRP12* however, in monotremes it does not appear to directly flank the gene.

The *NLRP-like* gene found on platypus chr3 is flanked by *Pgghg* and *Psmd13* which flank *NLRP6* in therians and this is further supported by both reciprocal BLAST against the human genome assembly and phylogenetic analysis (Fig 2, Online Resource 5a). In echidna the two flanking genes were found on scaffold_13, but no NLRP-like gene was identified in the region. Interestingly the NLRP-like genes located on platypus chr6 (*pNLRP-like 6*) and echidna scaffold_8 (*eNLRP-like 4*) also cluster with *NLRP6* on the phylogenetic tree although reciprocal BLAST against the human genome assembly suggests they are most likely *NLRP3* (Online Resource 5).

Analysis of the opossum inflammatory *NLRP-like* genes suggest the marsupial has *NLRP1*, *NLRP10*, *NLRP12* which group with the other mammalian equivalents whilst the putative *NLRP3* appears to cluster with eutherian *NLRP14*. However, it appears to lack *NLRP6* suggesting the gene may have been lost in marsupials.

### Potential loss of PYCARD in birds

PYCARD (ASC) is an adaptor protein that contains both a CARD and a pyrin (PYD) domain, allowing it to interact with the inflammasome sensor proteins and pro-Caspase-1. Recruitment and oligomerisation of PYCARD is essential for the maturation of Caspase-1 in inflammasomes formed by sensor proteins that lack a CARD such as NLRP3 (Mariathasan et al. 2004). In inflammasomes with CARD containing sensor proteins a second PYCARD-independent pathway has been observed (Broz et al. 2010). Despite being able to interact directly with pro-Caspase-1, processing of Caspase-1 in the NLRP1 inflammasome is greatly increased when PYCARD is also present (Faustin et al. 2007).

*Pycard* was identified in all species examined except for chicken. In monotremes *Pycard* can be found amongst the same flanking genes as those in therian mammals (Fig 3a). Interestingly in platypus there appears to be another *Pycard-like* gene on chr2 however it looks to be a pseudogene as it contains only part of the second exon of *Pycard*. The genes flanking *Pycard* in mammalian genomes were also absent in chicken (Fig 3a). To investigate whether this loss is common to the bird lineage we searched the zebrafinch genome for *Pycard* but were unable to identify the gene (not shown).

**Fig 3.**
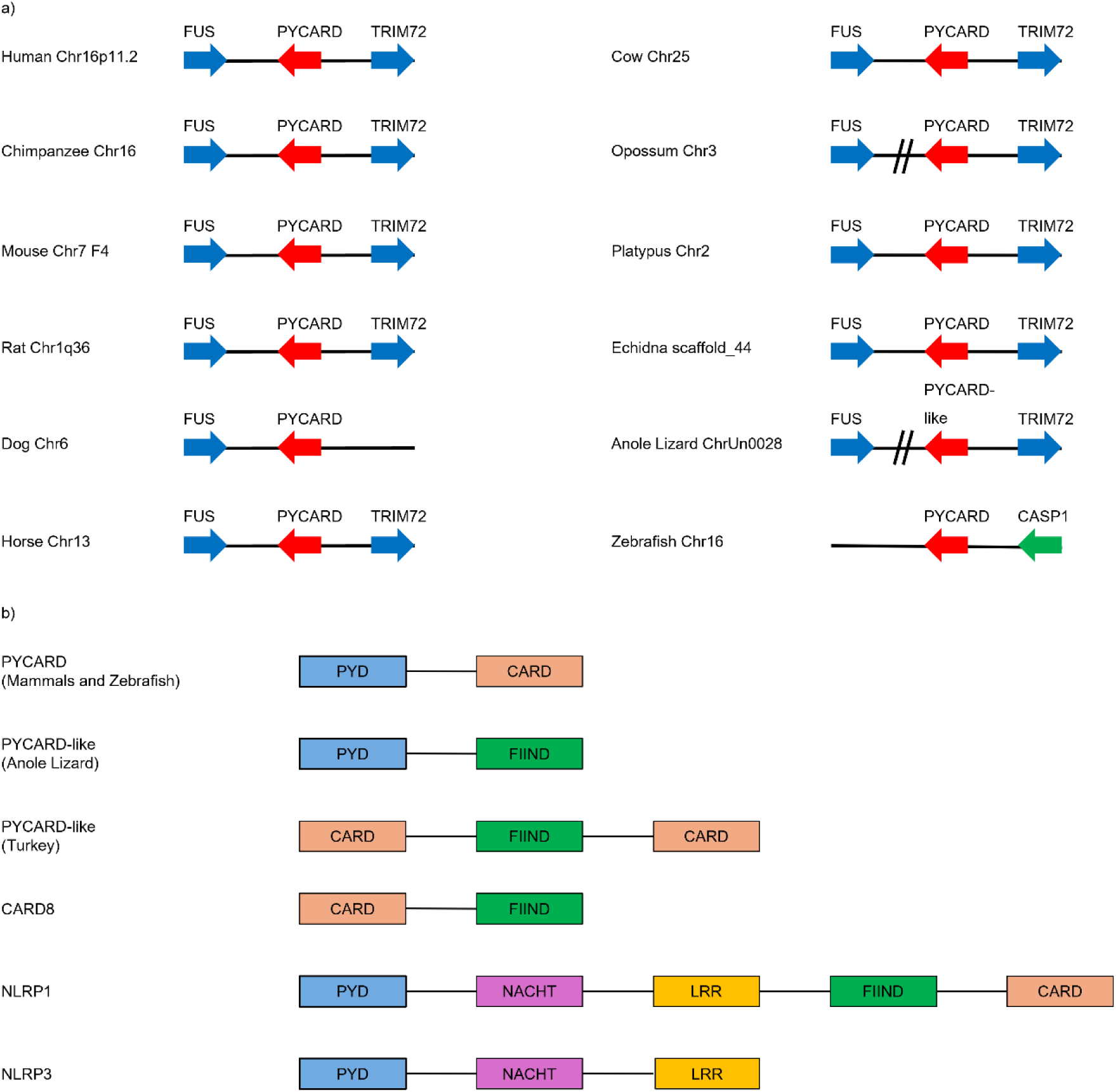
Analysis of *Pycard* and related genes and proteins. a) *Pycard* and flanking genes in multiple species. *Fus* and *Trim72* flank *Pycard* in each mammalian species with the exception of *Trim72* in dog. Anole *Pycard-like* is flanked by *Fus* and *Trim72*. Zebrafish *Pycard* is flanked by *Casp-a*. Red arrows show the gene of interest, blue arrows are genes with conserved synteny and green arrows are species specific. The double lines indicate these genes are on the same chromosome but not flanking. b) Domain organisation comparison of PYCARD and PYD/CARD containing proteins.

**Fig 4.**
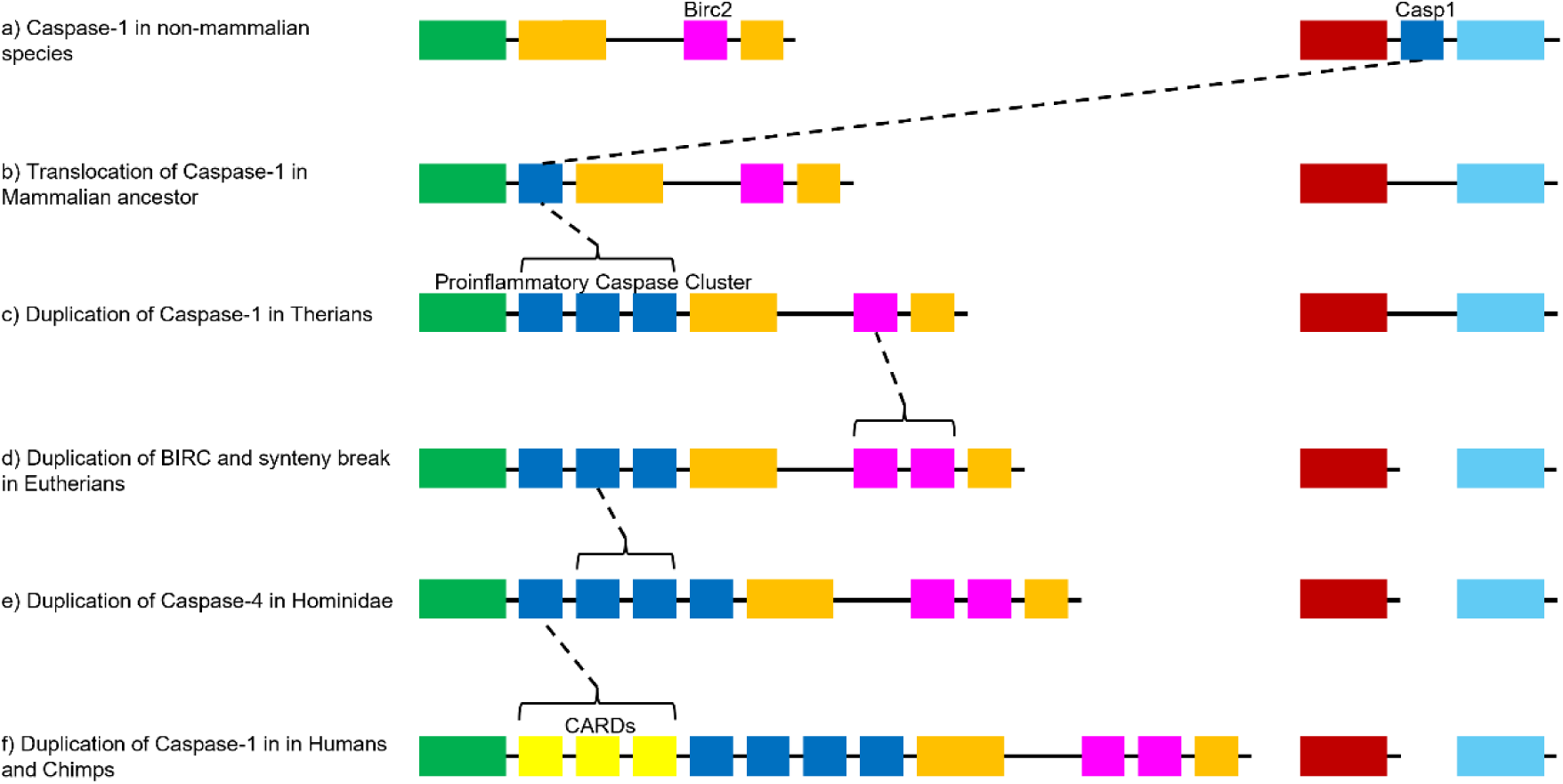
Evolution of the proinflammatory caspase chromosome region. In the mammalian ancestor, *Caspase-1* translocated to a different chromosomal region. In therians a duplication of *Caspase-1* resulted in *Caspase-11* and *Caspase-12*. In eutherians *Birc2* underwent duplication to generate *Birc3*. The genes that flanked *Caspase-1* in chicken separated with the upstream genes translocating to a different chromosome to the downstream genes. A duplication event in in *Hominidae* generated *Caspase-4* and *Caspase-5*. *Card16*, *Card17* and *Card18* identified in humans and chimpanzees are likely the result of *Caspase-1* duplication or duplication of one of the Cards themselves. A truncated Caspase-12 protein, the result of a premature stop codon, results in a shorter protein functioning as a negative regulator. Blue boxes represent caspases, yellow represent the Card genes, pink represent the Birc genes. Green boxes represent the gene cluster upstream of caspases in mammals and orange the downstream cluster. Brown boxes represent the gene cluster upstream of Caspase-1 in non-mammalian species while light blue represents the downstream gene cluster

Interestingly we were able to detect potential *Pycard* orthologs in turkey and in the anole lizard where the flanking genes were the same as those in mammals.

Analysis of the anole lizard PYCARD using the domain prediction site ProSITE revealed the lack of a CARD domain. However, the search identified a FIIND (Function to find) domain (a domain previously only found in NLRP1 and Cardinal) (Fig 3b). The automated NCBI annotation program predicted this gene to be *NLRP1* based on the presence of both of these domains but we suggest it is not *NLRP1* as the FIIND domain is located between the CARD and LRR in NLRP1 whereas here it is adjacent to PYD.

To help examine the conservation of *Pycard* in vertebrates, we identified a potential ortholog in the zebrafish genome but synteny analysis revealed that its flanking genes are different. Interestingly, one of the genes directly flanking *Pycard* is *Caspase-a*, the zebrafish ortholog of *Caspase-1* (Fig 3a).

Degenerate primers were designed to amplify *Pycard* in chicken. Initial designs had one primer in PYD and the other in CARD. These were unable to amplify *Pycard* (data not shown). The next set of primers was designed for the pyrin domain of *Pycard* which has a different sequence to PYD found in other genes and these also failed to amplify the chicken gene. Primers designed for CARD in *Pycard* were likewise unsuccessful. Both the CARD and PYD primer sets were effective with platypus DNA templates. We then looked for the flanking genes, *Trim72* and *FUS* which appeared to be absent in chicken. The *Trim72* primers were able to confirm the gene in human but not in platypus or chicken. The *Fus* primers were able to amplify the gene in both platypus and chicken.

Mammalian and zebrafish PYCARD contain a CARD and PYRIN domain. These domains are found in proteins involved in apoptosis and inflammation and mediate the formation of larger complexes. The PYCARD-like proteins found in anole and turkey had a domain replaced with the FIIND domain. Previously FIIND has only been identified in two proteins, NLRP1 and CARD8 (Cardinal)

### Gasdermin conservation in eutherian mammals

Gasdermin D (GSDMD) has recently been implicated in pyroptosis, an inflammatory caspase dependant cell death pathway (Fernandes-Alnemri et al. 2007; Liu et al. 2016; Liu et al. 2024). Cleavage of Gasdermin D leads to the generation of an active N-terminal cleavage product that is then able to oligomerize and create pores in the membranes of cells leading to cell death (Liu et al. 2016). *GsdmD* could be identified in all therians examined. However, in chicken and anole lizard the closest orthologous match appears to be Gasdermin A (GSDMA). This result was supported in both species by synteny (not shown). In monotremes, a gene resembling *GsdmD* was found on chr4 for platypus and scaffold_29 in echidna. Reciprocal BLAST against the human genome assembly suggests the genes are most likely *GsdmA,* however analysis of the flanking genes shows conserved synteny with the region containing *GsdmD* in therians (Online Resource 8). Based on these data we believe this gene is most likely *GsdmD* however further work is required to better confirm these results.

### Rearrangement and relocation of Caspase-1 gene cluster

Facilitating Caspase-1 maturation and activation is the key function of the inflammasome. Caspase-1 maturation leads to the processing of IL-1β and IL-18 which are essential effectors of the inflammatory response. *Caspase-1* orthologs were readily identified in the platypus and echidna genome assemblies. Reciprocal BLAST of the putative echidna *Caspase-1* shows highest similarity to human *Caspase-4* (59.46%) compared to *Caspase-1* (52.47%) and *Caspase-5* (56.13%) compared to the platypus *Caspase-1* which show highest similarities to *Caspase-1* (52.23%). A phylogenetic tree was generated for the mammalian inflammatory caspases (Online Resource 9). This shows the platypus and echidna *Caspase-1* genes form their own cluster while dog *Caspase-1* which is the result of a fusion (Eckhart et al. 2008), is located between the eutherian *Caspase-1* and the other inflammatory caspases.

The genes flanking the Caspase-1 subfamily in mammals appear to be well conserved (Online Resource 10a). The most obvious variation within the region involves the genes present in the matrix metallopeptidase (MMP) cluster which appears to contain species specific duplications (Data not shown). The chicken genome assembly identifies a single ortholog of the mammalian *Caspase-1* gene. Surprisingly, comparison of the genes flanking mammalian *Caspase-1* with those in chicken revealed that the gene resides in entirely different locations in mammalian and reptile genomes (Online Resource 10a). Interestingly the mammalian *Caspase-1* flanking regions have conserved synteny in chicken albeit without *Caspase-1* (Online Resource 10b). Indeed, the genes flanking chicken *Caspase-1* can be found in the same order in mammals except that genes 5’ to chicken *Caspase-1* are found grouped on a separate chromosome to those that were located 3’ (Data not shown). In monotremes, however, the chicken *Caspase-1* flanking genes are still located on the same chromosome suggesting the splitting of this region occurred after monotreme divergence. These observations suggest that a transposition event has occurred early in mammalian evolution that resulted in relocation of *Caspase-1* and later, the chromosomal rearrangement of the region (Online Resource 10).

### Complex rearrangements of the Caspase-1 cluster in opossum

Of all mammals investigated opossum underwent the most dynamic changes to the Caspase-1 region. These include a large expansion of the Caspase-1 subfamily through multiple *Caspase-1* and *Caspase-12* duplications (Eckhart et al. 2008) and this report identifies additional rearrangements. *Actin-like 6a* (*Actl6a*) has been inserted directly downstream of the third *Caspase-1* along with a large inversion which includes all genes from *Platelet derived growth factor D* (*Pdgfd*) through to *Contactin-5* (*Cntn5*) including *Birc2*. The full size of the inversion and insertion cannot be determined due to the large number of hypothetical genes between *Actl6a* and *Cntn5*. We then investigated whether similar events have occurred in other species and found that cow and dog genomes also feature changes to more distant parts of the region (data not shown). The genes upstream of *Gria4* are on a different part of chromosome 15 in cow (which is the same chromosome that carries the Caspase-1 cluster) and genes downstream of, and including, *Progesterone receptor* (*Pgr*) are on chromosome 21 in dog.

### Duplication of Birc2 generated Birc3 in mammals

The IAPs are inhibitors of apoptosis that function by interacting with certain apoptotic caspases and TNF receptor-associated factors (TRAFs) to prevent apoptosis (Burke et al. 2010; Mace et al. 2010). The IAPs have 3 baculoviral IAP repeat (BIR) domains and a RING finger domain. Of interest in this case are *Birc2* and *Birc3*, IAP members found downstream of the proinflammatory caspases (chromosome 11q22 in human). *Birc2* is present in chicken and all mammals investigated including the monotremes where it was found on chr20 in platypus and scaffold_46 in echidna (Online Resource 10c).

*Birc3* however, is only present after marsupial divergence and, given the high similarity to *Birc2* at the protein level, is predicted to be the result of duplication. Interestingly, *Birc2* is present in zebrafish with another IAP protein which is most similar to *Birc2* but also has homology with *Birc3*.

## DISCUSSION

### Evolution of the NLRP3 inflammasome

The immune response to external challenges is a vital characteristic of multicellular organisms. Many of the genes involved are highly conserved while others display extensive change. In this study we aimed to shed light on the evolution of the NLRP3 inflammasome and in particular the region containing *Caspase-1*.

This analysis shows that genes of the NLRP3 inflammasome pathway components are mostly conserved in vertebrates (*IL-1β*, *NEK7*, *PANX1*, *P2X7*, *TLR2* and *TLR4*, *MyD88*, the Nf-κB family, *Card9, MALT1* and *BCL10*). The conservation of these genes is likely due to their function in fundamental cellular processes other than the immune response. For example, Nf-κB is involved in apoptosis (Ryan et al. 2000) and the neuronal function (Freudenthal et al. 2005; Levenson 2004; Meffert et al. 2003).

Our analysis suggests that *Pycard* is missing in chicken and zebrafinch but we found potential orthologs in the anole lizard and turkey. It is possible that these could be lineage specific deletions, or it may be that the genes have diverged beyond recognition possibly to allow it to interact with Caspase-1 in different ways from what we know in mammals.

Based on phylogenetic analyses, *NLRP3* and *NLRP12* diverged from the same ancestral gene with a similar event giving rise to *NLRP1* and *NLRP10*. The majority of monotreme *NLRP-like* genes showed highest similarity to either *NLRP3* or *NLRP12*.

While platypus has one copy of *NLRP3* that cluster with the eutherian *NLRP3* a second potential *NLRP3* homologue clusters with the Opossum *NLRP3* which is more similar to eutherian *NLRP14*. Whether this gene cluster is the result of a divergence between the ancestral *NLRP14* giving rise to a unique NLRP family that has been lost in eutherians is unclear at this stage. The results suggest that an expansion of the NLRP family occurred prior to monotreme divergence as seen by the platypus NLRP repertoire (*NLRP3*, *NLRP6*, *NLRP10*, and *NLRP12*) with *NLRP6* lost in opossum and *NLRP1* arising after therian divergence.

### Pyroptosis outside of eutherians

It has been shown that mutating *GsdmD* at key residues prevented the proteins from oligomerizing and blocked pyroptosis (Liu et al. 2016). This implies GsdmD is essential for pyroptosis and the absence of this gene in chicken raises the question of whether pyroptosis occurs in this species. The Gasdermin family member GsdmA is present in chicken and is capable of inducing autophagy through its N-terminal while the C-terminal acts to inhibit it, similar to the behaviour of the GsdmD cleavage products (Shi et al. 2015). The putative *GsdmD* found in monotremes suggests the pathway may be functional, however, the absence of *Caspase-4/-11* orthologs suggests the noncanonical pyroptosis pathway may be slightly different to what has been observed in human and mouse (Kayagaki et al. 2015).

### Conservation of the NLRP3 inflammasome fungal response pathways in mammals

The Dectin family is located next to NKC genes and, like the NKC genes, are c-type lectins. The NKC has undergone massive expansions in monotremes (Warren et al. 2008; Zhou et al. 2021). However, the Dectin family has not undergone an expansion with eleven genes identified in monotremes compared to 13 in humans. The Dectin-1 subfamily appears to be mostly conserved although it was not possible to conclusively determine the identity of *Clec-like1*, *-2* and *-3* in monotremes. More interesting is the chromosomal relocation of the Detin-2 subfamily in monotremes. Of the six Dectin-2 subfamily members only three appear to be present in monotremes however the BLAST results and phylogenetic tree were unable to clearly determine their identities although it does appear likely that *DCAR* (*Clec4b*) and *BDCA-2* (*Clec4c*) is absent in monotremes. While Opossum appears to have lost most of the Dectin family genes the presence of *Mincle* (*Clec4e*) suggests the NLRP3 inflammasome response to fungal threats is likely conserved.

The duplications of *Syk* in the monotremes is interesting as the genes appear to have ORFs and should therefore be expressed and have functions. However, due to the locations of platypus *Syk-like2* and *Syk-like3* on small contigs we cannot rule out the possibility that these are a single gene.

Together this suggests that all three lineages of mammal can induce the NLR3 inflammasome in response to fungal threat.

### Evolution of the mammalian proinflammatory Caspase chromosome region

The *Caenorhabditis elegans* cell death gene *ced3* is thought to be the ortholog of the caspase family and it functions, like most caspase family members, in programmed cell death (PCD) (Yuan et al. 1993). Caspase-1, however, is involved predominantly in promoting inflammation although it can also cleave apoptotic targets under specific conditions suggesting that Caspase-1 has acquired its inflammatory function over evolutionary time (Miura et al. 1993; Wang et al. 1998). Based on current data on Caspase-1 homologs in zebrafish and gilt-head seabream and Xenopus, it appears that this change in function has occurred after the divergence of *Actinopterygii* (Lopez-Castejon et al. 2008; Masumoto et al. 2003). It is interesting that Caspase-a has an essential role during zebrafish development and requires zebrafish PYCARD for activation (Masumoto et al. 2003). Caspase-1, however, has no such role in mammalian development (Li et al. 1995). Furthermore, in place of the Caspase recruitment domain (CARD) found in the prodomain of the mammalian *Caspase-1*, the *Caspase-a* and *Caspase-b* prodomain contains a pyrin domain (PYD) and the interaction between Caspase-a and zebrafish PYCARD is through the PYD (Masumoto et al. 2003).

An amino acid change in the cysteine-active site of chicken Caspase-1 raises doubts as to its ability to cleave its substrates (IL-1β and IL-18) to their active states and for inactivating IL-33 (Johnson et al. 1998). Furthermore, chicken IL-1β is lacking the Caspase-1 consensus site necessary for activation (Gyorfy et al. 2003). The protein function was shown to be ineffective in comparison to an N-terminal truncated chicken IL-1β protein (removal of the prodomain) which showed 100 fold enhanced function in mammalian cells (Gyorfy et al. 2003). Interestingly a study of the IL-1 family in a variety of species, identified a lack of Caspase-1 cleavage sites (i.e. the necessary aspartic acid residue) in IL-1β of chicken, Xenopus and rainbow trout (*Oncorhynchus mykiss*) suggesting that Caspase-1-targeted cleavage of IL-1β may be limited to mammals (Huising et al. 2004). These data, together with the apparent absence of PYCARD, would support the possibility that the NLRP3 inflammasome is inactive in the chicken. However, several recent studies show that NLRP3 pathway components are upregulated by different stimuli (Chen et al. 2020; Huang et al. 2021; Karaffová et al. 2020). In comparison, all the mammals examined in our study have the cysteine-active site in Caspase-1 as well as the Caspase-1 cleavage site in IL-1β.

It is possible that the change from pro-apoptotic to pro-inflammatory function occurred as a stepwise series of events as evidence in mammals shows that active Caspase-1 is involved in the secretion of Il-1α and fibroblast growth factor 2 (FGF-2) (Keller et al. 2008). This is interesting since the latter are not normally substrates of Caspase-1. The Caspase-1 substrates IL-1β, IL-18 and IL-33 lack signal peptides to direct secretion from the cells, a feature also absent in IL-1α and FGF-2 (Keller et al. 2008; Qu et al. 2007). Furthermore, it is likely that additional specialization occurred leading to the proinflammatory functions found in mammals.

Relocation followed by massive reorganization of *Caspase-1* in mammals might have facilitated changes in its regulation and function. Following mammalian divergence *Caspase-1* relocated to a different chromosomal region in mammals, causing the flanking genes to separate and move to different chromosomes (Fig 4, Online Resource 10). No further change is observed in the region until after monotreme divergence where an event, likely the duplication of *Caspase-1*, results in the generation of *Caspase-11* and *Caspase-12* (Eckhart et al. 2008). Gene duplication events are known to be key facilitators of neofunctionalization and subfunctionalization (Conant and Wolfe 2008). The similarities between *Caspase-1* and *Caspase-11* make this duplication a likely explanation for subfunctionalization. *Caspase-12* also shares sequence similarities to *Caspase-1* but does appear to be more divergent. Interestingly, Caspase-12 in mouse has been shown to act as a negative regulator of the inflammasome by suppressing Caspase-1 binding (Saleh et al. 2006). The region in opossum has previously been shown to feature a large number of duplications (Eckhart et al. 2008), and we identified an insertion event, and a large inversion of the region adjacent to the third *Caspase-1*. It is possible that this duplication was due to a functional requirement by marsupials to provide a more robust immune response due to their young developing externally in the pouch. Further expansion only occurred after the divergence of the great apes, where a duplication of *Caspase-4* led to the generation of *Caspase-5* (Eckhart et al. 2008).

Some species show species-specific variations in the number of proinflammatory caspase members that are present (Table I). An interesting example is in canines and other carnivores where a fusion event between *Caspase-1* and *Caspase-4* created a gene which encodes the active regions of Caspase-4 with the Caspase-1 prodomain (Eckhart et al. 2008). The curious feature of this gene fusion is that the product still functions as Caspase-1(Eckhart et al. 2008). Studies in mice have shown Caspase-11 to be essential for Caspase-1-induced apoptosis although Caspase-11 can independently induce apoptosis (Wang et al. 1998). Caspase-4 is also required for inflammasome activation in response to certain gram negative bacteria using either a TLR4 dependent (Rathinam et al. 2012) or independent pathway (Kayagaki et al. 2011; Kayagaki et al. 2013; Sollberger et al. 2012). Data for Caspase-5 suggest a role in Caspase-1 processing in the NLRP1 inflammasome but no apoptotic roles. In the case of the NLRP1 inflammasome, Caspase-5 interacts with the complex through the adaptor Cardinal and thus processing of both Caspase-1 and Caspase-5 occurs (Martinon et al. 2002). Interestingly the region containing the Caspase-1 family in humans is known to be a site which frequently undergoes rearrangements in various cancers, suggesting the large genetic variation may be due, at least in part, to chromosomal instability (Cerretti et al. 1992; Du et al. 2010).

### Birc genes and the role of apoptosis inhibitors in promoting inflammation

While inhibition of apoptosis can be achieved in several ways, the IAPs are key cell death inhibitors and include BIRC2 and BIRC3. The Birc genes interact with Caspase-3,-7 and-9 though it has been suggested these interactions only weakly inhibit apoptosis (Eckelman and Salvesen 2006). More recent data suggest that BIRC2 may be able to prevent pro-Caspase-3 maturation through interaction with the Caspase-9 apoptosome (Burke et al. 2010). BIRC2 is also able to interact with TNFR-associated adaptors such as TRAF2, blocking the Caspase-8 apoptotic pathway and promoting pro-survival pathways (Mace et al. 2010). Conversely evidence suggests that BIRC2 contains cleavage recognition sites for Caspase-8 and that this is essential for TNF-related apoptosis-inducing ligand (TRAIL) mediated apoptosis (Guicciardi et al. 2011). Interestingly evidence suggests that Caspase-8 can also process IL-1β for pathways involving TLR3/4 and Dectin-1 suggesting that the Birc proteins may be regulating this inflammatory pathway (Gringhuis et al. 2012; Maelfait et al. 2008).

The Birc genes are located adjacent to the region containing the Caspase-1 subfamily cluster and were there before *Caspase-1* relocated to this region, evidenced by their presence in the chicken genome in the same region as they are found in mammals. Prior to marsupial divergence, only *Birc2* can be found in this region. Following marsupial divergence however, a *Birc2* duplication and divergence resulted in *Birc3*. These two genes have been shown to play an important role in signalling through the NOD1 and NOD2 pattern recognition receptors, both of which are NLR family members whose knockouts result in cytokine production attenuation (Bertrand et al. 2009). Given their role in the regulation of Caspase-8, it is possible they may also regulate the non-canonical Caspase-8 inflammasome pathways. During pyroptosis BIRC2 is degraded in a Caspase-1 dependant process which is currently not understood (Wickliffe et al. 2008).

## CONCLUSION

The fundamental machinery involved in NLRP3 inflammasome activation has been largely conserved throughout amniote evolution. The exceptions are the proinflammatory caspases which have undergone frequent lineage specific changes. While all major mammalian lineages have the genes necessary for NLRP3 inflammasome activation in response to fungal threats, the absence of multiple Dectin family members in opossum suggest a potential loss in redundancy. Whether the duplication of *Syk* in monotremes has led to new functions in fungal defence or added redundancy requires further analysis.

This analysis shows that Pycard is conserved in mammals however, we observed differences in non-mammalian species. Chicken and zebrafinch appear to lack Pycard, while green anole lizard and turkey have potential orthologues that contain FIIND instead of one of the ‘classical’ Pycard domains (PYD and CARD respectively). The potential effect of these changes on the inflammasome pathway are unclear however it indicates differences in the inflammasome complex in these species.

Caspase-1 is a key component of the NLRP3 complex and has undergone multiple duplication events during mammalian evolution resulting in an expansion of the inflammatory caspases subfamily as well as lineage specific duplications. Our data suggest that *Caspase-1* was translocated into a new genomic environment in early mammals, prior its expansion. We hypothesise that this relocation as well as chromosome instability in this region facilitated the amplification, functional and regulatory diversification of the *Caspase-1* subfamily.

Overall, we found that monotremes have well conserved canonical and fungal NLRP3 pathways, however the absence of several inflammatory caspases present in other mammals (especially Caspase-4) may reduce the number of noncanonical pathways available to monotremes and may result in differences in the inflammatory response.

## Supporting information

Online Resource 2

Online Resources 1, 3, 5

Online Resources 4, 6-10

## ACKNOWLEDGEMENTS

The authors would like to thank Emeritus Professor Jeremy Timmis for his constructive comments on drafts of this manuscript.

## STATEMENTS AND DECLARATIONS

### Author Contributions

DS: Conceptualisation, scientific planning, data curation, interpretation of data, validation, writing and figures, review and editing.

TD: Conceptualisation, scientific planning, interpretation of data, review and editing. FG: Conceptualisation, scientific planning, interpretation of data, review and editing.

