## Supplementary material for "Large-Scale Restructuring of the Caspase-1 Gene Cluster Region in Mammals": Online Resource 2

Journal of Molecular Evolution

Online Resource 2

a) Platypus gene sequences

>PYCARD_like_1

MVPVPRKLHFVDRHRKRLIDRVTGVDSMLDSLHGQVLNEEQYQNVRAEKTNPDKMRKLFS

FNPSWDQGCRDRLYQALRETHPHLVEELESYLVCKMGMKTVSST

>PYCARD

MGSARDHILTALENLTSDEFKKFKTKLLSCPLREGFGRIPRGPLMNMDVIDLTDKIVTTY

MENYGLELTTAILRDINQAEAAALQKAAGAAAPRVGPSKIALGEAEQQHFMDKHRQALIN

RVTSVDALLDALYGKVLSEQQYQEVRAEKPSANQMRKLFSFSVAWNRSCKDQLFQALKAT

HPFLIKGSDGSDQVPRSKRAKDHYTKRFEEYNIRVAFPGHNELPSGEGQPHFVDNYLICK

MGMEMVSSSWDHLIILYLPQRLEQCSAHNSKLAEGKECVCLLYRTLPSTSYGALPTVNAQ

QIRLTAFSPMEDRSNRKKEARENGQVVRQNQNGILFLKWLQSALAYNTTEHSCLSRFRCL

KWNGMDFVA

>TRIM72

MSAAPALMQGMYQELSCPLCLKLFESPVTAECGHSFCRSCLARLPQDPQAGGTPCPSCQA

PTRPEGLSTNQQLARLVESLAQVPQGHCEEHLDPLSVYCEQDRVLICGVCASLGKHRGHS

VVTASEAHQRMKKQLPQQRLQLQEACMRKEKSVALLDRQLAEVEETVRQFQKAVGEQLGV

MRAFLSALEKTLGQEASRVTGEAGTALQGERRGLASYLEQLRQMEKVLDEVTDQPQTEFL

RKYCLVTSRLQKILAESPPAARLDIQLPIISDDFKFQVWRKMFRALMPALEELTFDPATA

HPSLVVSPSGLRVECVEQKAPPPGDDPQQFDKVMGVVSHQLLSEGEHYWEVDVGDKPRWG

LGLISAEAGRRGKLHPIPSQGFWLLGFRDGKVYEAHVESKEPKVLKVEGRPTRIGIYLSF

QDGVLSFHDASDPDNLAPLFSFRERLPGPVYPFFDVCWHDKGKNAQPLVLVPASSVLSFF

SPLTPSAKVLLPHRPESSSGMKRGAEGWGRSLRGSEEEEHLGGARTGQRPNLCPECGRGF

SQRSDLVKHLRTHTGEKPYPCPACQRRFSRGSDLVKHQRAHRRAALRLCGLRPPLQPELS

LPGPSAAPPEREALFLQPQRPLLWPQVHPGQALADAARGRALSPTGPPVPCTTWSIYRVS

TVKEQTEGEAAEDGVDDQELAFNQQLVSFCCVNDNWYYLSTYLEFT

>FUS

XYSTQPTPQGYGSSGGYGSSQGSQSSYGQQSSYPGYGQQSASSSGSGSYGSSSQSSGYGP

PQSGGYGQQSGYGGQQQQSSYGQQSSYNPPQGYGQQSQYNSSSGGGSGGGGSGGGNYGQ

DQSSMSGGGGGGGYGNQDQSGGGGGYGGGQQDRGGRGRGGGGGYNRSSGGYEPRGRGGGR

GGRGGMGGSDRGGFNKFGGPRDQGSRHDSEQDNSDNNTIFVQGLGENVTIESVADYFKQI

GIIKTNKKTGQPMINLYTDRETGKLKGEATVSFDDPPSAKAAIDWFDGKEFSGNHIKVSF

ATRRADFNRGGGNGRGGRGRGGPMGRGGYGGGGGGGSRGGFPSGGGGGGGQQRAGDWKC

PNPTCENMNFSWRNECNQCKAPKPDGPGGGPGGSHMGGNFGEERRGGRGGYDRGGYRGRG

GDRGGFRGRGGGDRGGFGPGKMDSRGRVLGGEARDP

>NLRP-like_1

MEVSLTELHFVDQPSLREQLIQRVTSVDQVLDKLFGSVLSQEQYQLVRAESTNPNKMRTL

FSFSPSWTQNCKDLLLKALREIHGPLLKELERSPEMQSPQDGAAIRKRYRARMEEKFSLL

RERNARPGESVVLQHIFTQLLCLPEHPQREQKEHELLAVGWDHARTMEEKGQFVEVNALF

GPDKIRFRRPKTIVLQGAAGIGKTMLARKIMLDWAKGNFFEDIFQFVFYLNCREMNQLSE

RSLSDLISVHLGDPRASFDEIMSQPEKLLFIVDGFDELKWSFEEQEYDLCYDWSEKRPVP

ILMSSLLRKILLPEAYLLITSRLTSLEALNNLLQHPYHVEILGFSEADRKEYFCRYFGDE

NQAMQAFDLVKDNETLFTMCFVPLVCWIVCTCLKQQLKCGKDLTQTSRTTTDLYVNYLAA

LFPATQDRPSQDPPILRRLCHLAAEGVWTQKILFDGDDLRKHGLDMSDVSPFLQLSIFQK

DIDCENSYSFIHLSFQEFFAAMFYALGAEEEAMDDSITDIGDVKKLLEENEKSQWKGFLT

LTVRFLFGLLNEARVKDLEKKFNCKISQEIKKELLQWSKGHCEILDLETLFDCLYEIQEK

EFIRTVMNRFQRIHLEIDTRMKLLVSSFCIKHCQTLQSIRIGGGDLDCEPIIENDLDSRV

KYGMEWGETGVWEHCTEAWVGQRVGYSPVPQQQQQKWRFPMESPLKKAQLPSELAGKLPN

DAHENDYFTSRVKQAWKDLFSGLRTNQDLTELTFSKLFIESGMETLCEELRHPQWKIRTI

RLDSCDLTAADCQGLSSAFASNQSLKYLDLIFDDPENTGVKWLFEALRHTDCSLEALGLQ

RCHLTVASCKDLSSALASNQKLKRLYLTNCDLNARAVKSLCEGLRHPNCSLEILGLQSCG

LRDACYEDLSSILTGNKKLTRLDLFGNLLSESVKQRLQNDSKHANFNYERKSAQPQVGGP

KCPLNSGVPWALTLPGASFPSFSPMVGTMAVITLALPAISLSLLTSLVSFYILTSTLHSP

AAKLLLCLVLTCLAIDS

>NLRP-like_2

MAPAPRDKLLLSLKELREEELRTFKFKLLGIPLQDGYFHVPRGEVDGLKPVELADLLILY

YGEKYAVMVVLEVLKAMHMNGVVETLGQDTWEDSRETYRERMKKSLERAEERHGHLRDRV

DLRHRFTPLCLVTNSLPGAQEEEGPAPVAEGNGSSGIGEKEVHSIGLETLLDPEEEGTDS

PRTVVLHGEAGIGKTTLAKKLMLDWASGDFYPDTFDYTFLVSCREINLMAEKSLAELICH

CYGDGHAPVTEVLKRPERLLFIIDGLDELKYSLEEEGEEPGSELRDKQPVQTLLSSLVRK

KLLPESSLLITTRPRALEKLQPLLEDPRYVEILGFSEVEREEYFFKFFTDENKAKKAFNF

VQRNENLSSLCSIPLVCWFSCTCLKQQMVRGEPLTHTSKTITDVYMSYISIFLQPDRARA

KQPAYPTLKRLCSLAADGIRNRKILFQEDDLRQHSLDQADIAAFLSKANFRQESKGETFY

SFIHYSFQEFFSALYYMLEGDGNESNRDVNALLEKQEQTRSDGSALTLRFLFDFLSQETV

STLEIKFSYKIPPKLQKELKCIKARVKSLLSPIQKGPFELNISLNKIPKENNEQDSRDNF

QTMKQSKVSSKEGRMSVDMDKGFPQGIQGWIGDEK

>NLRP-like_3

MLFLRFDGLAHFELRLINQLIRIFCIPPAELGFVLFILRCLSAHYVPGAVLSTGVEDLLD

EKLNNFKFKLQDISLEEEFKRVPRGRLKPAEPVELADPLIRHYREDYAVKVSLEVLRAIN

QRQLAEELSQATGQVVVPIKVQSLKWSPREADDPDQGHNEGEDPAQRPQGLIQSSSEVMY

PPAVEYPSRIPQEDPHDVEDENSLLGANPEYLRLLLVKEHPGQLGSAGHKQMSATQVDAL

FDPEEEGRKPFRTVVLQGAAGIGKTTLATKILEDWAAGRLFHGRFDYVFYVSCREMNLVR

ERSVADLISHCCADKNGPVTDIVWLPKRLLFIVDGFDELRCSNERSRKDLCSDCKEKRST

DTLLASLMSKRLLPEASLLITTRPTTLEKMQALLEQPRWAEILGFAEAERKDYFYRYFKE

KKLARQAFGFLQGNDGLFTLCMVPIVCRIVCASLKRQMERGEALTQTSQTTTAGYLSYLF

GLLREDGTGPKAPRPLNLRRLCSLTAERIQNPKILLEEDDLRRHGLGSSGSTTFLKISIL

KKNHHPQKFYSFPHLTFQEFFAAVSYVQKTDFLLHVFYLLMRKNETRNNLATSSQFVKKL

VGQYGQFGSGYLTLTMRFLFGLLNEERPGEEPGLIEIGALATKLDHTVASFCVKNCPGVC

SVTLSSVYSAGGKGDDVNAVVEEGLSLGSQHESSHRLGKCFLPDAFCQNLSSALHTDPNL

LELNLSHIALGNAGLRVPSEGLMDQGCRLQKLT

>NLRP-like_4

MLLALLGLLLCGPRPSNVSLPDCGAAPGLGRPGDLLLAGVFALQTPPAPTPLRFQRPPQQ

PRPNKFSFSHYRMVQSFIFAVEEVNRRADVLPNLTLGFSIRDSGDSEFQALRATMGFLTG

RGQPAPNYRCGTGPPVAAVIGEDRSALTIPMATWLGLYKFPQVSYGATVAVLSDKTRFPS

FLRTLPPDSFRARGLARLVSHLGWTWVGLLAQDDDYGQQGSRALGTALAAAGVCVEFLLE

IPSSRAEERLRHLGRAVGRASARALLVFSGDERFLLLAGQLAGAGGPGKVWVGSDVWDPA

LLAPGGPLARTLGGALVFSSHRGQIPGFREFLGQLHPDRRPEDPFVITFWEEMFACRWSN

RTEGPRGGLGEAPWCTGEESLKGRDHPFLDMGDLSVTYSCYNAVHSVARALHALASCEDG

AGPFGGGTCARIRDFQPWQLLHYLWKVRFRNSHGEEVFFDALGDPPALYDIMNWQETPGG

TFHFQEVGKFDSTTLPGQEMVLNDSSILWLGMEKQVPRSVCSESCPPGTRQTPHKGKPPC

CFDCTPCPEGQMADKTDSLECRMCPGDDFWPNKRKDQCVKKAVEFLEFGEPLGAALASLG

ATLALATSAILALFIRHRETPIVRANGRALSYVLLTSLLLCDLCSLLFLGRPGALTCLLR

QVAFGLTFTLGVSCVLAKTIMVVIAFRATVPGSPLRRWLGPGLPGAVVLACSLAQGVICA

TWLAAAPSFPERDTRSSPGKVLLVCNEGSPVAFWCVLGYLWLLALVSFVLAFLARRLPDG

FNEAKFITFSMVVFVSVWGAFLPAHLSTRGRATVAVEVFSILCSSSGLLGCIFFPKCYIL

LVRPDRNTREFLMGRETAGPRYLTCQMGMKTVSLMSDNLMTLYLPQRLEQCYAHNYEPIV

GQGLSICCQIVLSERLVQCSALGLCSIRLATLLLRSEFSVSEIQVMHLGILCLSDPAVLS

GHKKQKVRLDRIDILPLMRLIALPDLGADEAVLRVEVQEGIFHQPAAGLSEERDGPETGK

SPPSNTPSGSSLGTLGIHHLGSFWPEAEMTCSIRSRLAEYLEELRDPELRKFKFHLEDLA

PAAGWAPIPWGRTEKADSLDLAHLLVAHVGERGAWELAVHVFERIHRKDLWERARAEAEV

RDSSGSVLGRHWEQRRRAEHLVETHEAQATRDPRDVYREHIRRKFRFFEDRNARPGECVN

LSQRYTQLLLMERHPTPRDAAPVSPTPQREDAGAPERQPGPLQLETLLEPDASWPEPPRT

VVLRGAAGIGKSMLARKIMLDWADGLLYRARFDYLFYVSCREMSRVGRGSLAGLISGRWP

RREAPLADILRRPERLLFVIDGFDELGRTPRRPPPGRRAGGWKAKMPVAGLLGGLLGKDL

LPEASLLITTRPAALARLQPLLEQPRHAEILGFSRAGRRDYFRKFFGDGRQASRALGLVR

DVEALSAVCFVPLVCWIVCTCLKQEMERGEPPGQPSKTTTAVYVFYLLSLLRPDPGGSGP

RGRPVLAPLCSLAAHGVWARKVLFDEGDLSRHGLAGSDVSAFLNLSVFQKDIHCQRLYSF

VHRSFQEFFAALFYLTDEGGREGGRGSRHSVTRLLEHYGRSETGYLALTVRFLFGLLNGE

NKSYLERQLGCPLSPTVKGELLAWIEAGARIGGRTLERGALDLFSCLYETQEEGFIRRAL

DPFQVIVACDLSTKMDHVISAFCVGNCRNASVVHLGSEEFSSEEAAEGQGPVGTEGTHLG

DQPLSPVEKCWLPDTYCEHLSSALRTNQNLAELVLDRNALGNRGVKLLCQGLGHANCKLQ

NLGLKKCRFSSAACQDISSALSANQNLVMMDLSNNALEDVGVKLLCAGLRHPKCRLQSLQ
LKKCYFSWAACEDLSSVLSTNPHLMELDLTGNALGDAGVQLLCVGMKQTSCRLKTLWLKI
CHLTRASCMELASVLSVDCSLTELDLSLNDLEDAGVRLLCEGLGQPKCKLQKLRLGICRL
TSAGCGAVSTALGANGHLKELDLSFNDLGDVGAQQLCDGLNHPNCKLQKLWLDSCCLTAV
ACESLSSVLAKNQTLTKLYLTNNALGDAGVGLLCERLKQPTCKLQTLWLFGMELKAETQN
ALAALRRTKPHLDIGS

>NLRP-like_5
MEFTERLRCAEHRTERLRESDPTESVDPSPALRELAVSPSRPSIRGGQGVGLLIVVLSSP
KRSVRRSAHTEHFVDRHRGQLIGRVTSVKPLLDLLHGKVLSEEQYQTVLAGATSFDQMRK
LFTYSLSWDRDCKNLLYQAMRKIHPQLVAELDGSLRFSGPRVPQLEKMGMKTRLGETDCT
VSVDTFPVHNELTVFETHSDVLVCLYPNCRQPWEGVSLAEDYREKYLKYVRWKFRYLEER
NARLGEKVALEVRYTPLLLVEEHRSLAQRQHELLALGRQPTSWVSRRVHVEALFDPDAEG
LEPPLTVVLQGSAGIGKTVLARKVMLDWAAGTLYPGRFDFAFYVHCRELNLERKRSAVQL
IQQCCDDDGVPLSEICRRSDRLLFLVDGFDELGWSSGGWPGAADDDDDLFVDWKDKRPVG
SLLAHLIRKRLFPKASLLITTRPAAAESLRPLLRWPRRAEILGFSEAERAEYFHHYFLDS
GRANRALAFIQENDVLFTMCFVPLICWIVCTGLRQQMDRREDLAQASKTTTAVYLSFLSS
LLRPARHLPSRVPPAHLRGLCSLAADGILSQRILFRKADLRKHGLPEDGLSAFLHVDVFE
KDVDCETLFSFVHLTFQEFFAALFYLLGPDETGPGPLPDARTLLQHYRLCDTGFLTLTVR
FFFGLLNRERVADLKATMDCVVRPEARRALVEWIDTSARKEALPGPTLQWLYCFYEIQEV
DFVERAMGSFRKIDIDVRTRMDQIVVAFCLRNSRNLCSIELMRFFNVAEADAAAAAAERT
PEVAADRRPQDWLQDTLCESLSEASANNRGLTHLALRNKNLGRRGAELLSKGLSHPNCKL
KSLRLVNCLLTPDSCRDFSSILSTDRNLSELDLSSNALKDSGLSLLCEGLRHPSCKLQTL
CLVKCLAHYQVHEEGVRSDYLVSSSALGENCRYTSQNDFTAEGGPFWKGLQSCALTSACG
PALSSLLTTSANLTKLDLFNNALGDTGVSLICEGLKHPNCKLQALLLRHCALTLNCCPDL
SSVLSINRDLSELDLSYNSLEDAGVCLLCEGLRHPNCKLQTLRLQKCKLTSRSCSDLSQA
LGINRELRELILHDNVLGDAGISVIWKGIQQPSCGLKILRLGKTHFSEKMEEEMRTVQKM
KPELKIRYRPARC

>NLRP6-like

MEVPDSEARPREGVIRDLLWEALEDLKQDEFKKFGHKLVTINHEGRKNIPRGRLENVDLG

DMVDALVQFYDGDVALDVTRKVLKKMDMKAAATKLKEKRRRGDDPPGPSDFPGPGLEGRA

GAPNLKSQVTVAGKGLAPGLGETEGFAEEAARPVPAASTGDHILAGAREGEERNARSVED

QQALTKSPDLARGAAADEERGRRPLGGGGGGRTRSGPRRSDANTFNELFHKDEDGRRPGT

VVLRGPAGIGKTMTRPEDHVRLGRGQAVPRPVRLRLLRRLQGGGRGGGGALAGRPGAGPL

PRPAGAREGHAGRAGPAPLRRRRRTSCPPCRGGGRAVLRPLREGGPAVLLRSLLRSRTVL

PGASLLVTTRPAAPERLRACLRSPRCAEIWGFSDRDKKKYFFRFFKDEGQARRAYRFVKE

NETLLAMCSVPFVCWIVCSVLRRQTESGHDPARASKTTTAIYLLFVTSLLDGCDAEARAR

GRADLRRLCRLARQGVWEDRGRFGPEDLRRHSLTGSVIPTLFLRVLTVQRDLRSEVVYQF

IDQSFQDFFTALSYLLVDAGEAEGEGPGGARPLPDGAGTAAAAAAAAGPDDAERVLGCAG

ARAVRPAVLAWLEREGREAGRQGALPSPAEHLPPAEHVPPAEHVPPRHVLPSEHVRLAEA

VGDEDPEEEEEEEEEEQEQLELLYRLYESQDWELTRRGLAARGSVRVVGVGLGRLDQAVL

AFCLRSAPPGLALQLHRCAFAGAADQARKGHRSWAGKLSRGLASGGSHRSDKLKSPKRSS

LWPLCEALGDPQCKLKTLTLSHCRLSDAECRDLAEALASSRTLTDLELLGNKLGALGMRN

LCLGLSQPHCPVETIRLQQTSSREAYLELVNVLWVSPRLQALDLGHSTLDGLTVARLCHG

LRHPNCRIASLSLRQCSLLPNSWAEVAAIFASTTTLREVDLSGNPLGAAEIGTLCEGLRR

PDCKVNRLDLSSVDLCEEAVKGLVALSKAKPELVVVNPALSAPPEANSDTFSMLYPICKT

GMKTVSPTSHNPITLYLPQCLERCSAQSKCLRSSIIIIVVTGGGHWTIGEGARLDLSGVS

GSSVA

>NLRP14-like

MALSIRDLLTQTLEDLLENEFRKFKWKLCEIPLWASPRGQGGAASIPRGVLEKADPLTTT

ELILSYCGSSSALDVAARVLENIQQRELSNRLRERVPGKISQEMYRSEIRHKYERVKDYN

SLPGAWRSLEQHYVAPLIIRRCRPASEREPELLSKGPRHLELLRLCGEGGSDRVHLDHLL

DGPGGRRPLTVVLQGAAGIGKSYTAHKIMLAWASQRLYHDRFDWVFLFNCRELGVEPRPR

SLVDLVLSDCPALGPHVGQIFSSPQRLLFLLDGFDELQLPRSLEDDERGAEEEGEEEEGQ

EEEAAASAGWRRAGAAVRRRRPAAATVRLLLRQRLLPGCLVVVTTRPSALEQVEGCIRAD

VHLEVLGFLEPERQAFFTRFFGDAKRGREAYEAVQGNEALSTMCFVPLVCWIVCTVLHKQ

LEKGQGLDGLQTTTQVFLHFLSILLRFHRRRGARAPADSLLEQLGTLAVHGLMARKVIFD

QEDLEAHGLPAAAPPTIFLSAVLRQGVTVETVYSFGHVMLQEVFAAIFCFLPGRGARPEL
GLAHLLEAGLQAENGHLLQTIRFLFGLSHPQCQAMLRQLLPHRAALTPSPEGALLSWVQR
SAESPRAEDPRFLLELLHCLYEWHCDGLVGRVASELNIRFLLFPLKRSDCLALAYCLGCC
ASVSCLHLYSCGLDQGDIRLLLPALGKCQALHLGLSDIPSGLMQEIGRSFSPKQSVTSLL
LQGLGSNHSNSQKETVFKVSALWGSEPCSLRVNNVDKETTLEFCFWAVPAHRPREVTLQG
TQLCESSFRQICRLLRFSYSKLESLRISGNLLTKGCIPHLELLVRANTGLTHLDLSGNSL
GDEAVVLLCAHLGAAAGQLRNLRYVDPGEVPRLFGPCFPSSIDQWDLLVLCAEHCTKHLG
SGPREASLPLNSLVENGLTQECVPALSSLLPKLPALTCLKLGFNSLGDPGLLILAPALSD
PTCGLLKLDLEANGLTDACMPTLAQALAQNQTLEALILNGNKLSNLSIPHLDVIWREATH
LCRLELLFNQLPSTTRSRYSMAQAEPRDDVITTTPLYRETRGHGTRPRLTQRSSRPVPMP
NDGHRAVRVF

>MIS12 (NLRP1 flanking) no synteny chr17
MQKGAGKEKGLRDAEANGPPPEVLEEWDVLVEKGFGEQISSSRVKYRFFVPPPPVPPPAG
SPPGFRDRVRPSAPEERLATDERWTNTVIITVVITVIVAVASLPGPSSPSSPPPGGVPAA
EMSVNPMTYEAQFFGFTPQTCLLRIYVAFQDYLFEVMLTVERVILKRLEAAPGSGVSPVQ
IRKGTEKFLRFLKERFDGLFGTMETVLLQLVLRVPDHVLLPEDRSHARHPRGPEELARLR
EEADRLRGRYEAEVRAGRALLAELEEQRAARAELEKTLRWFDGLENAWREHGSGDPRESL
AFLIRSSDRLRAVVGDVERKGRRLHLS

>PGGHG
MGYGALDDPAVFTSPTLPSDPRFLATLTNSYLGTRVYRDILHVNGVYNGALGDAHRADVP
SPVNVRLEAPEGVEVSQSFTLDTRTGTFLHVVETPEFTATHRIYAHRALTHLLAFSVTVR
RSAPQAQPITVRLRSDFTPKSRDLDLHLGPDFQGARYLSGRTLSPEVAGGPQPTVHMLWT
PAPPALTLPEARREATWQFLVAVAEAEAEVRRLFEEGAALLRAGSLYPAHVEAWGALWGA
SGLDLDAPLPLQRAVRGCLYYLLSAVPSPAPGVRDPFHGISPGGLSNGSRGEDYWGHVFW
DQDLWMFPNILLLWPEAARAILQYRVRTLSGAQANARDQGYKGAKFPWESAATGHEVCPE
DIFGTREIHINGAVLLAFEQYYYSTRDLQLFKEEGGWDVVSAVAEFWCSRVIWSPEEQCY
HLRGAGARGKLGGCVGPRTSCLSSSFLTPASQINHPIRNDISRGVIPPDEYQTDVDNSVY
TNVVARNSLRFATSLGRDLGLAVPEEWLRVAENLKVPFDPKRRYHPEYDGYRLGDSVKQA

DVVLLGFPVPCAMDPDVRKNDLEIYEAATSPRGPAMTWSMFAVGWLELREPERAQQLLNK

CFANISEPFKIWTENSDGTETVNFLTGMGGFLQAVLFGYTGFRIIRDCLKFDPVCPTEVR

HGQVTGVSYLGNKLNFSFSEEEVTVEVTWAQSQAPALEAVLEPSGRRLALPQGQTVSFPT

TAGRIQRVSSYTT

>PSMD13

MKDVPGFLQQSQSAGPGQAAVWHRLEELYTKKQYKQGCSFKHPLMKTVSLTWDNLMTLFL

PQRLEQCSARSHTASREFSWFEFWQGMNVDFGGPDHSKRLVQCSAHSCGINSLFRSSILC

RTPALPKEMASSRRKQFSAASCPGSEGRRYRPQRLARSKRLTDAVTAPRTALRQLHQRVR

AQVRLPLRFLPRFLGPLALPYRRKSWWCCHYRVNPLSLVEIILHVVRQMTDPSVALTFLE

KTREKVKSSDEAVILCKTAIGALKLNIGDLPVTKVSRPTPGSHAFSPGTPEGREGGGAEV

PEVSAASLILPVLGREIGTVAPWPATTCVLPYPPPTCPRVETIEDVEEMLNGLPGVTSVH

SRFYDLSSKYYQTVGNHASYYKDALRFLGCVDVKELPVSEQQERAFTLGLAGLLAEGVYN

FGELLMHPVLESLRGTDRQWLIDTLFAFNSGNVEKFQALKAAWGQQPDLAANEALLLQKS

QLLCLMEMTFTRPANHRQLTFEEIAKSAKVTVNERSDPVAPGSGPRRTAPRSAGVQVELL

VMKALSVGLLKGSIDEVDRRVHMTWVQPRVLDLQQIKGMKERLESWCTDVKSMEMLVEHQ

AHDILT

>EIF3F

MAAAAAAGPAAPAAPAAPAAPPAPAAPAPEAAAVAVPGAAAAGSAGSAGPGPAPGVPSGP

ALGGPFPGGRVVRLHPVILASIVDSYERRNEGAARVIGTLLGTIDKHSVEVTNCFSVPHN

ESEDEVAVDMEFAKNMYELHKKVSPSELILGCTQDSAWHMRLEQCSAHGKRSTRTVVVIF

AVRLARPVNAADGSGALPGVGGGDRADPKRLRVPFRRYATGHDITEHSVLIHEYYSREAP

NPIHLTVDTSLQNSRMSIKAYISASMGVPGKTMGVMFTPLTVKYVYYDTERIGIDLIMKT

CFSPNRVIGLSSDLQQVGTASARIQDALSTVLQYAEDVLSGKVSADNTVGRFLMDLVNQV

PKITPEDFETMLNSNINDLLMVTYLANLTQSQIALNEKLLSL

>MYADM

MPITVTRTTITTTNMSSSGGNHTIVGSPRALTTPLGIVRLLQLLFTCIAFSLVAHIGGWF

GPMGDWCMFSWCFCFAMTLVILLVEMGGLQPRVPVSWRNFPITFACYAALFCLSASIIYP

VTFIKHHDKSEEKDCRIAATVFSILAFLAYTTEVCWTRARPGEVTGYMATVPGLLKVVET

FVACIIFVFISDTNSYERHGALKWCLAVYCIFFILSLAAILLCVGECTSWLPCSFHTFLS

GYTLLAVLAYATATVLWPLYQFSHRYGGQSRPNHCLREYGTLCFWDKLLVVAVLTAVNLL

AYLADLIHSARLIFVHV

>PRKCG

MSANSIALYSPKHLVQCSVHRSSVAEESSNPKFLALKPAAYGNGDGGTRAFCPHLPPPPP

GIPAPGTTFGDTSGLAIFVTHTDVAQDGDKRGIGKQGLQCQVCSFVVHRRCHEFVTFECP

GAGKGPQTDDPRNKHKFKLHSYSSPTFCDHCGSLLYGLVHQGMKCSCEYADVPLGDKEDV

LGDSDPGFGGKFPSRWKPSDPELRHPSPSCEMNVHRRCVRSVPSLCGVDHTERRGRLQLE

IQAPGGDELHITVGEARNLIPMDPNGLSDPYVKLKLIPDPRNLTKQKTRTVKATLNPVWN

ETFIFALKPGDLERRLSVEVWDWDRTSRNDFMGAMSFGVSELLKAPVDGWYKLLNQEEGE

YYNVPVADADNCGLLQKFEACNYPLELYE

>CASP1

MQPEEFHFHRIFLLLLPEKLKEDGCELTVSMPSEHKRKKAAKFLYRTLQNIPKTQLLKDR

WCLIIESLTHGMISGLLDDLLQMQVINQEEMDTVREEHHRPAEKTRALLNSVIPKGDLAS

QIFIDSLCKKNPFVAAKLGLSAVPQALQAPKTLTESHPDGSGEILKLCPSEEREKLQKEN

EGEIYPVLVKAGRQRQALIICNIKFEELCERVGAELDIKGMKKLLEDLDYTVQVERNLSA

TEMESKLKMFAGRPEHKFSDSTFLVFMSHGILEGICGTKYKKQEPDVLSYSTIFRVFNNI

NCPGLKDKPKIIIVQACRGENEGMAWVSDSLGPSATSSQEPEDLENDAIHRTHVEKDLIA

FCSSTPDHVSWRDPKTGSLFIVQLIKCFQNHAWNCDLESLFLKVQRHFETPKQKLQMPTR

ERATLTKRFFLFPDEGTEAQRSEVTCPQSHGRQVAESEFEPMTSDSQARALSTEPRCFRG

PSVCPLTVKHEGGERSNMAQRKEHRPRSRRDLDSNPNSATLASTVYGGSGEGVKGAQKEC

GMGQTRGNHSCMKSSGYAEGQDGRAISAVSWVSSFSDEVTEAQRSYTVTQVPSGRGGFEP

MTSDSQARALSTEPRCFTVSLFVRENSFHPPTIMNVFLN

>GRIA4

MEKAGQNGWQVSAICVENFNDASYRRLLEDLDRRQEKKFVIDCEIERLQNILEQCSAHSK

PSVNTIDRYTLPLNDGCKEKQANDKEYRSRTEFEFCVEDDDSGPGVPSTEHWGQYKQIGL

DTIPVPHGAHSLNPHFKDEVTEAQRSEVTCLRSYRSMVFIERLLYFEHCTTHLREYNREG
PSNQLRYESETTRRHLITTARAKSDFPTGWWPKLIFDPRHQRDRITEDTKADGGLRNDTG
YHSLRSEIANEKSDLLTMRSHTFGHKATLNLCLRKVAGKRGSATCQLCDCGQVTSLLSDL
ICKMGMKTVSLPWDNLMTLYLPQRLEQCSAPNKRLTNTNITITLLCQIVSVGKHVKGYHY
IVANLGFKDISLERFMHGGANVTGFQLVDFSTPMVSKLMQRWKKLDQREYPGSETPPKYT
SALTYDGVLVMAETFRNLRRQKIDISRRGNAGDCLANPAAPWGQGIDMERTLKQVRIQGL
TGNVQFDHYGRRVNYTMDVFELKNTGPRKEAAFKAHLREAVSICLTPVPGKLSDSTFHSL
YPESNSSSRGLSGLGDKILARTLLRRRLQIPLSERVQNPNAALAHSQHSRHDLGLTSITR
QARGTAPQPLYFTALPEAFDNIDKPEVGYWNDMDKLVLIQDVPTLGNDTAAVENRTVVVT
TIMESPYVMFKKNHEMFEGNDKYEGYCVDLASEIAKHIGIKYKIAIVPDGKYGARDAETK
IWNGMVGELVYGKAEIAIAPLTITLVREEVIDFSKPFMSLGISIMIKKPQKSKPGVFSFL
DPLAYEIWMCIVFAYIGVSVVLFLVSRFSPYEWHTEEPEDGKEGPSDQPPNEFGIFNSLW
FSLGAFMQQGCDISPRSLSGRIVGGVWWFFTLIIISSYTANLAAFLTVERMVSPIESAED
LAKQTEIAYGTLDSGSTKEFFRRSKIAVYEKMWTYMKSAEPSVFTRTTAEGVARVRKSKG
KFAFLLESTMNEYIEQRKPCDTMKVGGNLDSKGYGVATPKGSPLSSPVEIKVISSRGSDT
VVTPYEEVTI

>GRIA4-like2
MENGNANPYCTLVPGGLYFVWIRTVGDIKAWPSGFGKVQYIRVGRHVSCPPAAYSLEGET
DINTINEEILAAERLVKQGTKGNLEKETKLKKILHTQSRSKTGDLVVVTVSQLGRWLVSGR
PSATSQNSSALGSSGTGEIRGRRPRFTARKEAMVNHFRIFTKKPLWIRYQNDRRWRWGVL
GEMCPWCRYGSEMTRQHKTREAMQIDKECAQSASQNTQPFHLSTPIAQRNEARPGSSETE
RHVRFHSHPASTHASSSEEHVIRTIQPPGKPKVTVLASYWREAREDLISSECRILFRVTR
GFEGGERADLSHPEKDGPVRVEDMSEESNEGELGMRDICSQYSRGVFAIFGLYDKRSVHT
LTSFCSALHISLITPSFPTEGESQFVLQLRPSLRGALLSLLDHYEWNRFVFLYDTDRGTY
CWGSYKLIGLDTVHVPHGTRSLNSRCTDEGTEAQRSEATCPRSHSRQVAERGLKRRSS

>GRIA4-like3
MERRGPRKKGEEEARQARQAARDPRVATEPRIEGNMGHLSRSGGRRGFPEPGEPAATHSL
TFPNQFHRPAGGDAYLFERCSEKPPSPVPKIQKTTRQSVCK

>PDGFD

MSTNRGCDWGAGGSRVLSTVSVHLLEQAPSRATECLLGTFLGLPCGFIACANSLSLGKKG

NEEIFKLNPETSHRFKCNSDKVTEEQRSEVICQGHTAASLRGKKRSPVPSPRGNDAQYLS

PVEKSCSRNRRTRVQVPVCLLGMRHQWVNVSHPKNRPTRAFLEVDRANGNLGKELLRLCQ

LRPQWPLPVAVVFIESRLRSEPCFNTWETRALSTESRCLLRTYCAKGTALTRWERRVQQW

AFDKASKRGRVIVGYTEGGSETGRGREVGGGEKIESLNCNSRNPDRYPLGPSIGTDTNMK

VVIRQKFTDSIGTVGDLYRKEETIHVTGNGCVQSPRFPNSYPRNLLLTWRLYSQGNTRIQ

LAFDNQFGLEEPENDICSLGSGWRVEDNPLLVKTHPCWATAARESRAKTRVCCTGGGNDK

PLPHFYQENSTDPLSERSQMEGGAFWERCVRGVAMGRKRLDGIRQEVLWDQVPSSVKRGL

RLRAPRVTGTVPNSITLNLPQCFEKFMAHTLSTTLCPRCDTVNTISARTAKQNRQPLVGV

TSWTRSTSLSRPTLTMQPQWPGTGERSSIRSYLLSTYCVQSTVRSTWESTIQKKTDAFPS

HNKLTV

>PDGFD-like2

MTEIHLNPLKDTEFSIKIQTLKVSISKQVTEGDEGTEAQRSEVTCPQSHSRQVAELGFEL

MSPDSQARALSTEPRCCCSVTSDLKLIVGRENVCLLLYCTLPSTWYSALHIDNFQPAASE

TNWESVTSSISGIDYHPPSVTDPTLTADALDQTVAGFDTVEDLLKHFNPETWQEDLENLY

LETPHYRGRSYHDRKSKVDLDRLNDDVKRYSCTPRNYSVNLREELKLSNVVFFPRCLLVQ

RCGGNCGCGSPNWRSCTCNSGKTVKKFHENSFETLSLLRVYRRKGHTASLHEELLPKPWT

GRPPWPALPPVRPSLLLTEGLVPSRRWDSSYSNRSIRSDVYREGELEALKPSD

>MTMR4

MDCSPEPRRRRQIRRFLEDPEEAELAQFVQEFPGGDGGGGGGGCRRPEPEEPSSRDPEAL

PAAPEPDPRPPARPWPPDGHQHISAPAPLSPLTRPRSPWGKLDPYDSSEVGAGPFSPRQD

DKEYVGFATLPNQVHRKSVKKGFDFTLMVAESGLGKSTLINSLFLTDLYRDRRLLNAEER

ITQTVEITKHSVEIEEKGIKLRLTIVDTPGFGDAVNNTECWKPLADYIDQQFEQYFRDES

GLNRKNIQDNRVHCCLYFISPFGHGYDSPRPPPGQSEDRGRGPGLRPLDVEFLKALHQRV

NIVPILAKADTLTPPEVEHKKRKIREEIERFGIRIYQFPDCDSDEDEDFKLQDQALKDSI

PFAVIGSNTVVEARGRRVRGRLYPWGIVEVENPAHCDFVKLRTMLVRTHMQDLKDVTRET

HYENYRAQCIQSMTRMVVKERNRNKLTRESGTDFPIPTIPPGADAETEKLIREKDEERAA

LSAREGTDIGIGRRLPCPRGTHGLEGTAGVKIDSAYGPKSRGPEEGVDIGCLKDPIDRPV

VFIERSLCAANLGEYNATDLVDPFPAHEELTVQPRVRHSYTRRSGVAQWLEAGPGSRRVE

GSNADSATYQPCDPRRVCCPSLGLSALVGKMGMETVSPRVLCAMSALKKYGRVNEWMSVR

QDPGRGQGRRGAAWLGGSPEPHRMGSAMAEAPFCPGKSPGPGPPFPPLIGPGRVDFGQMG

QEGGQRDRQEKQVPGTQGPHPSRMLPGRQTAGAPAVAFGHPLRPGSGDAQVLPSGRFPLR

PPMSPGPAPSPKRKSRFPSFRKGEVGADGSCPVSRPPRAAIAPPPGPSFGLDLFPGPSRP

SPGCPRFPPPGRRRDPHSRPARRRRLTGDGPPLGKRVFIGVILEPVGAVLSPRAGILIAG

PGCSSRVGGQDTRQGARTLLGLRRDNHNDNSNDVSDAYYVPSAEKQRGSVGRGRAWESED

VSSDPRSATCLLGDLGPATQPKPREVPSSGKWGRSPEVTEPVEAKRVGEGHRVENPAGPP

ARPEGRGGRPRDPSATMNLSARGSGSCSVLSCFGEEGPPSLEYIQAKDLFPPKELVKEEE

SLQVPFAVLQGEGVEFLGRAADALIAISNYRLHVKFKDSVINVPLRMIDSVESRDMFQLH

ISCKDSKVVSALFLALPLPPAHPGSRPLRGRCHFSTFKQCQEWLSRLSRATARPAKPEDL

FAFAYHAWCLGLTEEDQHTHLCQPGEPVRCRQEAELARMGFDLHNVWRVSHINSNYKLCP

SYPQKLLVPVWITDKELENVASFRSWKRIPVVVYRHLRNGAAIARCSQPEISWWGWRNAD

DEYLVTSIAKACALDPGGKVAGGSACGGNGEGSEAGDTDFDSSLTACSGVESSSGPQKLL

ILDARSYTAAVANRAKGGGCECEVLPNCEVVFMGMANIHSIRNSFQYLRAVCSQMPDPSN

WLSALESTKWLQHLSVMLKAAVLVSNAVDGEGRPVLIVALAKILLDPYYRTLEGFQVLVE

SDWLDFGHKFGDRCGHQENAEDQNEQCPVFLQWLDSVHQLLKQFPCLFEFNEAFLVKLVQ

HTYSCLYGTFLANNPCEREMRNIYKRTCSVWALLRAGNKNFHNFLYVPGSELVLHPVCHV

RALHLWTAVYLPPSSPCTLREESVDLYLAPAAQSQEFSGRSLDSCSVCSGPLCALSGCSD

LAWDSGAS

>ABHD11

MLRRACAWRLRPSRGLSLARAWSNEGPRPVPLSYTQFDGPTQEAPLVFLHGLFGSKTNFQ

SIAKSLARQTGRKVLTVDARNHGESTHSSEMSYEAMSADLQALLSQLGLPRCVLIGHSMG

GKTAMTLALQKASGPGPERPTPCIRSGKSRRAQLERGGSVGERWILRSVKVPFGSPPPTR

SCPYHGGGRSLEGQGEPELVERLVSVDISPEETTGVSDFPSFVAAMQAVRIPKELTRSQA

RKLADEQLKPVIQEVSVRQFLVTNLVEAAGRYVWRVNLEALTHHMDALMGFPQLPGTYSG

PTLFLGGSNSQFIRPSHHPKIRRLFPQAQILSVPGAGHWVHADQPHDFTAAVRDFLT

>IL1B

MPAYGEQDSSREVEETDDKKTKRQPGLESTMARVPDQSRDLMECYSGDGEDQFYEVDGPS

QIKSGFQDLKARTCQETRIHKEDKCSPCQMGIELKVTELPSSHGFRKAVVLVVAVERIKR

QAVSYNTSFMDRDLMDIFTSIFKEEPISCSTWEQTLVTDSLYHYLRCQEVTIWDEEHKSF

TLNTMANPCELRALHLIGANATQEVKLNMNFYYKTERLAGPTVKQPVTLGIKGGNPGNLY

LSCVKKGGKPTLQLEVVNKSDLLGKNQERFIFNKSTEGTSTTFESAAYPDWYISTSREED

EPVFLGASKGEEAITNFFLH

>NEK7

MTGPNRILLPAQILLPFLIYAYGNPTLFSDLTSEKVASYLICKMGIKTVSSTWDNLMTPY

LPQRLEQCSAHILSMIPSNLAVLSKVTTELLLPEKQRGLVERARAWESEACISSCLQDVS

TWMSARHLKLNMSKTELLIFLPKPCPLPDFPIILDGTTILPVSRARNLGVIFDSALSFTP

HIRSVPKAYRSRLYNIAKIRPFLSTQTVIIPKALRPDMGYNTLANFRIEKKIGRGQFSEV

YRATCLLDGVPVALKKVQTFFWVFYTPSDEEFVHSHGFSPSSFNLCSEDLILKNRVFFGP

NFLGLLHLLCSLSEMKLSYLDAKL

>GSDMA

MLHAARHLLLLLLRLLALGTGGSPTSPTPTWTPARGCYRAEEDGEPTFRCSYAGLGAIPE

GIPNDTRKLFLDANQLGEVPAGAFEHLPVLSELDLSHNAIARLSGAAFRGLEGSLRLLDL

SANLLAAVPAEAFSGLRAATNLSANPWRCDCALQRLLRGMKLAEGTGAGIVCATADRPEL

VGRQVLGLEGEAGPCGARQGRRGTDAALLVTVGGWLALVGVSLARYVRRNHEEVRYLICK

MGIKTVNRMRDNLITLYLPQRLEQCLAYKAPPSPPLYCSSLEADFPSGFQAGLDLPEYSW

EEALRGKRRGLVLAGPWLGSEGSLRCSVVPKCHLLGVSPAGGRSNIPFIPDKKQKTFPSE

TIGSKSLEEEEEEERDFRVLQAEVEMEMYALKALTLGQREGLLTTLLGIMGQAKALQTLE

DTVEQALDMEEPVQLAQPGSAILPILKEDSGRLNPTLSGTVLYLLGALRELSEEQQQLLA

MSVEKEILPQQMRLLETILEQHFLREESGPGQLHSALLSGLRGEEWAVTQALLSLSGLDL

DENELLFTFDPEALPQLSALYAALSIFHLLAKTCSFSSSGLPTQPGILASLLPQGPDWGQ

GAVACSPSYLICTVRIKTVSPSGDVDCVQPDDLVSTPVLSAVSGRRARLRQTGPCCSPRR

RAPAEGEEEGDQGCSSVGLGEEGGPPNLNVSTHSPRTMTAMFENVTRALARQLNPQGDLT

PLDSLIDFKRFRPLCLVLRKRKGTLFWGARYLPTDYALLDLLEPGAAPTESTDNPHFRFK
KLLDLRLEGKVDVPNTVKVTGGAGLTQSSHLEVQTLSVAPKALDTLRDERADQGPTPSRK
LLPEHPFLQELRPRGENLYVVMETVETVKEVTLERAGQAQGGFSIPLLAPLGLQGSLNHQ
EAVTIPQDCVLAFRVRQLVTKGNGEWDIPHVCDEKLKTFPPEDKTEEEKFTCESVEGAEV
PEDFGALQEEVESEARQLAQLTPEARATLLRSLRALLGKNHQLRLLEGSLEGALHKGTPG
ALEDLGNVTLSPQAMGSILYFLGALTELSEAQQKLLAQSMEKKILLTQLKLVERAMEENF
QQSQGGDFPLPPQLLSSLGDEDLTLTEALVGLSGLELHRAGPRYTWEPATLPRLCALYAG
LSALHLLAAPAS

>GSDMD_like
MTFSPTVTENSQPGKTRSGVGRCRIHGLAWSSHRWRRSGMPRLVFMPRPAFFRRSHGGRY
RPRHGQGVLSHHPDPTADPSWLPLQAIRVTGRSLRRGRKICFLSWDRWSEEVSSVVLQSG
LHVAKPYFLIKMKGSPQPPGSMAPIFAQLTKNVAKKINSEGELLPLLSMNNSKRFRPLCL
VRKKRKGTLFFGARFRPTNLSLLDVLDSDLPAPELKREDKFGFQDRVDGRLKGKVDLRDS
LLSVQVSGEIKRVQNYSLEVQIVLISPEDLDKMQKERKLKKNEPEELKELRRLGENLFVV
TKVVETLEEANLSSERQAEGGCLLKLLSIHMKVQGPIRSGRGKTGRPSCHLPDLHLWSQS
INSWYLLSANYAQTTLALHNHQEVVNIGKGCTLAFGLGHLIFRDKWKILSMPSKEKTFLS
KGLEEDPFVKDLAMAKDAKGFEGLRQEVRQEKQYLIYLDRQLKETILQAVQDLLGQREEM
QKVEDALEDAMDGKATQKLEGPGNILLTILKEDSDHVVPELTGTVLYLLGALLVLSDTQQ
HLLKLALEKKLLPQQLKLVESILEQTFPMSQEGHFFLTPGPEDEERSFTTALLEQYGLEL
SGPNSQFLWKPDALASLSALYGALSLLDRRN

>NAPRT
MAEAERAASPLLTDLYQVTMAYGYWRAGRARERAHFDLFFRQCPFGGGFALAAGLRDCLL
FLRRFRLRDPDVDYLASVLPPDTDPAFFDYLRGLDASEVTVRALPEGSLAFPMVPLLQVS
GPLPVVQLLETTLLCLVNYASLVATNAARLRLIAGPEKRLLEMGLRRAQGPDGGLSASVY
SYLGGFDATSNVLAGQLRGIPVAGTLAHSFITSFSGQERLQSGALAPGDLSAQAETWLTR
VCELLGRPVKDAHPGERAAFVAYALAFPRAFQGLLDSYSVMGSGLPNFLAVALALADVGH
RAIGVRLDSGDLIGQAQEIRRIFRTCATRFQVPWLEFIPIAVSNNVDEALLAQLAQKGSE
VNLIGIGTNVVTCPLQPSLGCVYKLVAAGGRPRLKLSEEKEKRTLPGCKAAYRLGGPDGA

PLMDLLTLVEEPPPQAGQELRVWPLGSGEESRTLTPATVETLHRLYFQRGQASLEISPQE
CESLPTLTQARAFAQESLSRLSSAHKRREAPEPYQVALSEKLHALLESLSRSSRGLSLIC
KMGIKPVSLTWDNPMTLYLPRRLERCCAQNPEEFLTLLMRHVLGLEPLLRLQCGGREELS
YWSGQRSWWCPASSSCLSLPSWAGLRLAEGGVGLAVTQFRGNSLGLWVATARAQLTADRS
GTHKQIGPRGTWKSSEEAALWIRGIIIVIVVAALVAASPVCSERLDSSLPCGLCDRRHRA
ALLRASPGPLGPAGGPLG

>MROH6
MAAGAGEGVRPELGEAEGPGNPEPPPQPKAKPTRARGGRPKV
TRRPRPAVAPGSQPPPCPVGALTLAALAEEIQSHRGDRAGPGQRPGDRGDGRTAEPASRP
PAPEGNGGRRGDRATASGEAGEDAASSPGNDQRRPQKRPPKQPRPQLPQGEARPECRPSS
PAQSTFHPLASASPCAPGPEQFPLASCFLTDLAVHTVACLTDAGFSGTQATAVCLSSTLE
AHGTILRDKVEELVHGLHLQIHRFSEGRARRAALRVLCSLAVEHAPDVVHGLLSHSLPCD
SAVELWRGLSRNQRVNVTVLVQLLWKLKGQPRVLGGSPAGPDGALQEPLAATRALGEMLA
VAGCVGAMRGFYPQMLIALVTQLHQLARCPPDNLSKARGPPQSKAAHPRSHAHCAVEALK
ALLRADGGRMVVTCMEQAGGWERLSGPDTHLEGVLLLASAMVAHADHHLRGLFADLLPLL
RSPDATRRLTAMAFFTGLLQSRPTVRLLRAGSILERLGAWQGDPEPSVRWLGLLGLGHMA
LHAGKVQHVEVLLPALLGALGEADGRLVGAALGALRRILLQPRGHSCTDSICPDVGARLW
PLLDDARDPVRSSAIGLFGTLVGRSPLLQRCATRDLVLDSLVPLLLHLQDQSPDAAEQSA
EWTLARCDRFLHWGLLEEIVTMAHYDSPEAFSRTCRRLVRWYPGRVPGFLDQAQGYLRSP
QVSIRRAAGMFIGFLVHHTDAGAVKEGLVDSLLHTTLLLYATGGQLLNEFSPEHLVQCSA
LNNYEGRISGDLRELECDPEASVRSATHVTLHQLRLASQDWASRPGRFSPRRLLRPRGRP
ARPWPLYEEGPFKRRSRAGLWGSHMGA

>ZC3H3
MEMEEKEQLRQQIRLLQGLIDDYKNVHGNSRAQPAAAAGPRWPLPAYRGRGTFGVGYPRP
VRGDFFPHRGLSWRKKYSLVNRPPGAAEQPEGRAPPSQDRPGPSPPDPPRRVRLGPDQNV
VVGIEAPSDPGSAGGSRTHRDVPRSDPGLQKKEGGAGASNGEEDADLVCRKERGDRRVGN
SAGSGPGGPGEPRRTVSENARGLTGQAPPARPRSSEGAAGGKAGPPVPDALRLQRLRPGR
EPPLRNSLAQAPLDVSGPGRRAPATRTAREPSLPGPCRTPKFKKTNYTWVASTVKAPRGP

PRRSLSPRAAAEAARRAPSTGAADGPIKPQPKADPAVRPRKPAAPSKPGGPSSKYRWKAA
GPTPATAAAFQWRAEAPGRSDAPPASPDRADLPAPSQASGGPSGWKPAFGETALSAYKVK
SRTKIIKRRGSVSLPGDKKSSLLPPATPKSHYSLRRKHGARAKSSPVLKKNPNRGLVQVT
KHRLRRLPAARAHTPGKEDCSRGSTSSDGFSREVGRQASAQNGARHIGRFTPVRAALPAP
VAAEPEDWVCLPVLKSGAFPRAGGQKAKPWGESHAIRSPRAGDAPLYFEVPLSRAPSGSE
LPISPLGGTDGRPDDFRRGSPRRPTGGGRRSDGRDPPGPAPHPSRRALAPGADTGRRASA
G

>Syk-like1
MSRRAACGPVRGPWRASPWASLCASRVGQAVSRPWRAGPCADRGVRARVRTVACEPVCEP
RGAGPCAGHAVRAACGPVCGPWRAGPCADRGVRACVRTVACEPVCEPCGAGRAPAMSCGP
RAGPCADRGVRARVRTVACAPVCGPWRASPCVSHAAQAARSPCRVGRVRARVRTLAFGPL
CEPRGAGRVPAMSCGPRAGPCADRGVRARVRTVACGPACGPWRASPRASHAAQAVCRPCR
AGRVRARVRTVACEPRGAGRAGRGVRAPVRARAPAMWGGPCANPCANASHVARAVCEPAC
MTGWRGPGPSAGRALRGRIHFPKPGSRGSPQCPSPPGPGVHFCFFRRGPATNRLHAGRNA
LPGPGPVPGGVCRVHGAVPDAHRVPRAEPDVSRVQSAELDASFGGQSAELGVYRVQSAGL
DAYRVHRAELDVYRVQRAVLDVSRAQSAELDASFGGQGAELDACRVQSAELDAYRVQSAE
LDAYRVQSAVLGVYGVRSPVLSAGERTAERVLFLERLRGAERCAERWGEDAIHSFVRPFV
HSFIDPFDGGASRPSGEEEEEDGEEEEGEEKEGEEEGDEEEEEEEEEGEEEGEEEEEKRK
RKRGGGGGRREEEEGEEEEEEGDEEEDEGDEEEAKRKEEEAKRRKGKRKGRRRRKRRKGR
GRGGGKEEGEEEEKRRMERKKRRKETRKRRETEEEEEEEKRRKRMKEMRRRRRGGRRRRR
RRGGGRGGGRREEEEEEEGKRRRRKRRGEGREEEEEEEEGEKKRRKRRGGGKEEEEEEEGKR
RRRKRRGEGREEEEEEEEEEERRRKRRGRGGKEEEGGGGRREEERRRKRRRRGGRGGCCVGF
EAAGGPGRQSRLEAREPPGGRPDDPVGARRIPSPRLFQVRAGGGARMASGGGEAAAQLPF
FFGNITREEAEARLEEAGLGEGLFLLRQSRSSLGGFSLSVSSGGRVHHYTIERDVAGAFA
IAGGRSHPGPAELCAFHGREADGLVCRLRDPCLRPPGLRPRAGPFEGLRETLIRDYVRDT
WNLQGQALEQAILSQRPQLEKLIATTAHEKMGWFHGAVSRTQAEDALLAAPKADGKFLVG
DDDDDDDPRQLERSLAHSKRLTDATDYHAPDWSIPSAQSSALRIIPSMMMMMMMTPPPV
LSTEPDTRVRSRGPEGSFALCLVHGGRALHYRIDRDKAGKLSIPDGKKFDTLWQLVDHYS

YKADGLLRPLATACPRAGQAHDGQRPTRFRRGGGDGRGGGTLPGGTPPPPRTGVTTPNHP
AHLRPALPPPPPPHRASSAALCTQ

>Syk_like2
MRARACVCSRVSARVCGFQAGGLIGRIHSFQRAKKGYLLSADCVRSTVPNAWKAQFGNRE
RPSQPNNGLRAERCAGRLLLDASYYMQRLGWTPPTGRLLLYAERWAGRLLLDASYCMQSA
GLDASYRTPPPVCRALGWMPPTGRLLYAERCAGRPLYAERCAGRRLLYAERCASQGACSG
PFLPPSPFVRMLGSLLILSRPAKRCDEHLRAENAVVGVSRVQSAVLDSHYVQTAVLSVYY
VLGVGSVQSYVPGAYRVQSAMHIVSTDRALSAGSQSPSNPLMRLYKETRDALAHAKLGPH
AAGHGHQAAVNRLESSAELNPYVMQRGRRGAGQPLPEELREALPMDTAVYESPYADPEEI
RPEAVELDRSLLTLEEGELGAGNFGTVKKGFYRMKKAPPQTLRDVPDAPKESPQNPPDPR
KPPGPPKRPPGPPQGPSESVQSPPRTPKNPSGPPQNPPDPPQGPPPRAPPQKVQDGPGRH
TRARTHTAPAPVLSVSGSVAAPERARVCGGRWRPRRGDKAVAVKILKDGGGGGGGGVDEA
VKEELLREADVMRRLDNPYIVRMIGLCRAEAWMLVMELADLGPLNKYLQKNRHVQARNLT
ELVHQVCMGMRYLEENSFVHRDLAARNVLLVTQHYAKISDFGLSKALNADQNYYRAQTHG
KWPVKWYAPECINYYKFSSKSDVWSFGVLMWEAFSYGQKPYKSSGRPIRQQRERVQHNNP
APNELTFHARVPGFPVYHVVCVRVSPCRVTSPVCAPCVLCLCVRETHVCASETLRPVPQG
MKGSEVSAMLEKGERMQSPEGCPVEVYDLMKICWTYKVEERPDFAAVELRLRNYYYDISN

>Syk-like3
XELREALPMDTAVYESPYADPEEIRPEAVELDRSLLTLEEGELGAGNFGTVKKGFYRMKK
PSYGLPQTPRTPQKHSRSPVQSSRPPTDPHKPSGTPSDPPTGPREPHRRPPGPPGPFMDP
PQALSDHPVPQGPPHKPSGTFQTPPRSPPKTLQTLANRQDPPKDLQVPPKDPPSPSSPPP
GPLKTLQDPLKTLQTPPKDPPPRAPPQKVQDGPGRHTRARTHTAPAPVLSVSGSVAAPER
ARVCGGRWRPRRGDKAVAVKILKDGGGGGGGGVDEAVKEELLREADVMRRLDNPYIVRMI
GLCRAEAWMLVMELADLGPLNKYLQKNRHVQARNLTELVHQVCMGMRYLEENSFVHRDLA
ARNVLLVTQHYAKISDFGLSKALNADQNYYRAQTHGKWPVKWYAPSASTTTSLQQSDVWS
FGVLMWEAFSYGQKALQGAARSPAGDRSSGRPIRQQRERVQHNNPAPNELTFHARVPGFP
VYHVVCVRVSPCRVTSPVCAPCVLCLCVRETHVCASETLRPVPQGMKGSEVSAMLEKGER
MQSPEGCPVEVYDLMKICWTYKVEERPDFAAVELRLRNYYYDISN

>ROR1

MALALVCCVTLGKSLLCLSALICKMGIKTGSPMWDRDYVQPDLLVSNPVLSKVPGTQDEY

EEDGFCQPYRGIACARFIGNRTVYMESLHMQGEIENQITAAFTMIGTSSHLSDKCSQFAI

PSLCHYAFPYCDETSAVPKPRDLCRDECEILENVLCQTEYIFARSNPMILMRLKLPNCED

LPQPESPEAANCIRIGIPMADPINKNRKLLEDKGRCLLPPLDSPKCSALCPMHGKCSVNS

LSRLISDHKCYNSTGADYRGTVSVTKSGRQCQPWNSQYPHTHTFTAVRYPELNGGHSYCR

NPGNQKEAPWCFTLDENFKFDLCDIPACNLTDPRGRFCFVTSHFRRCGTIQFRAAQMFSR

LWARQFLTHFVSSFPPKFRDCPLQDSKESKEKNKMEILYILVPSVAIPLAIALLFFFICV

CRNNQKASSPPVQRQPKPVRGQNVEMSMLNAYKPKSKAKELPLSAVRFMEELGDCAFGKV

YKGHLYLPGMDHTQLVAIKTLKDFNNPQQWADFQQEASLTAELHHPNIVCLLGVVTQEQP

ACMLFEYTNQGDLREFLIMRSPHSDVGCSSDEDGTVKSSLDHGDFLHIAIQISAGMEYLS

SHFFVHKDLAARNILIGEQLQVKISDLGLSREIYSADYYRVQSKSLLPIRWMPPEAIVYG

KFSSDSDIWSFGVVLWEIFSFGLQPYYGFSNQEVVEMVRKRQLLPCSEDCPPRMYGLMTE

CWNEVPSRRPRFKDIHGRLRSWEGISSHTSSTTPSGGNATTQTTSLSASPVSNLSNPRYP

NYMFPAQGMPQGQIAGFIGPPLPQNQRFIPINGYPIPPGYAAFPAAHYQPPAPTRVIQHC

PPPKSRSPSSASGSTSTGHVTSLPSSGSNQDANIPLLSHVSIPSHPSGMGGPVFGNKTQK

PYKGDSKQPSLLGDSNLHGHTESMISAEL

>AUH

MAAAAAAAGVGLLGVGRGARGRFRFRLRAASASGAVRGLGSEPREEDELLRLRFLPDQDK

DEGTEARRSEVTRPQSHSRQVAELGFELMSPDSKARALSPEPVRMHVCLCVWTRVPASVD

ACAHVCMSVGAGAHVCMSVGPCAHVCMSVSARAHVCMSVGARAHVCLSVGAGAHVCLSVG

PCAHVYMSVGACAHVCMSVGAGAHGGMSVGACVRVGTRVHAWGQVCPLSSPGIVVLGLNR

PQAKNALSWNLIKQLSQSLEALKSDKKVRTVIVRSEVPGVFCAEECSAPSRRLTDTDLMI

RRGRRVCEEEEEEEEEQEEEEEEEKEEEQKGEEEEEEEEQEKEEKEEEEEEKEEEEEKSRPEGAS

QDGGRTAQLPVPTIAALDGLALGGGLELALACDIRVAGGSILWKAPAPPNGLFRPPEVPA

QRAPPTGSPGSARCSIGCPVELQARTLGARLIILAWPPLPDGALSARLVALRRVGGLQQR

VLGVRLVALQSVQLQVRALPAPPGLGRMQLQDGALGARLVALRRVQLQERPLGPRFVALG

RKRPPRPPGLWIRTGRFRIGRSDCRSSPWGGWGCLSERSGLDSSACASSAKMGLVETKLA
IIPGAGGTQRLPRTVGPALAKELIFSGRLLDGDEARAAGLVTHSPPQNPRGDAAFLRALR
LARDFLPQGPVAVRAAKLAIDQGMEVDLATGLAIEEACYAQTVPTKDRLEGLQAFREKRT
PRYKGE

>NFIL3 (partial sequence)
XMTTVASNEVAEARRSAAAPPPDGPPPAPAGLRRPRPGPRAAMEALKPVPPRDDPEEDPA
AAVASAAPGPPPARLPPPLPPPPPPPPSSSPSSSSRRRRREFTPEEKKDAQYWEKRRRNN
EAAKRSREKRRLNDLVLESRLVALGRENAALRAELLALKARFGLLPPPPVKLEPPEAGSP
RGSEADEAGGGKTPSDGEDEQRVPKGPGPAAPAALPHKLRLKVRGAAPLKREGSEADLPP
PFALQVTRLQAWGLWPPPAPDGGGLSDRGAALRGLGGGRAVAASDSGPAGKRPVRSPDHL
SVKPLDKSPVGQAARQKTCQVIDPSVKPPNGLSIRPPKQQPKNLPGRSPNHRSIEPPDSP
AVRTPDKSGTRRPNPPTSRSINNPSRPRIACQSNHPTDNRPVVRTPRTTRRRTCQIAESA
VSQTTRWPINQTPPPPPPPPRPAGKRPVSQITQSAVIQTTGPARSMASRQPGKEAVRTPD
E

>DIRAS2
MPEQSNDYRVVVFGAAGVGKSSLVLRFVRGTFRETYIPTIEDTYRQVISCDKNICTLQIT
DTTGSHQFPAMQRLSISKGHAFILVYSVTSKQSLEELQPIYEQICQIKGDVHKIPIMLVG
NKSDESQRELDAGEGEALAARWNCSFMETSAKMNYNVQELFQELLNLEKRRAVCLQVDGK
KAKQQKKKDKLKGKCSVM

>ADAMTS10
MIKQSVLHHPMQALCTILLISLILQLMDHEPSDPLAHCRSSSPMERGNQTSISEFLLLGM
SNQVEQRQLLFLLFLWMYLLGVLGSLLIIILVIVSDPHLHTPMYFFLTNLSLADVCFLST
TVPKMLVNIQTPSKSITYAACLVQMYFFILFISLDHFLLTGMAYDRYVAICHPLHYTTIM
SPRLCGLVLAGSWLISSLHAPTHTLLVVRLSFCSNREVLHFFCELYHILKLSCSNILINE
VAVFVAAVVIGLAPLTGVLFSYTCIIFTILRIPSKGGSQVLDHAKYLASGQHPSQRQHSS
CCRFPPPPPPHKLLTAMKGALGDGVIQEGQLGGNEDSGPIRVKLQGTGEPQETTTASRKLL
SQLKNKQPQSRSGHQCPMAVCFALAEEFLSSLKSYEITFPVRVDHNGAFLDFAPPQRQRR

SLGTRPPEPAEPRVFYKVEAPHTRFLLNLTLTSHLLADHFSVEYWKRDGLDWRHHIRREC

LYAGHLQGQRLSSKVAISNCHGLHGLIVADEEEYFIEPLSGRGSGVPEGEGSPHVVYKRS

SLQRPHLDAACGVLDEKPWKGRPWWLRPLKTAPTKPLGNQTQRGQLALKRSVSQERYVET

LVVADRMMVAYHGRRDVEQYVLAIMNIVAKLFQDSSLGNIVNILVTRLILLTEDQPTLEI

NHHAGKSLDSFCKWQKSIVNRNGHGNAIPENGIANHDTAVLITRYDICIYKNKPCGTLGG

MCERERSCSINEDIGLATAFTIAHEIGHTFGMNHDGVGNGCGARGHETAKLMAAHITMKT

NPFVWSSCSRDYITSFLDSGLGLCLNNAPPKQDFVYPTMAPGQAYDADEQCRFQYGVKSR

QCKYGEVCSELWCLSKSNRCITNSIPAAEGTICQSSTVDKGWCYKRVCVPFGSRPEGVDG

AWGLWAPWAECSRTCGGGVSSSTRHCDSPRPTIGGKYCLGERKRYRSCNTDDCPPGSQDF

RELQCSEFDSVPFRGKYYTWRTYRGGGVKSCSLNCLAEGFNFYTERAAAVVDGTPCRPDT

IDICVNGECKHVGCDRILGSDLREDKCRVCGGDGSSCETIEGVFAPTLTEGGLEHVQGQL

SDRPKPPPCVLQATTPRAGAFSELRLTTFLVEEDPPRVTALPENRPHHNRPVGLRSVSLP

PPALLNFQARCPATPAPSPGYEEVIWIPKGSVHISIRNLNLSLSHLALKGENDAFLLEGK

PGPSPQLRLPLAGTIFHLRQGADQPECLEALGPTNATLIVMVLVRSDLQGIRYRFNAPIT

HEALPPTYTWHYAPWTKCSALCAGGSQVQAAECRKQPDSSPVPSHHCKAHAKLPERQRSC

NTEPCPPSWAVGNWSGCSRSCNRGARTRSVVCQRRISPNEEKTLDDSACAQPRPHVLEPC

SSQSCPPEWAALDWSECNPSCGPGLRHRVILCKSGDHSATLPASQCSAATKPPTSMRCNL

RRCPPPRWVAGEWGECSAQCGFGQQLRSVLCTTHTGQLSGDCTAALQPPATQQCETKCES

SPTESPEECRDVNKVAYCPLVLKFKFCSRSYFRQMCCKT

>MYO1F

HAVRPGPTWAPLGYQWNLYCE

ELRPRGPHRTPPTTALLISQALLRCLLIIVIIVIVIGATPDCIPLSARDPTGQRAPKSVP

TMSFSLIQGTGQSPLYPSGPRTKQNGQNQAVPFGALRNSPWRARLSPRPPTATPHTSNSQ

WVLGNPVDVAGDCKLAVGRECACYIVTLYSALHTGSKERFHWQSHNVKQSGVDDMVLLPR

VSEEAIVENLKKRFLDDYIFASSQEGWGRGGGTKSGGEGRGWSEFPQSDSHWLNVKATQE

KVKQRDSPEIVALDPAVQTYIGSVLISVNPFKQMPYFTDREIELYQGAAQYENPPHIYAL

TDNMYRNMLIDGENQCVIISGESGAGKTVAAKYIMGYISKVSGGGDKVQHVKDIILQSNP

LLEAFGNAKTVRNNNSSRFGKYFEIQFSRGGEPDGGKISNFLLEKSRVVTQNESERGFHI

YYQLIEGASQDQRQNLGIMTPDYYYYLNQSETYKVDGTDDRSDFHETLNAMQVIGIPTEV

QQLVLQIVAGILHLGNISFREEGNYAQVESADCESALQAIQWRGAGARGRGPGKKGPERF

LGIWTLTPVLAFPAYLLGVDSGRLNEKLTSRKMDSKWGGRSESIDVTLNVEQAAYSRDAL

AKGLYARLFDFLVEAINRAMQKPHQEYSIGVLDIYGFEIFQRNGFEQFCINFVNEKLQQI

FIELTLKAEQEEYVQEGIKWSPIEYFNNKVVCDLIENKLNPPGIMSVLDDVCATMHATGG

GADQTLLQKLQAAVGCHEHFNSWSSGFVIHHYAGKVSYDINGFCERNRDVLFSDIIELMQ

SSEQDSIRPNEPGSTRTLNAVLDTYAFIRMLFPEKLDADKKGRPTTAGSKIKRQANELVS

TLMKCTPHYIRCIKPNETKRPRDWEESRVKHQVEYLGLKENIRVRRAGFAYRRPFQKFLQ

RYAILTPETWPHWRGDERQGVQHLLHSVHMEPDQYQMGRTKVFVKNPESTPHRIQSLCPT

AVFFIPIARSPPLMLHARFYLSGHILGFGSQVPTGQIPGTPALNLLPTFLPQLFLLEEMR

ERKFDGFARTIQKAWRRHVAVRKYEQMREEASNILLNKKERRKNSLNRNFVGDYLGLEER

PELRRFLGKRERVDFADSVTKYDRRFKSIKRDLILTPKHLYVIGREKVKKGPDKGQVQEV

LKKHLDIQVLRSVSLSTRQDDFFILHEEAADILLESMFKTELLSLLCKRFEEVTQRTLPL

TFNDTLQFRVKKEGWGGGGSRNVNFSRGSGEMATLKVSGKTLMVSIGDGLPKSSKPTKKG

TPQSQSRGRRPAPARSAPGPPRGTCRNGAPPPMAPSSSQQQLEQMYAGHQKQSRGPPAAM

LPKQGASRRTRARPPSEQNLEFLNVPDQGMAGMQRKRSIGPRPPPGVGRPKPQPRAPGPR

CRALYQYVGQDVDELSFNVNEVIDILMEDPSGWWKGRLHGREGLFPGNYVEKI

>TMEM131

MQGTFPPPASGSERNRSTPAADVARRLAAGDDVGRDLAGEEASILVEIISRLLRGSEEAC

NVIRPSLRKAVKSCSNNNNDASKAKRPSQFSGKITVKAKEKSYSKLEIPYQADVLDGSFV

RKHSGSFFCAPFANRGVIKLTSGPRILSTFWSWSVVWYSPKRLVQCSALDTHSNTVDGKR

GEAEEKSRAGGEGKLRSGEDDASRHGKSRFASVGNFQCLERRCPAAFDPVKLFLSGLTGV

IFRRDLGPVVSSGQDSSSDPIERPVYLTNTFSFAILIHDVLLPEEARIMFQVQNFSKPVL

IPPNESRSVFTLFFIPSSSSVHIDSNILLITNASKFHLPVRAYTGFLEPQIKTVPTQRRA

PGLNPHFADEVMTMVFVKRLLCAEHRSKRWGRYGVISSSHYFVVHPQAEEPFIDFGILSA

SEASHILFAVFNSNPIENTRVNIRNRFIYTNAHLLLYSLAIKGWHVIGDGLTVELLAAER

GNRTAIDARIPELVDASASAQSSVILASGYFAVFGVKLIAKDLEGIHDGAIQITTDYEGL

KPLKRAEDPSGPRTRIYFRSVCDLSRKSVFAGFPFMALKQPGDGI

>CLEC_like1

MAKTDLLIFPPKPGPLPDFPITADVTSSVKQRLRLSAPYGIGTMSNMISLYLPRVLIKQC

LANKMASEVTYADLKFQDSFKAQRIQEFDNIQEIEHPALSPVWRCSALGLLILCLLLLIG

LASLGILCDAAVTRPLLRTRVFQAKKTNTEQLNGLEKNLSLQLETIANISKEKEMIQSNL

TDALQEMATKLCRELSSSKPKSQICSSNVSPTTPHRHQSEKSHLRSVYGIHPADPATPPS

AIREWERVGFGMGAGPTKKPRATDFHSVACGFHRGLIAAAWGVASTRPLLRTRVFQKKKT

NTEQLNGREKHLSLQPETTANISEEKEMIQSNLTDALQEMATKLCRELTKNKQEYNDAEF

SDPLRSRWGTEQSGGHAGRGWRLKQYSASGKCSIITIGSLTDNVYEPICKLRYWSLPDAL

NNYNRLIDCWEEGMMPSDLRQKNVLPNKMREEKHPAPSPARRCSVLGLLTLCLLLLIGLA

SLGILCKYHWMGLEARKTNTEQLNGLEKNLSLQLETIANISKEKEMIQSNLTDALQEMAT

KLCRELSSSKPDSQICSSNVSPTTQHRHWSEKSQLRSVCGIHPTDPATSPSAIRERERSF

ISLQYNQYIWVGLSRNSSSSQWKWEDGSALSPDLISFSSKDTTVGKMCATTCDNVFHSSL

CTNNRYYICEKPAGLVKKFTAD

>CLEC_like2

MCQNGFQGWCLTLILKLFLNSFGILLAFHIIIFKKKSSTRPLLDTAVTRPLLQTRVFQAQ

KNIEQLNGQEENLSLQLETMANISKEKEMIQSNLTDALQEMATKLCRELSSSKPDSQISS

VTSICERGMKIPWITSSVRLLRSYARAAERCWRKSKHQADLTHFKFILSCLNSALSSARQ

NFSSLIDTHARHPRRLFRTFNSLLRAPVPPPPPSLTPNDLATYFLMKINTISYTHSLGEL

IRSHGFDYHLYADDTQIYISAPVLSPSLQARISSCLRDVSTWMLARHLKLNMSKTELLIF

PPKPGPLPDFSITVDGTTILPVPQAHNLGSSNVSPTTPHRHQSEKSQPWSVYGIHPDDPA

TPPSAIREQKREGRSWDGTRNFISYFQCNYNIWVGLSRNSSSSQWKWEDGSALSPGLLKF

PSYALNERKTCAYISAYHLSIDSCTNSHYYICEKAAGVVKKLTSV

>CLEC_like3

MGGDDDGGDDDDGGDDDDSGKISQTHPDLSRELSRAATMQDEDRYNVLNIPRRPFSQGPA

PTDKASSSSSAHSHIWRLLSLLVLILCLLKLIGLALFGIRFFQAQSQENGEATSPNDKPG

NPHADRSWSCSKQLEHLSSQNSNLSAALREMSTKLCQGITRNQPDVLDDFVEYNYEPYPE

RWICARDNCYYLSMVTKKWGESRKSCEALNSTLVKIDNKEELSRLVEYWWKLSRGDPERA

LNIWDNTLKTEDMIPVLRDLYNLMGETGMANEVTYADLKFRDSPVAQRIQEFDNIQEIGI

GSPRDSTFIFLKNFPINRFILHSRHQHQKPRATDFQSAASPVASPELSVGDAVTPPFLWT
GVFQAKKNIEQLTGVEKNLSLQLDISANISKEKDLIQSNHSFALKKVATKLCRELTSIKQ
APDSLSNLVSDHACKPCPENWHWHRDSCYWKTSVLNLDESRKVCAERNSSLVKIENKEEL
GTDLKVVGQKQEGPGGPVGIIRGELQVPPPASNNADKVLANRSLLLQLGGIPQNVIGTSQ
TPKTSSTNKLFVKERTKKKFHSSVFSDVPVHSVWRLLSLLLLILCLIQLIGLVGFGIRLG
RDSRQLAAGHLRQERGVPALSAPSEEPWDSLTYDLYYSGKMDHNPRAIPAVLTDAAFSNA
SLSLKIPLAPTDSFLFRGICRSCFKQKEFLLSQNSKLSADLREVTTKFCRKLMENKTAAL
DTAVPDTPRRESLRSAGKHNPRYLFLQYFVTAKIERYHWMKEEVTYTELSFHKTDDVEKV
LQMDLTEKPEQPAPRSLRSRVPLGQLPMCLLLLLLLSGLIVLGILYISLYFFSQNQTVSP
VKRDGNGLEPAVIENLRSGLPGQKERSFVTRGNMVLDDIYICSHAHIAGSSWEAMGEARM
VPDSGDGVDGVRFVDS

>CLEC_like4
MELTDPRAENMEDEDEDGYIMMSIQQQTSIKGLAGSERVIRQPNHVAIQDGNVSETLQQL
ARRLCQELVSKTKGHKCNPCSSLKYHQGNCYLLSHRNRTWEDSRTYCASKNYVMLKVDNK
EELAYINRNTNKIRWIGLSRRAIDSPWMWEDGSVLATDLFQISGDGEANRHCAFFHNGKI
QAADCQESYPTLFPSLAKEGFKACWPSSSDWELHVGSDYLDLHSAWHTARTKQNTTVIIE
KQRGSVERARAWESEPRSTMQDEDGYIMLDFKSRIHATSKGPSGAPGLPPSWRWMTLALL
ILCLGMLIGLTVLGSMCFWGTSGPKGPHKSSWGLGASDVPIFQKETLDRFIVAADLKVAN
IHSPASPTDSRQQALLQQITQKYCQELSSKPGGHKCSSCDHNWRFHGGKCYGSFKNNKTW
EESKKYCDDRNSTLLKIDTQEAWELNSLSVSSLIFLVPTLCRRLSWILGARKEENFIQGR
PDFTRWIGLSRPSSGGRWTWMDHSALTDNLFELLGDGDEGKHCASIRKKKISTTFCRELH
YYICEKVGIVKVDQLV

>CLEC_like5
MSVEDVYILKELAIIVMINNIYSSNSTESSPTLVLGTWWRLLTVMSWLFCLPLLASTVIL
SLKVIQASELIGKQEEILANLSLQQRVCSERLQVCQVQIQMSTSPESNCSLCLEPWVMNG
ESCYLFFDGWKNWASSSEFCVQEKSELLKIGSKEELHLVQGSAQSAPGLPPSWRWMTLAL
LILCLGMLIGLTVLGSMCFWGTSGPKGPHKSSWGLGASDVPIFQKETLDRFIVAADLKVA
NIHSPASPTDSRQQALLQQITQKYCQELSSKPGGHKCSSCDHNWRFHGGKCYGSFKNNKT

WEESKKYCDDRNSTLLKIGHSRSLGAEFLVSVIINISGAYAVQKALLDLGSTQRRGRPDF
TRWIGLSRPSSGGQWTWMDHSALTDNLFELLGDGDEGKHCASIRKKKISTTFCRELHYYI
CEKLPATPPLRPFADSAILPEPPASLIVPDLHLGRQCGYLEAMQDQGTYDSLYWVTPDPP
PVTRPSLKTALRTVLGTYSSNSTESSPTLVLGTWWRLLTVMSWLFCLPLLASTVILSLKV
IQASELIGKQEEILANLSLQQRVCSERLQVCQVQIQMSTSPESNCSLCLEPWVMNGESCY
LFFDGWKNWASSSEFCVQEKSELLKIGSKEELEIKTASPKWDKDCVQPDLLVSTLVLSTV
PGTQTNPQIDFSHWLLRWMFSSCFKSFGTIMAYWREHGPGSQKDWVLIPKSSTCLLCDLG
PTALMYIAVMYLFAVMSVSSCKLILGRAFINRNIERKKTGSSWSYWVGLTQDKCYGDWRW
RDSTVPSSDL

>CLEC_like6
MGMKIVSLTWDNLITLYLPQRLEQCSAHRQSCLDSSMEDEDGYTILSPRTRVFAREPAAS
GKALNHLAPSYLASLLSYRNPAHRYRSSDDNLFTYFDLVHLPSNLSPMSCLWPGTPSLFM
SESLPAVSPRWRLAAVTLGIVCLGLLGAIGILVFQALRPVNGPCSSEMSKNHIQPSKSPP
VRKRDSTKAPETITMVPEPRSRICPPNWNRNGDSCYLFRYTLDNWNRSKMFCESQRSHLL
RINSHEELVFIQHLTSNNSQMSVWIDLTSRSDGSWMWGDGSVFSPHLFDIRQTNSLNRCA
WIHENLVFDALCSSLAYSICEEKTVNSLWAGTASTNSVRPYSPKLLVSSSAGRQAEGNGG
DVPLPRSGKPVHRSVHRSPVQMRAKYSSTQDMLEDDGHTTLSLHSRTSSAARSPEPSGPD
SIPSFLQPSDGGLEKVKDEKKVEEEKEEAEAEKEEKDEKEEEGEDEEEEDEEKEETGAGE
GRSGRGGGRGGAGGDTAVKEHCSHLGRVAPSRAWRPVALTLLIVCLVLLLGLGILGFEFF
QVSRLSNAQRTAISQQEERLGNLSWQLQDLRAQTRKLTGTLQQVAGRHPGGDILEAGGDA
SLKGGGEDRGRDVDLRVICVEMVVEAVRANEFTKGELVIPLSFSKFFYSYWTGLSRNGSG
QPWVWLDGSPQASDLFQVVVDSDSPRSRDCVTILNGKTFSKDCKELRRCACQRAAELDLG
NLSLILPPVCCVTLLCITLACGFVPFIHPTLSPIADEGTEKFSDLPKVTQQTFGKDGIRT
HDL

>CLEC_like7
MLLIQCSTQSECSVNTSDSSGNSGFQRHKRHKPFAAKLGIDLKCTMARGANGEFNEEDCR
GLRSNHMVPLIMMFPLKRFLPPGGGSVVKEPENQVELRNACGQGQCPFGPYAFPTFLHPI
SALIRLTFVCKGHSQGLFYKRKHAQRSRRERGGPGTDLRGQEANPVADTAPPSFLSHSPE

NLALSSTRRFPSTHIFDPSPNPVSSTFVTSPNSDLSSPSDRYRVNTVTHPNLPREAAGLG
GTSPGLGVRDARGSDDAAFGNDVALENASTGARGVGMGKMSPDGSVKLFLSPEDCGFVSD
SKSQREQRAEECIYYTLDHKCLLYAKHRSKRWGTHTVIGLSHVGLTIFIPILQLRYFAVD
ASGSAKQGDEWGYVSRGTADVLGGRRLVFSHIVIIIISVKHLRRANRHTKRRRSYESSPH
MGSPSKDGENSYFPDEEARAQRSDLPKIAGQAYGRTGIRPGPALFDYILLPPSVGQFPPY
ASPLKPLGDRQLAPCPL

>CLEC_like8
MEEALSTVLCTRGGKVWEKTAFGKEVEKFNFGYAEFEVMMGYPGLLGGTRSSLGAGNSKT
IMAVGFYNFANFLWRYKRKLSTNVAEGVCVTQLPKMNQRKSVDLFASIPLILLVCISAGV
SILACAYPLDMEDDVQYAELKFNMPEEKPKQKLPEKKAEDSSSSSPRWFLAAIMLGIFSF
ALLGAMVVLGLKITSSVKWGLTVSLTWDNLITLYLPQCLEQFSAHIIHARGLRDRQVENY
THLEGKTARFTDQKDVAGASPQDWQEVENLRRKLENITEEHRALLLQNEQLREALKKAKN
YTGPCPQDWFWHGESCYIFSSHFKNWKMSQENCASQGSQLLKVDDQDELEFVYRAVAHSR
NPFWLGLRRSDARSRWVWEDGSTPFVGL

>CLEC_like9
MVPDQAYQDRVGGETGLVPSMAPEIVYAELRFKMDNEKLGAPPAPAPQKATPPPLRPRLP
ILQMALLLLLLLLLLLSFLVAFVVFYRRSHLCQEDRSLKSTRILSEFVCVKQSSQSPVFS
CSYIETHQETEDLPSPHQLPQNQTTLDCINEGSEMEGRSWDCCSRGWRPFQSRCYFISAD
KMSWDQSQQNCHQMGAHLVVISSDTEQKFLESILDNKAAYFLGLTYLEVSKDWRWVDQTP
YNKSVLVCVQPEYLVPAPTLGTSSQITPVIIHTYSVTTKEKYISNF

>CLEC_like10
MHHVGEASQTPVIPKPGVTVFEFGKNYGHSIEPNPSIPVQGPSPYPSSLKTQIVSEQSIS
TTPHTTTTIINNTITNHNPIPHMLRQGSSDLSACPPVTQHGFAQLCKDPEGLAPLNHTQV
SCLRERSQVEGRMWSCCPQDWRPFQTSCYFFPRDVLPWNEAQSKCLKQQAHLVVINTEAE
KNFITQGKPLNSSIHLGLKKRKKERQWRWEDKTPYNPADTFGVEGDPQEGDPALGAREER
CAVLTLRQTSSRWRWSGTDCVDATAHWICEMPRRSF

>CLEC_like11

MSPPKGAGPQNRNFNGHFPSRLSGLLGRGCCCPPWSPWAAATVPVLLLSACFIARCLVTQ

RAFSQFCEEEGTLLRLNDFFSEISCYSSGSGSIRDCCPRGWKHFQSHCYFFSSDTLTWSS

SRLNCTGMGAQLVVINSQEEQEFIFQSKPRRREFFLGLTDKHVEGKWTWVDGTPYDPSRS

FWDLGEPNNIVGVEDCATTRDSPNAKETWNDVTCFSFHYRICEMPAQTLSAGKKGM

>C3AR1

MGTCWGHRMGRMSMPWDNTSSPDSPLQRGHSLPQLLSMVILSFTLVLGLLGNGLVLWVAG

WKMQRTVTTIWFLHLTAADLLCCLSLPFSLAHLALGGHWPHGQLLCKLVPATIILNMFAS

VFLLTAISLDRCLLVTRPVWCQNHRGVRVATAVCGAAWVLASVLCLPVLLYRETYVDGPL

WRCGYIFDHHASKDGLEDNDLLGFSNSSPPTPEMDDPGTLDIPLRARDALLTGSVWLLGP

SLPWGPLDTEAGHPSAIPNPDLSLDPGGPRGDLLPTGMPSQSPDELDLVMDSWDELIKIY

TAQSQVPGPLVAMTLTRLSLGFLVPLAIMVACYSVTVIRVQGSRFAVGQTRTFWLAARVV

AAFFACWAPYHLVGVLSLLATPNSPLDHALATWDPLTHALASANSCINPLLYALAARDFR

VRARLSLRSILEAAFSEGLSHFSTCPPSQSDVLSIDPLGPEV

>NECAP

MAAEAEYESILCVKPDISVYRIPPRASNRAYRASDWKLDQPDWTGRLRITSKGKVAYIKL

EDKVSGELFAQAPVDQYPGIAVETVTDSSRYFVIRIQDGTGRSAFIGIGFSDRGDAFDFN

VSLQDHFKWVKQESEISREAEKPDTHPKLDLGFKEGETIKLSIGNITTKKGGPAKPRPVG

AGGLSLLPPPPGGKISIPPPASSVAICNHVTPPPVQTSSQGGGEAGTGGLSRGRRVSGPQ

SVDPSPGALTSLSCRPPTHGHSDILLDLDAPAPSTKAPAAIPAAPDLWGGFSTAARPSPV

LIQQGDPPFNTTTCSVQAGSPKVPIANGFLAQGPDDTQIYISAPVLSPSLQARISSCLWD

VSTWMSARHLKLNMSKTELLIFPPKPGPLPDFSITVDGTTILPVPQARNLGVILDSSLSF

TPHILSVTKTYRFHLYNIAKIRPFLSTQTATLLLRALVISRLDYCVSLLSDLPSSSLAPL

RSILHSAARLIFPQKRSGHVTPLLKQLQWLPIDLRSKQKLLTLGFKALHHLAPSYLSSLL

SFYRPPRTLRSSAAHLLAVPRSRPSRRRPPGHVLPRSRNALPPHLRQTDSLSLFKTLLKN

HLLQEAFPD

>BIRC2

MGMKTVSPTWDHLMTPDLPSAQASAPHIVSAEQIPTLLFFFFSGPQFPPLKNGDEDCEPH

EGGVGNGKRRRSCGHVEGRKFWRRKFGRSPVPGLQPAGAAAAAVAAGKPSGAEERPMDPG

QQPAPAGGQPPAPPGTPASAAAANAAAGTPEPAGPGGGGGPGGAGPGGGGGPGGAGPGGG

GGPVAAPAPVAAAAAAAPPPPAGHQIVHVRGDSETDLEALFNAVMNPKTANVPHTVPMRL

RKLPDSFFKPPDPKAHSRQLGAVSPGTLTPSGVLSGPGASPSAQHLRQSSFEIPDDVPLP

AGWEMAKTSSGQRYFLKEHKAVDLNRCAKAKCFKIKSHNSGTKNEKLVMVWLDGVAHPTS

LRLGTCCQVLQLGGEDLFSGGHDGSEENGDPATPTLSTNLLALVFGSSFHQHPQSDNGLT

EAGFCDGVQFRVADKDLWFISTSETSECKKFSDKLSTRDNKHQSHRSDHNVAGPKEGHVV

PDERDGPDEPSSATEYNEFRIRAQKHYRLALSKRVANAIVIIKQSPLCFLSASMNQRISQ

SAPVKQPPPLTPQSPQGGIMGGGSSNQQQMRLQQLQMEKERLRLKHQELLRQALRNINP

STANSPKRQELALRSQLPTMEQDGGAQNPVSSPGMSQELRTMTTNSSDPFLNSCLICKMG

IKTVSPTWDNLISLDLPQRLEQCLAHIKNVDARSVSAVIFKHPVSTHPPPICCSSGTYHS

RDESTDSGLSMSSYSVPRTPDDFLNSVDEMDTGDSISQSNMPSQQNRFPDYLEAIPGTNV

DLGTLEGDGMNIEGEELMPSLQEALSSDILNDMESVLAATKLDKESFLTWLYCTFPSAQY

SALHTELTEDLKILVTLLATPKSGLFAPNAEEEAQHFHVKPKPGEEEGAVTRKEVTSSGK

WGLTVSLTWDDRMTLYLPQRLDQCSAPMTSSVKWGLTVSLTWDNLMTLDLPQRLEQCSAH

CKRLTKTNMTILWASVPSSVKWGLTVNLRWDNLMTLDLPLRLEQRSAQGSPGPARLLKEP

PLPALRCPGLDQSFRRPAPGRVLSTRKALDTYRWVDGSGEAADWEPERLAIPLSALLWVA

QLAREIDAPPQSCWEKPVSPAGAGDQAARRARCLLQATGRPGNKQTELTPRVLAWSCLKL

QTVHKTTSRGETRPVSPATPTEYFVMNVVANSMFLSNLMNGNSGYELKYDFSCELFRMST

YSTFPTNVPVSERSLARAGFYYTGASDKVKCFSCGLMLDNWKPGDSAIEKHKQLYPSCSF

IQNLLQTHNPGASSYSAFCPPPLGSLSPTATISPSLEPSGYFSGSFSSFPLDPVTSRAIE

DLSQPRPHVDNSAMSSEEARICTFQSWPLTFLSPSALAKAGFYYTGPGDRVACFTCGGKL

SNWEPKDDAMSEHRRHFPGCPFLERQTRDASRFNVSNASMQTHAARVKTFLNWPARIPVQ

PEQLASAGFYYVGRNDDVKCFCCDGGLRCWESGDDPWIEHAKWFPRARKEEMNGEPGYPY

CDPADVASWHELSRSDRQIRSVSSTVTYRLCEYMIRMKGQEFVNQIQARYPHLLEQLLST

SDTPVDESADPPIIHFGPGENPAEDAIMMSNPVVKAALEMGFSRRLVKQTVQSKILTTGE

NYKTVNDLVSDLLNAEDETREEEKERQNEEMASEGLHYRIGAGVAIPSTMEACHRARGFL

GGSSLERHQKLEEQNSPLPTAFAFFSLVAFGLLSAGVISEQERDVIKQKTQTSLQARELI

DTVLVKGNAAANIFKNCLKEDDCILYKNLFVEKNMKYIPTEDVSGKNKKAPINTKELE

>CARD9

MSDYENDEECWNTLEGFRVKLISIIDPSRITPYLRQCKVINPDDEEQVLNDPSLVIRKRKV

GVLLDILQRTGHKGYVAFLESLELYYPHLYKKITGKEPTRVFSMIIDASGESGLTQLLMNE

VLKLQKKVQELNLLLSSKDDFIKETRVKNSMLRKHQERAEKMKEECQAFSQELKKCKDENY

NLAMSFAKQSEEKNAALMKNRDLQLEIAQLKHNLMKAEDDCKVERKHTTKLKHAIEQRPSQ

EVMWEMQQEKDLLLVKIQELENSIREGKREKNSLYITVLEEDWRQSLLEHQEKETTIFRLR

KDLRQAETLRNKPGMLLIGREALSKSAPFPTVRGGGEPAPGSLVRPSGPEAEDEGGAGAEL

SQGGAARSIVPAVMFIERLLCVEHLVKHGQGKGQEHLDNDPPCMEEKEMFELQCTTLRKDS

KMYKDRIEAILQQMEEVAIERDQAIMTREQFHLQYSQSLIEKDGHRKQIRELGEKCDELQL

QLFTREGQLLSMESKLKRLQLETPNLSLDLEEMSPRNSQEDKTLTYPTAPHSVQSERPSHN

LDRGDLQPAAASPVPLHRRLSDRSETCAPIVGRSGEGPAFSFPKGKQGQCTARGLNTEFKV

LHTKGKVSNGEPPDKERRRMKDNFEHYRRKRALRKNQTSRHLGEVDRDNTTGSDNTDTEGS

>SNAPC4

FEGTRNAVELRKFWQNHEHPSINKAEWGEKEIEKLKEVAAKHSYLNWQKIAEELGTNRSP

FQCLQKYQTYNKDFKRKEWTKEEDQMLTQLVGEMRVGSHIPYKKIAYYMEGRDSMQLIYR

WTKSLDPNLKKGFWTPEEDAKLLRAVAKYGERDWFKIREEVPGRSDVQCRDRYLKRLHVN

LKKGKWSPSEEEKLIELVEKHGVGHWTKIAAELPQRTGSQCLSKWKIMIGDKKKQRNRKR

QRKSRRGPRGGQHVPSSSEESEIEFTDNSEEEEEKKSDQEKEPAVPSYTVPNIDLWIPSG

EPRATPRGAVTIQTPRPRARTMGSGSGGATPGPTGDAGEGQPRGQGASMDVSATPKGFPC

ARSTDIPLENPEPQGNPEPLENPEPLEKEDPGSGKQVLKLTPDEVKKVLRSNTYILRRKL

QEKLKKPRLPTSSVMPQVSAGMSGDQVLQVWENTVEKRLQRLKVGLRRNIRRSNLDRRLL

LAVTPWVGNVTLPRFQNSRKGLVLQTKVKKRPLQTVLLTSTPMFTLFIQLFQIDADGCMK

VIQERKANQVEVVQVEPGNPEQLRQASCSSQDTSGCQPQKDPAKGNSRRRSAPRVEDGQL

ARDGKVTAQAGAPSAPQPQRPKPKTVSQLLREKRLRESKATMPNPVILAPQVLVSQPVLL

RHPLQPVIPSGPATATLAQAGPPVKPPAPSPTSTPISSASLRALASGTLAVASENSSGQV

SSEASRAVGVTEDPKAKEEGKSSAPLTVVIPPPGDGRGRVQDGGSPQPGVPPNPSPGPGQ

LLAISAPGGFTLARGAPSGGKQGPPEASPLLPPAPVRFQQPRLLPALAVRPSGPQGGPSE

ILPFTWVVTPQGLVPMSVQAFVGLHGPSKTVGDLGAANCQGPVGTTLVPPGGPTLRLLLP

PATTPLAVTSSGAPGTNTAVAPGSSLSSHRVAGEPAPPTVELSHSASLPSAAPPDREVSR
SPEEVSAAEKSQPPQPTPSPQSQNPNGGQRAEPLGSHGLPASATKSPSSGEGEAPSSQSH
MPKDLPATQVGRCPDRTGASGNSAPTTAPLPARVGKDRPIAPRPEATPSPGGHPAPADPP
PRGHPPPGGDRPPGPVAPDKNKLDLTLISLEEEAVVREWMKGKRGVHVPPLKNNLPYLPP
SVYNLKTLSGLLREKKNLERSAASLVAGDGGPAGSELSPEGLEAIRALVRERLKDNPAYL
LLKARFLAAFTLPAFLATLPPPGVPTTLSRRPEESEEEDEDDGEEEEKDLTDEESGEEDH
PPGEAQPGPEAATSGQEAKKTEVKGRLGVSWSGSEVTAAKKRNGGAPGGRGPERDRCSGE
PSEMPGLRFLPYREDGEPVSCLHLHLRVIFVVFPMTDPEETLMDLLGTEMKPSPPVVNCE
FGFPVRRHPIKEPPCRSGPGYNEVKSGPSGSALKQKLPQQAPPSLEFLHRLLHFLSTPRA
KDSPDTLKCACKAESGQRWWLPVGLKGHRGMLSFPEALRINIRARWRRPLQSILNRLCKT
LDFFVRSRHRTPCPGPASVGCHLKRSPSQEVHWAPV

>ARAF
MSARSGPKGAGPAPAAPPLSLLPPPPAGSSPQNRSQDGGDGGAVTDSAPVEEDQASGGGM
GSEGFGQRSCLLRHVARDSRWRQTHHSRNAAVGKGSTPQPSVSPPGAPLMKRSLGTASLI
SNGDHLASPEPPMSPAPGPMEPARRPAPDGGEAGRTGGTVKVYLPNKQRTVVTVRPGMSV
YDSLDKALKVRGLNQDCCVVYRLIRGRKTVTDWGTAIAPLDGEELIVEVLEDVPLTMHNF
VRKTFFNLAYCDFCHKFLFNGFRCQTCGYKFHQHCSSKVPTVCVDMATARHSIHTYASVQ
DLSGTGVQEPLSSQSLREPLTPQPVSSQQHLSPEAFSFPEAPGQALQRIRSTSTPNVHMV
STTGPVDSSLIKRAARNFNAEGTGNGEEGHPQGTGGGSSGPPQVSPGRKYSHPKSPSEPR
ERKASADDKKKVKALGYRDSSYYWEVPKSEVQLLKRIGAGSFGTVFKGKWHGDVAVKILK
VTSPTSEQVQAFKNEMQVLRKTRHVNILLFMGFMTRPKFAIITQWCEGSSLYHHLHVAET
RFDMVQLIDVARQTAQGMDYLHAKNIIHRDLKSNSIHPTRPQEMRDIFLHEGLTVKIGDF
GLATVKTRWSGAQQVEQPSGSVLWMAPEVIRMQDSNPYSFQSDVYAYGVVLYELMTGSLP
YSHINNRDQIIFMVGRGYLSPDLSKISSNCPKAMRRLLPDCLKFKREERPLFPQILAAIE
QLQRALPKIERSASEPSLHRAQADELPPFLFSTLRLVP

>RAF1
MTGPVKGPIEFRECGFRTGGEYVRQQPGKSPGDPPAVPEFDEKEVIGGGGTANIPCFSVV
RDLLDRGCLCSAPGLKPGYGIRDRPSTDGPPDRMRLVGRASWVCAWGARRCGCEDLKGYR

SYARAVAGLQERGGCAQVSGKMFLSGPAEPSRGGLRWKTRHVNILLFMGYMTKDNLAIVT
QWCEGSSLYKHLHVQETKFQMFQLIDIARQTAQGMEQHQLSGCWKGLWVLRGPLGPVKYI
GLGLSFIGRGSHIGEETPRGMPSASSWPNGTALVEGFWDLEFLALEERKGLGDWALSVTP
RERCLNRALLIRHSSLTGPPRHISSRRLDGEDRRLRSGNGEIAVERLSAGGAAYGIDPLD
DEGTDAKVSAVPGPVPGPALWAPVSSLRSSCLENRLTPVPLSFCPVKLAYGSSVDMASVV
VAEQYRISLWSTIQLDRETLGLGELTVALVVVGLELLVYVHHVHPVEKRHGLAARARAWE
SEVLPENVAAAMRLALWDGRRRPGGQIGQPPGARGWIQGFPGISPRHICIEHCTKRWDSE
PHGELVSDGVHVVKNQSRLEQRLARSQRSTNPTVTVIFSSVPVGKRAPEVIRMQDNNPFS
FQSDVYSYGIVLYELMTGELPYSHINNRDQIIFMVGRGYASPDLSKLYKNCPKAMKRLVA
DCVKKVKEERPLFPQILSSIELLQHSLPKINRSASEPSLHRAAHTEDINSCTLTAATRLP
VF

b) Echidna gene sequences

>PYCARD
MGSARDHILAALENLTSDEFKKFKTKLLSCPLREGFGRIPRGPLLNMDVIDLTDKIVTTY
MEAYGLELTTAVLRDVNQAEATALQMAAGPATPGPVLSGPAFGEAGQQHFVDKHRQALIN
RVTSVDAILDALYGKVLSEEQYQAIRAEKPSTKQMRQLFEFSLSWDRSCKDQLFQALKVI
HPFLVKDLENS

>ITGAX
MGERLFTCAECGKSLGQASTLAHQCHSHTGECPAPAPSAASASTRSSTSTRKLVSSPTTA
SFSRSSDLFQHAQAWRGDWPYPCPNCCHRFGQSSRCLDRQRLRRSERPFSCSRGAWGSLF
SEKIPNCHTYLRKDGTTILPISQAHNLGVILNSALSFTPHIQSVTKTCRSHLCNIANIRP
FLSIQTTTLLRLEQCFAHSKRLINAIIIIIITGEVTSSLIYKMGIEPVSPMWDRDYIQFN
LLVSNSICLHLPQRLVQCLAYTLIPCLGFNLDSENPTIFQMDRDGFGQSVVQFDGARLVV
GAPLEKGSSNQTGQLYECEFGAGTKECKQLPLQIPSDAVNMSLGMTLAASTTPSQLLACG
PTVHQACGENLHLNGFCFLLGSNLQLTRRIPPALQDCPKQESDIVFLIDGSGSINSGDFD
RMKQFVRAVMRRFQGTNTLFSLMQFSNSFEIHFTFNEFKRHPNPEMLLYPISQLTGLTYT
ATGIKKVVYVSSSVGTAATEKLFQQSQGARRDATKILIVITDGQKYEDPLEYSDAIPLAE

AAGVIRYAIGCLEQCFAHSPNFPVPFGQVGTAFKKYEAREELDTIASDPPQDHVFQVNNF

GALDNIQSQLQEKIFAIEGTQTGSSSSFQQEMSQEGFSALLTPDGPVLGAVGSFDWSGGV

SLYPPSGGSIFINISQENGDMNDAYLGYSADVATQRRIRGLVLGAPRYQHTGKVVIFRKI

AGSWNQWAEVTGSQIGSYFGATVCAVDVNRDSSTDLVLIGAPHHYTQNSGGLVSVCPMPL

WREDWWCKSFLRGQPGQPLARFGAALAVLGDTNGDGLADVAVGAPGEEENRGAVYLFHGA

APAQISPTYSQRIGGSQLSPGLRYFGQALSGGQDLTQDGLVDVAVGARGQVLLFRSRPVL

RVDVSLRFMPSELARAVFDCGEQTVPHREVGEARVCLTVRKTSRDRLGPSDIRSSVTYDL

ALDPGRLSSRAVFDQTKTRTLRSERSIGLGKSCEAVKLLLPECVEESVTPIVLRLNFSLL

GQPIPSSGQLRPVLAEGSQDLFTASLPFEKNCGADSICQDDLSVTFSFSSQQSLVVGTSP

DLNVTVTVRNEGEDSYGTLVTFSYPPGLSHRRVSVLQNPRQPRALRLTCESARPRSSSCS

ISHPIFREGAEVTFLVTFDVPPTADFGSTLLLSANATSDNNTPKTSRAFFQHQLPVKYGV

NVLVNSGEQSTSYLNFSASEEKSGHVLQHQYHVNNLGLRSLSITVAFWIPVELDGLPVWM

APEVITPQDFPVQCVSDTENPRDPGFRALLQQSPDLNCSVAVCRRIRCSIASFQVREELT

FTLKGNLSFDWVGQMKQKRLSVLSSAALLFDTGRYAQFPGQEGFLTAQVKTVVERYEVYD

PLPLIVGSSIGGLVLLALLTAGLYKLGFFNRQYKDLMAEGAPTAGETPDMPHPKHLLDCE

PVVGCEALGNLGQDGPMEIPDKGGHSYFIYTMWIKTESLCDTNHV

>TRIM72

MSAAPALMQGMYQELSCPLCLKLFECPVTAECGHSFCRSCLARLPQDPQAGGTPCPSCQA

PTRPEGLSTNQQLARLVESLAQVPQGHCEEHLDPLSVYCEQDRVLICGVCASLGKHRGHS

VVTASEAHQRMKKQLPQQRLQLQEACMRKEKSVALLDRQLTEVEETVRQFQKAVGEQLGV

MRTFLSVLEKNLGQEASRVTGEAGAALQGERRGLASYLEQLRQMEKVLDEVTDQPQTEFL

RKYCLVTSSPAPSLTIVHTSHKFLGPTPSSNRFHTHLGFPSRLQKILAESPPAARLDIQL

PIISDDFKFQVWRKMFRALMPALEELTFDPATAHPSLVVSPSGLRVECMEQKAPPPGDDP

QQFDKVMGVVSHQLLSEGEHYWEGFWLLGFRDGKVFEAHVESKEPKLLKVEGRPTRIGIY

LSFQDGVLSFHDASDPDNLTPLFSFRERLPGPVYPFFDVCWHDKGKNAQPLVLVRQEE

>FUS

MGVIRSSPPKTISRGLYGQTVGVPLPVLEGPTSGSVDVFRLQLWTSLSLDWGCGEFDFAF

ILLQSTPSQPHRAMDLIPPSLGRATRSRAVSLMASRVTVATANRLTLRAMARAATVPPTD
RRRTARGQLTGWVFAYRQCSPNQCSPNQLALPHPLSSMAWSQKDLDSNPGSASLLLGDFG
TPSTPQGYGSSGGYGSSQSSQSSYGQQSSYPGYGQPSASSSSSGGYGSSSQSSSSSYGQP
QSGGYGQQSGYGGQQQQQQSSYGQQQSSYNPPQGYGQQSQYNSSSGGGSSSGGSGGGNYG
QDQSSMSGGGGGGGYGSQDQSGGGGGYGGGQQDRGGRGRGGSGGGGGYNRSSGGYEPRGR
GGGRGGRGGMGGSDRGGFNKFGGPRDQGSRHDSEQDNSDNNTIFVQGLGENVTIESVADY
FKQIGIIKTNKKTGQPMINLYTDRETGKLKGEATVSFDDPPSAKAAIDWFDGKEFSGNHI
KVSFATRRADFNRGGGNGRGGRGRGGPMGRGGYGGGGGGGGSRGGFPSGGGGGGGQQRAG
DWKCPNPTCENMNFSWRNECNQCKAPKPDGPGGGPGGSHMGGNFGEERRGGRGGYDRGGY
RGRGGDRGGFRGRGGGDRGGFGPGKMDSRGDHRQDRRERPY

>PRSS36

MFPIHLLLLLVFSCDPGDTQVPDCGIPHPSARIVGGSESRPGAWPWQVSLHHDEGHICGG
SLIADSWVLSAAHCVVDNGTMSEAEGWSAHLGLWSQDTRRSYEQHRAVVTILIPENYTSV
EFGEDIALLRLATPANITDFVRTVCLPRATHRFPSGATCWATGWGDLQEEDPLPPSWGLQ
QVELRLLGGTACQCFYSQPGPFNLTFQLLPGMLCAGYREGRRDTCQGDSGGPLVCEEGGH
WVQAGITSFGFGCGRRNRPGVFTAVARYEAWIREKTRATFLSQPEPPPSEPPEEKNCTLA
LPVCGQAPHPGTWPWRSEVVVPGASKPCHGVLVSDKWVLAPASCFLGSDTLSPNISGWRV
LLPPGVRSVPITHFVPNENFTQDADYDLALLKLEAPVNLSEDIQPLCLPHADHYFLPSSR
CRLAPWGRGGSPVGPHSVLEAELLSRWWCHCLHGQQVETAAPGLLCPLYREDSHGCWNAS
RWSLLCREAGTWFFAGAGRAPEGCARPKVFSALQSHGVWITRVTHGAYLEDQLSWDWNLK
EEEMTNQTCPHSTAQWACGLRPGPEGAKGPWPWIAEVHVAGGQLCGGTLVAPGWVLAAAH
CGLRTGATPPSLGHEISRKVTRVHLPTGSSSQPSFALLELGGPVEPSPSALPVCLHSGTI
PAGARCRVLGWKDPRDRGERGSHADERSRKPVVRVVGRAQEFRGDVGAGTINNMVDIAGG
WFMAENAAGSGEGETTRSKVTNDLLAKTNGLYSILILLDLSAAFDTGPR

>NLRP-like_1

MGHIELRKLDEVERSEISVGEQHRFVGRRFVGKKAGREVAVQPYWIFLTQLSLDIKECLV
QCSAHSKRSINTIDDDDDDEGPAIRERYQAQMGEKFRILRDRNSRPGESEALRHRFTRLL

CLPEHRQREEKEHELLATGVDHARAMEIQGQFVEVNALFDPDQERRFQPKTVVLQGAAGI

GKTVLARKVMLDWAEGNFYEDAFHYVFYLNCREMNILSERSLADLISVHLGDSQVPFDKI

MSQPGKLLFIVDGFDELKWVFEEQESDLCHDWRESRPVPVLVSSLLRKILLPEAYLLITS

RLTALDTLNRLLLHPRHVEILGFSAADRKEYFRKYFRDENQATQAFSLVEDNETLFTMCF

VPLVCWIVCTCLKQQLKRGEDLTQTSRTTTALYVSYLTALFPASQDHPGQDHPPGPILRR

LCHLAAEGLWTRKILFDGDDLRKHGLYVSDVSPFLQMSIFQKDIDCENCYSFIHLSIQEF

FAAMFYTLGPEEGRMNCPDPDFGDVKKLLEKYRKSHETGFLTLTVRFLFGLLNEERVKML

ENKFNCKISQEIKRELLHWSQNQCEFLQQGTLFECLYEIQEEEFVAKVMDHFQEVDVCFF

TKMTFLASSFCLKQCQNLKKIKITAVYEEESRQATEPQTDDPENKSSFNFTCWTNLLSVL

STNQSLTELSLRFPQIDDLTMLVLCDGLKQPNCRLQKLINQNLIHLDLTGNNLGDGGAKL

VFEALRHANCHLQSLRLEDCSLTAACCRDLCSALTSNRNLIYLHLGRNELGDTGVTALCE

ALKTPSCSLRSLWMQDCGLTDACCPNLASVLRCNQNLVQLKLWGNDLKDSGVDMLSGSLR

APQCRLQELGLGNCKLTDACCERLSSALGRNRTLIHLDLQCNFLTESGLIPSLLLPVVQK

ATGPMMYQLQNNYSKTPEPASLQDGQSGQSLQLPKEEVHTYSHQNKDLFRKMFPHTVNKR

QRNQSYRHHQERNFL

>NLRP-like_2

MGMKTVSPTWDNLITLEPPQCLEQCFAHTSLVVVGSAGLLGGTAQTLDIGQTPTLPSDGS

EVSPLMGPRGALHFVDRYRGQLIGRVTSVKPLLDLLHGKVLSEEQYRTVLARATSFDQMR

ELFAYLPSWDVDGKELLYQAIWQIHPQLVAELESSLICKMGMKTVSPTWDNLINLYPPQR

LEQCFAHIWLPSPTFHGNCPLKGHHDLLLAKSNGSYSVLILLDLSAAFDTVDHPLLLNTL

SDLGFTDSVLSSFSSYLSGRSFSVSFAGSSSPSHPLMVGVPQGSVLGPLLFSIYTHSLGD

LIRSHGFNYHLYADDTQIYMSAPALSPSLQARISSCLQDISIWMSARHLKLNMSKTELLV

FPPKPCPLPDFPITVDSTTILPISQARKLGVVLNSALSFTPHIQAVTKTSRSQLRNIAKI

RPFLSIQTATLLVQALILSRLDYCISLLSDLLSSQPSEGVSLVEDYREKYLKHVKWKFQY

LEERNARLGEKVALGLRYTPLLLVEEHRNLPQRQHELLALGRPPTSRVPRRVRVDALFDP

DAEGLEPPLTVVLQGAAGIGKTVLARKVMLDWAAGTLYPGRFDFAFYVHCRELNLKRKRS

AVQLIQQCCNDDSIPLSKISQRPDRLLFLVDGFDELGWSSGGWAAEKDDDDDDDDDLFVD

WRDEKPVGSLLAHLIRKRLFPKASLIITTRPAAAEGLRPLLRWPRRAEILGFSEAERAEY

FHRYFPDPGQAERALAFIQENDVLFTMCFVPLVCWIVCTGLRQQMERGEDLAQASKTTTA

VYLAFLSSLLRPARHLPSRVPPAHLRGLCSLAADGILSQRVLFREADLRKHGLPEDGVSA

FLHVDVFEKDADCETLFSFVHLTFQEFFAALFYLLGPDEARPGAGPGPVPDVRTLLRHYR

LCDTGFLTLTVRFLFGLLNRERVADLKEKMDCVVCPEARPALVEWINASAQKEVLPGPTL

QWLYCFYEIQEVDFVERAMGSFRKIDVDVRTRMDQIAVAFCLRNSRNLCSIELMRFFNVQ

EEEAAVAAAAAAAGTPKAAAYQGSQDWLQDTFCESLSEASAKNRDLKHLALKNKTLRGRG

AELLRKGLTHPNCKLKSLRLVNCVLPPDSCRDFSSIVSTNQNLSELDLSSNALEDSGLSL

LCEGLTHPSCKLQTLWLQSCALTSACGPALCSLLSTNPNLTKLDLFNNALGDAGVRLICE

GLKNPNCKLQALLLRHCAVTLNCCPDLSSVLSINRHLSELDLSYNSLEDAGVCLLCEGLR

HPNCKLQTLRLQRCKLTSRSCPDLSQMLGVNRELREITLHDNVLGDAGISEIWEGIQQPS

CGLKILRLGKTDLSEKMKKEMKTLQNVKAELKIRYWFRGSQGGRGLLVRSPSPEHVTLAL

SGEG

>NLRP-like_3

MTCSIRSRLAEYLEELRDPELRKFKFHLEDLAPAAGWAPIPWGRTEKADSLDLAHLLVAH

VGERGAWELAVHVFERIHRKDLWERARAEAVVRDLSVSALGCQWEQRRRAEHLVEPHEAE

ATRDPRDVYREHIRRKFRFIEDRNARPGECVNLSQRYTQLLLMERHPTPQDAAPESPTAR

LEAAGAPERQPGPVQVETLLEPDASWPEPPRTVVLQGAAGIGKSMLARKIMLDWAEGRLY

RARFDYLFYISCREMSRVGRSSLAGLISGRWPRREAPLADIVRRPERLLFVIDGFDELGR

SPRRPPPGRRAGGWKAKLPVASLLGGLLGKDLLPEASLLITTRPAGLARLQPLLEQPRHA

EILGFSRAGRRDYFHKFFGDGRRAARALGLVRDVEALSAVCFVPLVCWIVCTCLKQEMER

GEPPGQPSKTTTSVYVFYLLSLLQPDPGGSGPRGRPVLAPLCSLAAHGVWARKVLFDEGD

LRRHGLAGSDVSAFLNLSVFQKDIHCQRLYSFIHRSFQEFFAALFYLTGGDGREGGRGSR

HSVTRLLEHYGRSETSYLALTVRFLFGLLNEENKSYLERQFGCPLSPTVKGELLAWIEAR

AQIGGRTLERGDAGPLLLLTQEEGFIRRALDPIQVVVARDLSTKMDHVISAFCVGNCRNA

SVVHLGSVEFSSEEAAEGQGPVGTEGTHLGDQPLSPVERCWLPDTYCEHLSSALRTNQNL

AELVLDLNALGNQGVKLLCQGLGHANCKLQNLGLKKCRFSSAACQDISSALSANQNLVMM

DLSNNALGDVGVKLLCAGLRHPKCRLQSLQLKKCYFGWAACEDLSSVLSTNPHLMELDLT

GNALGDAGVQLLCVGMKQPSCRLKTLWLKICHLTRASCVELASVLSMDCSLTELDLSLND

LEDAGVRLLCEGLGQPKSKLQKLRLGICRLTSAACGAISTALGPNGHLKELDLSFNDLGD

GGAHQLCGGLNHPNCKLQKLWLDSCCLTAIACESLSSVIAKHQTLTKLYLTNNALGDAGV

GLLCERLNQPTCKLRTLWLFGMELKAETQNTLAALRRTKPHLDIGS

>NLRP-like_4

MALSVRDLLTQTLEDLLETEFRKFKGKLCEIPLWDPPGGRGGAAGIPRGALEKADRLTTA

DLILSYCGSGSALDVVARVLENIQQRELGNRLRERAPDIGSSVKITGKLDTEKPRALPKR

TSGAAVCEGNCGSNNSDRGPCSNYNYGGVDKHLLNVKHRPKRRGRAKTIGPEAVPSHTGL

TIEGGKEDGDLIWKMGMKSGSPPWDNLITLEPPQRLERCFAHRKISQEMYRSEIRHKYER

VKDYNSLPGAWRSLEQHYVAPLIIRRCRPAREREQELLSKGPRHLELLRLCGEGGSARVR

LAHLLDGPGGRRPLTVVLQGAAGIGKSYTAHKIILAWASQRLYHDRFDWVFLFNCRELGV

EPRPRSLADLVLTDCPALGPHVGQIFSCPQRLLFLLDGFDELQPPRSPEEDEREEGEEGG

EEAAGRRAEAAVHRRRPAAATVRLLLRQRLLPGCLVVVTTRPSALEQLEGCIRADVHLEV

LGFLEPERQAFFGRFFGDAKRGREAYEAVRGNEALSTMCFVPLVCWIVCTVLHKQLEKGQ

GLDGLQTTTQVFLHFLSILLRFHRRPRARAPADSLLEQLGTLAVHGLLTRKVMFDREDLE

AHGLPTAAPPTIFLSAVLRQGVTVDTVYSFGHMMLQELFAAIFCFLPGRGGSARAGARPP

PGGGAAGGERPPAPGDPLLCQAMLRQLLPGRAALTPSPEGALLSWVQRSAASPRAGHPPF

LLELLHCLYEWHCDGLVGQVAGQLNVRFLLFPLKRSDCLALAYCLGCCPSVSCLHLYSCG

LDQGDIRLLLPALSKCQTLHLGLSDIPSGLMREIGCGFSPKQSVTSVLLQGLGPNNSSSQ

KETVFKVSALWGSRPCRLCTKRRLRCLQANRVGRAPRPVERARARESEAVSSNPRSANCQ

LGDFGQVILLILINRIY

>MIS12

MSVNPMTYETQFFGFTPQTCLLRIYVAFQDYLFEVMLTVERVILKRLEAAPGGGEGGGVS

PVQIRKGTEKFLRFLKERFDGLFATLETVLLQLVLRVPDHVLLPEDRSHARHPGGREELA

RLREEADRLRGRYEAEARAGRALMAELEEQRAARAQLEKTLRWFDGLEGAWREHGGGDPR

ESLAFLRRGAHRLRDVVGDVESKGRRLESA

>PGGHG

MGLTVFIPILQMRSPVPPPLPSLTLNDLATYFIRKINTVRPSRCKLTDAECRDLAEAVAS

SRTLMDLELMGNKLARTSTMDYGGVDDPAVFTSPTLPSDPRFLATLTNSYLGTRVYRDIL

HVNGVYNGALGDAHRADIPSPVNVRLEAPEGVDVSQSFTLDTRTGTFLHVVETREFTATH

RIYAHRALTHLLAFSVTVRRATPQGPPVTVRLRSDFTPKSQDLDLHLGPDFQGARLAAFA

CYFQVSNGSAGMFELRVDRPERSPEEGACRYLCGRTLSPEVAGGPQPTVHMLWTPAPPAL

TLPEAQREATWQFLAAVAEAEDEVRHLFEEGAALLRAGTLYPAHVEAWGALWGASGLHLD

GPLSLQRAVRGCLYYLLSAVPSPAPGVRDPFHGISPGGLSNGSRGEDYWGHVFWDQDLWM

FPNILLLWPEAARAILQYRVRTLSGAQANARDQGYKGAKFPWESAATGHEVCPQDIFGTQ

EIHVNGAVLLAFEQYYYSTRDLQLFKEEGGWDVVSAVAEFWCSRVTWSPEEQCYHLRGVI

PPDEYQTDVDNSVYTNVLAQNSLRFATSLGRDLGLAVPKEWLRVAENLKVPFDPKRRYHP

EYDGYHLGDLVKQADVVLLGFPVPCAMDPDVRRNNLEIYEVATTLQGPAMTWSMFAVGWL

ELKEPQRAQQLLNKCFDNISEPFKIWTENSDGTDAVNFLTGMGGFLQAVLFGYTGFRITK

DCLRFDPVCPAEVRHGQVTGVSYLGNKLNFSFSEEEVTVEVTWAQSQAPALEAMLETSGQ

CLALPQDCEPTVG

>PSMD13

MPRAAGAEGGRFPACPARRGTGSECAFPVGVAGEGLGNGRAPSGPPPQGIPRVPTRSLRP

VLGAPRGAPPWCPPPAAAAAAMKDVPGFLQQSQSSGPGQAAVWHRLEELYTKKLWHQLTL

QVLDFVQDPCFAKGDGLVKLYENFISEFEHRYDPPCVLCPGSRGRVRCRTDPSVALTFLE

KTREKVKSSDEAVILCKTAIGALKLNIGDLPVTKETIEDVEEMLNGLPGVTSVHSRFYDL

SSKYYQTVGNHASYYKDALRFLGCVDVKELPVSEQQERAFTLGLAGLLAEGVYNFGELLM

HPVLESLRGTDRQWLIDTLFAFNSGNVEKFQALKASWGQQAIIVAAQAGEPVHPELAAPR

SVQGTFLLYCPPPSTWNSAPHTPDLAANEALLLQKSQLLCLMEMTFTRPANHRQLTFEEI

AKSAKVTVNEVELLVMKALSVGLLKGSIDEVDRRVHMTWVQPRVLDLQQVRRRAPLCLGA

GGPLGIIIIIIIIIIIMMAFIKRLLCAKHCPKHWGGYKIKGMKERLESWCTDVKSMEMLV

EHQAHDILT

>MYADM

MPITVTRTTITTTSMSPGGGNHTIVGSPRALTTPLGIVRLLQLLFTCVAFSLVAHIGGWF
GSMGDWCMFSWCFCFAMTLVILLVEVGGLQPRVPVSWRNFPITFACYAALFCLSASIIYP
VAFVQYLHKGEQKDCGIAATVFSILAFLAYTTEVCWTRARPGEVTGYMATVPGLLKVVET
FVACVIFVFISNTNSYERHGALKWCLAVYCIFFILSLAAILLCVGECTSWLPCSFHTFLS
GYTLLAVLAYATATVLWPLYQFSSRYGGHSRPSYCPQNYGNPCLWDRLLAVAVLTAINLL
AYLADLIHSARLIFVHV

>EIF3F
MAAVPEATTTTTTTTAAAPDPAAAAAAPSPAVPAVPAAAPSAAPSVPVPGGPFPGGRVVR
LHPVILASIVDSYERRNDGAARVIGTLLDEGTEAQRSEVIIMTVFVKRLLCAKRCTKRRG
GYEVIRLSHVLALSLHPHFTDEVTEAQRNEGTEAQRSEVRIMTAFGKRLLRARHCTKRWG
GYEVIRLSHVLALSLHPHFTDEGTEAQRNEVTEAQRSEVIIMTAFVKRLLCAKHCTKRWG
GYEVIKLAHGGLTVFIPILQMRELRPREVRHCTKHRGGYEVIRLSHVLAHSLHPHFTDEG
TEAQRNEVTEAQRSEVIIMTAFVKRLLCAKHCTKRWGGYEVIKLAHGGLTVFIPILQMRE
LRPREHLQQCFGHRTIDKHSVEVTNCFSVPHNESEDEVAVDMEFAKNMYELHKKVSPSEL
ILGWYATGHDITEHSVLIHEYYSREAPNPIHLTVDTGLQNNRMSIKAYISASMGVPGKTM
GVMFTPLTVKYVYYDTERIGIDLITKTCFSPNRVIGLSTDLQQVGTASARIQDALSTVLQ
YAEDVLSGKVSADNTVGRFLTDLVNRVPQIPPEDFETMLNSNINDLLMVTYLANLTQSQI
ALNEKLLCL

>CASP1
MGRNLDLRSGQWEVAVLKAVPASPGALVVNLRIPQVPDQAKDQVCQLKERGPTCLLPLCQ
YCNTRIMNKELYNSLAIRSRNKFPLRASSYPNIPLILGHRKALSSPRLMEEQTPPLIFFP
GKGPLLGSCERKFQNSLTEQTGNAAKAVSLSPDLPAPAPQEAEGEWLRSSMAQWKEHRIW
SQRSRVQIPALPIVSCMTLAQLLKDRWCLIIESLTHGMISGLLDDLLQTQVINQEEMDTV
REENQRPAEKSRALLNSVIPKGDLASQIFIDALCKRNPFIAAKLGLSTGLVIPHSSRDMI
LLESLLTSERTEAQMGHFSSKPRIVAHHGLTPLFIRVASILVLPDHTHGAEEKFQLISRG
KQIQNSLSRFHSPKPTPEDRQGQSSNTQLVAHSPVYPAMAASCSAPQALQAPKALTESHP

DGPVEILRLCTSEEREKLQKENAGEIYPVLNKTGRKRQALIICNIKFDELLERVGAELDI

KGMKKLLEDLDYNVQIERNLSATEMESTLKLFAQQPEHKFSDSTFLVFMSHGILEGICGT

KFKTQDPDVLYYSTIFRIFNNLNCPGLRDKPKIIIVQACRGENEGMALVSDSLGASAYSS

QDLEDLENDAIHRTHVEKDLIAFCSSTPDNVSWRDPKTGSLFITQLIKCFQNHAWGCDLE

SLFMKCWTFSTAAFPQTGVDHTTYIPIDQSLQRAQHCAKHLGKNNTIELVDAILDWIQAN

GVNIHGQSSGNLDICGKQCELLWTGDISAKFALYSLKLLVQCSAPTIYLPHLDHQADNII

RAIIHQYHSKVEFGNRLEQCFTHRAEHGDTKQQQEEEEEEQQRSTCLSLAQKARCQPCHQ

NRHTEGKGRGMGRIRTLLIIHLPDRSIAIARINRFSSNLSQNSLQLSSVELPVILFYPLL

FTHPLPKNNNGTFCFTWQLDSPEAPSVRGRKNIIEFPIGDGGTEKLSDLTKVTLQSLGKW

ARTAAGNAYVEGLISSSRASRKNHTFLLPDATPCSEDQGTQ

>GRIA4

MGGWRGGRGGEEGTGSVWEGLLEEGWHRAFWAEKDTWTAAAPAAAPPPPAAASCLPLKAC

DLSARLYNMLNSLDSPYCKNTGFPKVLPPQFHQCCKQTKMVQECIRRSCQLERSRIYLIR

KMEIKTVSPMWDVQPEQLVSATVLRTGLAAYIYYTPSTVLNAGAKKQQGLMDRAWIQLCH

SSGYSILQAIMEKAGQNGWQVSAICVENFNDASYRRLLEDLDRRQEKKFVIDCEIERLQN

ILEQIVSVGKHVKGYHYIIANLGFKDISLERFMHGGANVTGFQLVDFSTPMVTKLMQRWK

KLDQREYPGSETPPKYTSALTYDGVLVMAETFRNLRRQKIDISRRGNAGDCLANPAAPWG

QGIDMERTLKQVRIQGLTGNVQFDHYGRRVNYTMDVFELKNTGPRKTQRVLGWCASRGCL

KAEQEIETPQKFGEDGLRGTQHRCQPQRSMNNKQNNLAEGCLSGLKCPLEEDTRQLENIG

HFWDATNPTDPIIPVHHYFLGQSGQDKILARTLLRRRLQILLTERVQNPSAALAHSQHSR

HDLGLTSITRQVQGTASQPLYFTDLPEAFDNINKPEVGYWNDMDKLVLIQDVPTLGNDTA

AIENRTVVVTTIMESPYVMFKKNHEMFEGNDKYEGYCVDLASEIAKHIGIKYKIAIVPDG

KYGARDAETKIWNGMVGELVYGKAEIAIAPLTITLVREEVIDFSKPFMSLGISIMIKKPQ

KSKPGVFSFLDPLAYEIWMCIVFAYIGVSVVLFLVSRFSPYEWHTEEPEDGKEGLSDQPP

NEFGIFNSLWFSLGAFMQQGCDISPRSLSGRIVGGVWWFFTLIIISSYTANLAAFLTVER

MVSPIESAEDLAKQTEIAYGTLDSGSTKEFFRRSKIAVYEKMWTYMKSAEPSVFTRTTAE

GVARVRKSKGKFAFLLESTMNEYIEQRKPCDTMKVGGNLDSKGYGVATPKGSPLRKWKHL

TTYIIPQFPKAEWPQNYKAYAAIPKGGGLRRTNQLQDQEQGSKPSVKYGSDTKWQYLLST

YCVRSVVLSIRGIQYELEDTIPALKDLTI

>PDGFD

MLLALGHVASHVRYFQVSLPKESSGGNGWNQLFQQILLHLYSKERNSHGSLNRNSRNPDR

CTLGPSIDTDTNMKVVIWQKFIDSTGTVGSRKQQYLYRKEETIHVTGNGCVQSPRFPNSY

PRNLLLTWRLSSQGNTRIQLAFDNQFGLEEPENDICRYDFVEVEDISETSTIIRGRWCGH

KEIPPRITSKTHRIKITFKSDDYFVAKPGFKIYYSPVGIAYHPPSVTDPTLTADALDQTV

AEFDTVEDLLKHFNPETWQEDLENLYLETPRYRGRSYHDRKSKVDLDRLNDDVKRYSCTP

RNYSVNLREELKLSNVVFFPRCLLVQRCGGNCGCGFPNWRSCTCSSGKTVKKFHEHAILF

PDIHSVKKNTISEQPTLPMLKFSGLKSTVSGSTLFGISGDEVTEAQSSVVTCPQYTSVRA

GIRIQVLLSPRPVLYPLGNAASLQEERLPKPRTGHPPRPALLLVRHSLLQTEGLVQSWQW

GSNCSNRSICSDVYGEGELEAA

>MTMR4

MAICRMGVRRMQWSGEGRAGTLGIVTVESEVAPAKGAKVKAKGNASHTGDSGSCFPHPAR

TLRSSAANLLTVPRSRLSRHRPPAHSPALVEGVTVRPSVRPSAGDMDGSPEPRRRRQIRR

FLEDPEEAELAQFVQEFPGGDGGGGGGCRRPESEEPSCRDPEALPAALEPSPEPDPRPP

ARPWPPDGHQHISAPAPLSPLTRPRSPWGKVDPYDSSEDDKEYVGFATLPNQVHRKSVKK

GFDFTLMVAGESGLGKSTLINSLFLTDLYRDRKLLNAEERITQTVEITKHSVEIEEKGIK

LRLTIVDTPGFGDAVNNTECWKPLADYIDQQFEQYFRDESGLNRKNIQDNRVHCCLYFIS

PFGHGLRPLDVEFLKALHQRVNIVPILAKADTLTPPEVEHKKRKVRGRGWRRRRKSGRKG

RERGRAQQPSPRQIREEIERFGIRIYQFPDCDSDEDEDFKLQDQALKDSIPFAVIGSNTV

VEARGRRVRGRLYPWGIVEVENPAHCDFVKLRTMLVRTHMQDLKDVTRETHYENYRAQCI

QSMTRMVVKERNRNKLTRESGTDFPIPTVPPGADAETEKLIREKDEELSWTLWPSLGNPA

SCRETRLSETHGLRAGPRVAATILRLLVPNPGPVGLLILGGSSRQTRRLEQCFAHSKRLI

NAITLILLSPHFTDEQRLEQCFAHSKRLINAITLILLSPHFTDEQRLEQCFAHSKRLINA

ITLILLSPHFTDEQCLEQCFAHSKRLINAITLILLSPHFTDEQRLDQCFAHSKRLINAII

LILLSPRFTDEQCLEQCFARSKRLTNAIIIIKTSRLEAGRRSKPPGPRAPTLVGWFLSWG

ASGTPPRSRSARRRTLVVVVVVVVVVVVVACPLGAPLGNRVFIGVILEPVGGEEGPPSLE

YIQAKDLFPPKELVKEEESLQVPFAVLQGEGVEFLGRAADALIAISNYRLHVKFKDSVIN

VPLRMIDSVESRDMFQLHISCKDSKVVRCHFSTFKQCQEWLSRLSRATAQPAKPEDLFAF

AYHAWCLGLTEEDQHTHLCQPGEPVRCRQETELARMGFDLHNVWRVSHINSNYKLCPSYP

QKLLVPVWITDKELENVASFRSWKRIPVVVYRHLRNGAAIARCSQPEISWWGWRNADDEY

LVTSIAKACALDPGGKAAGGAACSGNGQGSEAGDTDFDSSLTACSGVESSSGPQKLLILD

ARSYTAAVANRAKGGGCECEEYYPNCEVVFMGMANIHSIRNSFQYLRAVCSQMPDPSNWL

SALESTKWLQHLSVMLKAAVLVSNAVDGEGRPVLVHCSDGWDRTPQIVALAKILLDPYYR

TLEGFQVLVESDWLDFGHKFGDRCGHQENAEDQNEQCPVFLQWLDSVHQLLKQFPCLFEF

NEAFLVKLVQHTYSCLYGTFLANNPCERELRNIYKRTCSVWALLRAGNKNFHNFLYVPGS

ELVLHPVCHVRALHLWTAVYLPASSPCTLREESVDIYLAPAAQSQEFSGRSLDRLPKTRS

VDDLLSACDTSSPLTRTSSDPNLNNHCQEARVDLEPWSGQPEGADPPAGDHGPVGPPRPT

PELGHLPPPPASRKDFRTDKTLTRHKSCPPSCKVPSPVALWPPESGSLDPQLKGGEEIPE

PAPELPARDTSLRTRDGPGEPPEEPSGKEAAGPPADAASSGQDGTGNLPEAPCQELAPDA

SGKASQAPVPGQKAPDPGSDGRRAARRAGCPALEDPLAAPPLRRDPSGPAGSGASCQGPP

NPGPDPGPREEDGGRRENGRNGPSAESSRFGKAPPELGRKPVSQSQMSEFSFLGSNWESF

QGMVASLPSGEPAPRRLLSYGCCGKRSGGKPTRPTGLCPAGQWAQREGVRSPICSSQSGG

HCAGPAGKSNRTWPLGRPKPASGPKPAPASCPSPGPPLYLDDDGLPFPPDVVQQRLRQIE

AGYKQEVELLRRQVRELQMRLDIRHRCAPPAEPPVDYEDDFTCLKESDGSDAEDFGSDPS

EDGLSEASWEPVDKKETEVTRWVPDHMASHCYNCDCEFWLAKRRHHCRNCGNVFCAGCCH

LKLPIPDQQLYDPVLVCNACYEHIQVSRARELMNQHLKKPIATASS

>ABHD11

MLRGARAWRLRPPRSLCLARASTHGRPSTCVYLYVIIMAFIKRLLCAKRCCKRWGGYKVI

RPVPLSFTQFDGPTQETPLVFLHGLFGSKTNFQSIAKSLVQQTGRKVLTVDARNHGESTH

SSEMSYEAMSADLQALLSQLGLPRCVLIGHSMGGKTAMTLALQKASGPGPRVWPELVERL

VSVDISPEETTGVSDFLSFVEAMQAIQIPKELTRSQARKLADERLKPVIQEVSVRQFLIT

NLVEVAGRYVWRVNLEALTHHMDALMGFPQLPGTYSGPTLFLGGSNSPFIRPSHHPKIRR

LFPQAQILSVAGAGHWVHADQPRDFIAAIRGFLT

>IL1B

MPAYGEADSSREVEETDNQKTTRRPGLESPMARVPDQSSDLMDCYSGDGEEQFYEDNGPS

QVKSGFQDLKARTCQEARACQEDDCSPCQMGIELKVTELSSSRGFRRAVVLVVAMEKLKR
QVVACDMSFMDRDLMDIFTTIFKEEPVSCTSWEQTLVTDSIYHYLRCQEVTIWDQEHKSF
TLNTMANPCELRALHLIGANATQEVKLNMNFYYKTEHPAGPTVKQPVTLGIKGGNPGSLY
LSCVKKGDKPTLQLEVVNQSDLEGKNQERFIFNKSTEGTSTTFESAAYPNWYISTSREED
EPVFLGASKGEEAITNFYLN

>NEK7
MRINLKWKKSARCLDFCQQRSSAHAQERRRLTVEVIQGCGLVVRDRRRERGASVLSSGER
VVSRASLWSSREGSLGMEAKGGHDELRKLKGVKRSEVSVGKQHRFAERKCVGEKTGEKVV
KALRPDMGYNTLANFRIEKKIGRGQFSEVYRATCLLDGVPVALKKVQFNVQDRHSQLSMR
IESKAFEKVSLVHLAATLYVGRD

>GSDMD_like
MGVKTKTPKREKPDQVWEDAASTDKLVVATAGGEAGYRDWSSKEATAAATTVWLSGKSPG
FGVRGQKSSLCSRKLSQSLLSPLEGWQPSPDALLGRKGPISLFSPVFQLGVYHPAIPQAK
PARMFAQLTKKVAKKINAEGELLPLLSMNNSKRFRPLCLVRRKRKGTLFFGPRFRPTNLS
LLDVLDSDLPAPELKREDKFSLQDIVDGRFRAEVDLPDSLLSVKVSGEIKRVQKYSLEVQ
IVLISSEDLKRMQDERKLKKNEPEELKELRSLGENLYVVTEVVETLEEARLSSESKAEGS
CFLKLLSIHMKALCNHEEVVNIGKGCILAFRLGHLIFRDNWKILHTPTKEKTFPSEVLEK
GDPFVKDMAIAKDARGFEDLQREVNEEKQHLKYLDRQLKETILQAVQDLLGHREEMQKVE
DVLEDAMDGKGTQRLEGPGNIFLTTLEENSGQVMPELTGTVLYLLGALLVLSDTQQQLLK
LILEKGLLPQQLKLVKSILEQTFPMSQGGHFFLTLGPEDEEQSFTLALLEQYGLELPGPN
SSFLWKPDALASLSALYGALSLLDRRN

>ZC3H3
MRTARRPLWACAPPPARLLLMRRRRAPSPRQRGGPFAPEAGCRLMTSTAGGAQARAKAQA
QAQSRPPAMEEREQLRRQIRILQGLIDDYKNVHGNSRTPPAAGPRWPPPSYRGRGTFGVG
YSRPARRDFFPHQGPSWRKKYSLVNRPPGAPEQPEGGTPPSRDRPGPPPPDPRRQVQLRP
DQNMVIRIQAPSDAGSAGGSRARQDAPRSDPGLRKKDGEAGGSSREEGTSLVCRKEKGKP
RVVNSVGGGPGRPREPRWTASENARGLTRQALPARPQSSEGDAVGKAGSRIPDALCLQRL

RPGREPLLKNSLAPATKTTREPSLPGPCRTPKFRRTNYTWVASTVKAPRGPPRRPLSPRA
AAEGARKAPSSGAADRPVKPQPRADPGVKPRKPAAPSKPRGSSSKYRWKATGLTPAAAAP
FQWRAEAPGKSDGPPASPDGADFPAPGQASGGPGGWKPTFGESALSAYKVKSRTKIIKRR
GSVSLPVDKKSSLLPPATPKGHSSLRRKPSPRAKSSPASKRTPNRGAAQVTNHRLRRVPA
PRAHTPGKEAAALVGREPPAAAGTRSSPVPTCWRAVGPDAKGSTLDKPLDHQKEHCTKRL
TDLPVGPPDLRRSAVREIPQCSSESVCREPGILFYSGWIGGLRSGSVRETLWSGLRNREF
GGRRERRFGPGDGNLLETSEMPFH

>MROH6
MAARVGEGAQLELARAEGPGSPDPPPQPKAKPTRARRGRPKVTGHPRLVVAPGAQPPPCP
VGALTLAALAEEIQSHRGGQAGPSQQRAVQGDRQTAEPASRRPALEGNGGQHGAASAETG
EDTASRTRKKQQQQQPQQQPPQEEPRRESQPCSPAQSTLQPLASASPCTPGPEQFPLASC
FLTDLAVHTVACLTDAGFSGTQATAACLSSTLEAHGTILRDKVQELVHGLHLQIHRFSEG
RARRAALRVLCSLAVEHAQDVALRLPARTPGSFSSAVELWRGLSRNQRVNMMVLVQLLWK
LKGHPRLPGSSPAGSDGTLQEPLAATRALGEMLAVAGCVGAMRGFYPQMLIALVTQLHQL
ARCPPDSLSKACGHPQSKGGHPHGHAHCAVEALKALLRADGGRMVVTCMEQAGGWERLSG
PDTHLEGVLLLASAMVAHADHHLRGLFADLLPLLRSPDATRRLTAMAFFTGLLQSRPTVR
LLRAGAILERLGAWQGDPEPSVRWLGLLGLGHMALHAGKVRHVEILLPALLGALGEADGR
LVGAALGSLHRILLQPCGHSCTNSICLDVGARLWPLLDDGRDPVRCSAIGLFGTLVGRSP
VPQFCAIRDLVLDSLVPLLLHLQDQSPDAAEQSAEWTLARCDRFLHWGLLEEIVTMAHYD
SPEALSRTCQCLVQWYPGRVPGFLDQARGYLRSPQVPIRRAAGMFIGFLVHHTDAGTVRE
GLMDSLLHSLRELEWDPEASVRSTTHVTQHQLRLASQDWAARPGRFSPRRLLRPRGRPAR
PWPLYEEGPFKRRSRAGLWGSHMGA

>NAPRT
MGSRKPTCLPETVWEDKRKYDEAERRYYEREAAQAAPEASPPPEALNGLSQEESGDSGGQ
DPRMQKKRKRSPRHKTPGLDLALVGLSADHVWFDKPLFDRAERTFRAMLADGPAGEGPEA
EAAEPPPAGTPCDHGNRTACHHVVRGIWVNKFYFDRAERAFVERTQVSAPPRPLDLPALP
RPEAHQASGWGTPDEGYITAVPTPAAAGLPPAGEEGPTGSPPFALGWPGPSSPPANGKPQ
PAGLRAPTAEPRFASAERCFYQAAFDGHPPGKVRLQEREGWQDAARRGRKDRRNRNPPGK

RPKRAEPCGPKEADSPPPTCYFPREDSEPRWPGKPPLGGPGARHRAARTLRTARPALARK
APPAASAPGPLLDTRDPHSVSAIRPKKMATSFLMQEKIWFDKFKYDEAERKFYEQMNGPV
NSSSCPQENGASTILRDIARARENIQKSLAGLKTVLQSPPETPSQADAGASAPAGSSTGP
SGDHNELASRVASLEVENQSLRGGEKGPRETGPERGGWEGASSGAGVGSEAELLVGTEGS
PFCHTVIQDLQLAMSKLEARLSTLEKCSPSHRSPAPQTQHVSPMRKVEPSAPAPAPKAAT
PAEDDEDDEIDLFGSDDEEEDKEAARLREERLRQYAEKKSKKPGLIAKSSILLDVKPWDD
ETDMAKMEECVRSIQLDGLVWGGSKLVPVGYGIKKLQIQCVVEDDKVGTDILEEEITKFE
DYRGPDIGRGRSSRARGPGDARERNTTGSGPDQHRIKPGSGPDQDQTRTGSRQDQDRIRL
YQDRIRTGSGPDQDRIRPGSGPDQALPGQDQVRIRSGSDPDQDRIKTGSGADQDRIRPGP
GQNQARTRTGSRQDQTRTRTGSGPDQDRIRPGPGFTRTGSGPGPDQNRIRTGPGPEPDQD
QFRPRRSPDQDRTKPSPGPDQNQTKPRPGPDQDQRRPGRSPDQARVRTGRSPDQARIGRS
PDQDRTKPKPGPDQNQTKPRPGPDQDRTRPGRSPDQARIRTGRSPDQAQIRTGRGPGEAQ
TRPGSGPDEAQTRPGSGPDEAQTSPGSGPDEAQTRPGSGSDEAQTRPGSGPGEAQTRPRS
GPDEARAKPRPGPDQDRTRPRPGPDQDWTKPRPGPDQDRAKPRPGPDQDRTRPGRSPDQA
RIRTGRGPDQARIRTGRSPDQARIRTDEARAKPRPGPDQDRTRPGRSPDQARIRTGQSPD
QARIRTGRSPDQARTSPEPGLAQARARPIPVTMAYGYWRAGRAQEQAHFDLFFRRCPFGG
GFALAAGLRDCLRFLRRFRLRDPDIDYLASVLPPDTDPAFFDYLRGLDTSGVTVWALPEG
SVAFPMVPLLQVSGPLPVVQLLETTLLCLVNYASLVATNAARLRLIAGPEKRLLEMGLRR
AQGPDGGLSASIYSYLGGFDATSNVLAGQLCGIPVAGTLAHSFVTSFSGQEQLPTGALAP
GDLSAQAETWLTRVCEHLGRQVKDAHPGERAAFVAYALAFPRAFQGLLDSYSVMRSGLPN
FLAVALALADLGYRAIGVRLDSGDLIGQAQEIRRVFQNCAAHFQVPWLEFISIAVSNNVD
EALLAQLAQKGSEVNLIGIGTNVVTCPLQPSLGCVYKLVTVGGQPRLKLSEEEEKRTLPG
CKAAYRLGGPDGALLMDLLTLAEEPPPQAGQELRVWPLGSGQESRTLTPATVETLHRLYF
QRGQECESLPTLTEARALAQESLSRLSSAHKRREAPEPYQVALSEKLHALLESLCRSSRG
L

>SYK_like1
MDTYRAPSAVLDARRAQSSVLDAYRSAVLDTSQSSSVVFIERFLCAEHCTKRLEKYKLAT
YRDSPYPTTGSEHSAVLDTSYSVQSAVLGAYRAQGTVLSAYCRQRAVLGVYRVQRGVLDA
YCMQSAVLGVYCMQSAVLGLYPVQGTVLSAYHMQSAVLGVYCMQSAVLGPYPVQGTVLSA

YHMQSAVLGVYCMQSAVLDASHCIHSFIQSYLLGAYCSPSNPLMRLYKETRDALAHAKIV

PHAAGHPHQAAVNRAENAAELNPYVTQRGRRGPGDAQQEEPVWALPMDTAVYESPYADPE

ELRPDAVELDRGLLTLEDGELGAGNFGTVRKGFYRMKKGDKAVAVKILKDGGGGGGGVGS

GGVDEAVKEELLREADVMRRLDNPYIVRMIGLCRAEAWMLVMELADLGPLNKYLQKNRHV

QARNLTELVHQVCMGMRYLEEHSFVHRDLAARNVLLVTQHYAKISDFGLSKALGADQNYY

RSAVLGVYRVRSAVLGVYRVRSAVLDASYRVQSAVLGVYRVQSAVLDASYRGQSAVLGIY

CVQSAVLDASYHVQSAVLGVYRVQSAVLGTSYRVQSAVLGVYRVQSAVLGTSYRVQSAVL

GASCVQSAVLGVYRVQSSVLDASYRVQSAGLGVSWGQSAVLGVYCVRNTVPSIQCVQSSV

LGAHRRFVRPDATPLTAGGDAQTHGKWPVKWYAPECINYYTFSSKSDVWSFGVLMWEAFS

YGQKPYKGMKGSEVSAMLEKGERMQSPEGCPAEVYDLMNLCWTYKVEERPDFAAVELRLR

NYYYDISN

>Syk_like2

MCEPCRTGCVRAVCGPCGGHFVGRERAVSRRPCPSRVGALCEPCGDHVWALEGVMYEPWA

SRVTQAVGETCAGCVWAVREPCGGHFVGRERAVSHRLCASRVGALCEPCGDHVWAMGGVM

YEPCHAGHGGHFSGRERAVSRRPWASRVGALCEPCGGHVWATEGVMYEPCVSRVTQAMDE

TCAGRVRAVREPCGGHFVGRRVRAVWGPCANHVETICGPWKGSRVSHVTQAVDQTCAGRV

REPCGGHFVGRERAVSRRPCGGHFVGRGRAVWGPCTNHVEACVGQEWVMYEPCRTGCVRA

VCGPCGGHCVGRERAVSRRPCVSRVWALCEPCGDHVWALEGVMYEPCHTGHGRDMRRPCA

GRRVQAMCVPWFAGRGQSVGRPFCAPCESRAAQAVCMPRFAGHGRAMCVPRFAGRVRAMC

VPCFAGRVRAVSEPCAGHFVHHVQAVCRPWASRVQAVGESCAGHFAGRVKAMSCRPCAGY

VPATFCRPCAGYVRATFCRPCAGYVRAMICRPWAIRVHFARRVKAMLLRPCAARVQAMCM

PCFAGHVRARLVQCLASSKRFTNAIIIIIRLVQCLAPSKRFTNAIVIIIIIIIITIINLS

NGDEAQTPGAMASSGGDGWGHLPFFFGNITREEAEAHLEEAGLGEGLFLLRQSRSSLGGF

SLSVSSGGRVHHYTIERDVSGAFAISGGRSHPGPAELCAFHGREADGLVCRLGDPCVRPP

GLRPRAGPFEGLRETLIRDYVRTTWNLQVGGRGREAEIAGDGFRGSGGASIQGQALEQAI

LSQRPQLEKLIATTAHEKMDWFHGALTRSQAEDALLAAPRAEGKFLVRSREPAGSFALCL

LHGGRPLHYRIDKDKAGKLSIPDGKKFDTLWQWAHSLELSLRIKISIIEIVIDDNDDDDK

IIIIRININNDDDDDDDSPYPTWAHSLELSLRIKISIIEIVIDNNDDDDKMICIKINVNN

DDDDDSPYPAWAHSLELSLRIEISIIEIIIDNNDDDNKIIIKINVNNDDDDDDDDDDDSP

YPTWAHSLELSLRIKISIIEIVIDNNDDDNKIIIIKINIDDDDDDDDSPYPAWAHSLELS

LRIKISIIEIVIDDNDDDNKIIRIKIHVNNDDDDDDDSPYPTLVEHYSYKADGLLRALDS

ACPRKHNGSDVLGSRPPLPGDHPRAGGLIGRLHSFQRTKKSTVLSAWEVQVGNIERQSLP

NNGLGAQRCAGRLLLCAERCAGRRVQGTVLSAYRRQRAVLGVYRVQSGVLDAYCMQSTVL

SANCVQSAVLGPYPVQGTVLSAYRMQSAVLGVYRVQSAVLDAYCMQSAVLGPYPVQGTVL

SAYHMQSAVLGVYCMQSAVLDASHCIHSFIQPYLLGTYCVRSTVLSAWKSTNWQHIETVP

TQQWAQSTALYWTPPNHHQSYLLSAYCVQSTVLSAWEVQIGTLCRTLCWAPTVCRVLCWA

PINQSIVFIERFLCAEHCTKRQGKSKLAAYRDSPYPTVGSQSRSLSCAKRCAGRLLCAER

CAERLLFIKGAGAGRRSAGRQQGAPGVARMLRTAHARIPSSAPLPSPSNPLMRLYKETRD

ALAHAKIVPHAAGHPHQAAVNRAESAAELNPYVTQRGRRGPGDAQQEEPVWALPMDTAVY

ESPYADPEELRPDAVELDRGLLTLEDGELGAGNFGTVRKGFYRMKNAGAVVLICCRLGLP

ERSVQCSARRKRSINTIDDDDDDDDDDDDDDDDGEGRLEQCLAHSKRLTNIIIIIIITTII

NLSNGDEDLSAERGRPGGRVRSREPAGSFALCLLHGGRPLHYRIDKDKAGKLSIPDGKKF

DTLWQWAHSLELSLRIKISIIEIVIDDNDDDNEITCIKIKVNNDDDDDDDDDSPYPAWAH

SLELSLRIKISIIEIVIDDNDDDNEITCIKINVNNDDDDDDDRPYPAWAHSLELSLRIKI

SIIKIVIDNDDDDNKIICIKINVNNDDDDDDDDDDDSPYPAWAHSLELSLRIKISIMKIV

IDDDDDNEIICIKINVNNDDDDDDDDSPYPTWAHSLELSLRIKISIIEIVIDDNDDDDKI

ICIKIHVNNDDDDDDDDSPYPALVEHYSYKADGLLRALDSACPRKHNGQHPPPPPPPVTSP

LHPTRPVTSSGTEPAPPAHPDTWPLGAANPDCPRG

>AUH

MTSDSKARGLSIKRLLRAKHRAKRQGGVPGPPPDGRPPPRPPMEALKPVPQRDAALPPPP

SPSPSTSSRRRRREFTPEEKKDAQYWEKRRRNNEAAKRSREKRRLNDLVLESRLLALGRE

NAALRAELLALKARFGLLPPPPPSPSSAVKTEPPEPGAGPEADEAGGGKTPSDGEDEQRV

PKGPAALPHKLRLKFRGAPAAAPAEGPPPFALQSAVGPSVRARVGHGGDGVREALATRRR

RPVDRIHRSTSIHPAPASPGPGRSVDRIYQSINQSVDQYLPSAGFSRSQSVGRSDLINQS

VLIERWLLPVPIAVFLERLRCARRYTERLGESSTAADTFIELRPWVAEEKRLPGRARGRW

RRPRCRCRAGGWGGVWRGARFRFRWGPGAVRGWGSEPRDEDELSLRFLPDEDKGIAVLGL

NRPQAKNALSWNLIKQLSHSLDALKSDKKVRTVIVRSLVPGVFCAGADLKERAKMEAGQV

GAFVAKVRGVVHELVQCSAHGKRSINTIDDDDDLDFPKRLVQCSARGKRSINTIDDDDDD

DLDFPKRLVQCSAHGKRSINTIDDDDDDDLDFPRRLRLVQCSAHGKRSINTIDDDDDDLD

FPRRLVQCSARGKRSINTIDDDDDDDDLDFPSAYAWNGAGHIRLERCWAHSKRLINAIII

IIAMMIFSRVTLGESLPSSGPQFPHLLNGDEDLIPPYPPQRLERCRAHTRLPVPTIAALD

GLALGGGLELALACDIRVAAASAKMGLVETKLAVIPGAGGTQRLPRTVGPALAKELIFSG

RLLDGAEARAAGLVTHSPPQNGQGDAAYRRALALAREFLPQGPVAVRAAKLAINQGMEVD

LVTGLAIEEACYAQRQSSSSSIVFIERFLCAERCTERLGSPSRQHLETVIIIIIINRIY

>NFIL3

MTSDSKARGLSIKRLLRAKHRAKRQGGVPGPPPDGRPPPRPPMEALKPVPQRDAALPPPP

SPSPSTSSRRRRREFTPEEKKDAQYWEKRRRNNEAAKRSREKRRLNDLVLESRLLALGRE

NAALRAELLALKARFGLLPPPPPSPSSAVKTEPPEPGAGPEADEAGGGKTPSDGEDEQRV

PKGPAALPHKLRLKFRGAPAAAPAEGPPPFALQSAVGPSVRARVGHGGDGVREALATRRR

RPVDRIHRSTSIHPAPASPGPGRSVDRIYQSINQSVDQYLPSAGFSRSQSVGRSDLINQS

VLIERWLLPVPIAVFLERLRCARRYTERLGESSTAADTFIELRPWVAEEKRHREPAVG

>ROR2

MEEKGVEKEEKGVEEEEEQKEEGGRRKRRGEEEDERGEEEEEEEERRMKKGGGGGDGEKGG

EKEEEEEEEEQKEKGKREGEEDGFCQPYRGIACARFIGNQTIFVRSLQMQGDIENRVTAAL

TMIGTSTQLSDECSRFAIPSFCHFVFPLCEPGGGARPEGPTAPARPEGPEAPAAPARPRP

LCRDECEALESDLCRQEFGIARSNPLLLMRLELPRCRDLPPPGTPDAQRCVRLGVPPPPL

PTRGAHFTSPPGGARHTPPTPHHQGALTLPHHQGAPSTPNPHLTTRGRSLHLTA

>DIRAS2

MPEQSNDYRVVVFGAAGVGKSSLVLRFVRGTFRETYIPTIEDTYRQVISCDKNICTLQIT

DTTGSHQFPAMQRLSISKGHAFILVYSVTSKQSLEELQPIYEQICQIKGDVHKIPIMLVG

NKSDESQRELAAGEGEALAARWNCSFMETSAKMNYNVQELFQELLNLEKRRAVCLQVDGK

KAKQQKKKDKLKGKCSVM

>MUSK

MLKEEASADMQADFQREAALMAEFDDPNIVKLLGVCAVGKPMCLLFEYMAYGDLNEFLRN

MSPRTVCSLSHSNLAPRMRISSPGPPPLCCAEQLCIARQVAAGMAYLSERKFVHRDLATR

NCLVGENMVVKIADFGLSRNIYSADYYKANENDAIPIRWMPPESIFYNRYTTESDIWAYG

VVLWEIFSYGLQPYYGMAHEEVIYYVRDGNILSCPENCPLELYNLMRLCWSKLPADRPSF

TSIHRILARMCERAEAAMHA

>ZAP70_like

MDDSRIPKQLLYSELNWGGRKPGELTECFKDIVSSCGPEPPKDPKETPDRTAGEELRITS

WPMPCPGLVNPGPGGTAKVSPSPEATATQSAWPQCGPLAPAPLWEGASRGLGLIPGGCII

SASPGATAATVTGPAFRSPFPTPNGELGSATEMPDAAAHLPFFYGSISRAEAEEYLKLAG

MADGLFLLRQCLRSLGGYVLSLVHDLRFHHYPIERQLNGTYAIAGGKAHCGPAELCEFYS

RDADGLCCTLRKPCNRPSGLEPQAGVFDSLRETMVRDYVRQTWKLEGEALEQAIISQAPQ

VEKLIATTAHERMPWYHNAISRDEAERMLFSGSQPDGKFLLRPRKEQGTYALSLIYGKTV

YHYLITQDKAGKYCIPEGTKFDTLWQLVEYLKLKADGLIYCLKESCPNASVPTGTAAPTL

PAHPSMPRRIDTLNSDGYTPEPESCQVGLGRAAGGCSRKKAEAEPCGSGRLVAGGDAKAG

EKSRILPMDTSVYESPYSDPEELKDKKLFLKRENLMMDEVELGSGNFGCVRKGVYKMRKK

QIDVAIKVLKSGNEKAEKEEMMKEAQIMHQLDNPYIVRIIGVCRAEALMLVMEMALGGPL

HKFLSSKKEEIPVSNVVELMHQVSMGMKYLEEKNFVHRDLAARNVLLVNQHYAKISDFGL

SKALGADDSYYTARSAGKWPLKWYAPECINFRKFSSRSDVWSYGITMWEAFTYGQKPYKK

MKGPEVISFVEQGKRLERPTDCPPEMYTLMNDCWIYKWEDRPDFSMVETRIRTYYYSIAS

KADLATSPVQGAEAACA

>ADAMTS10

MAIACRLLSWALAFSLSLPSQPASAFQSQEEFLSSLKSYEITFPVRVDHNGAFLDFAPPQ

RQRRSLGTRPPEPAEPRVFYKVEALHTRFLLNLTLTSHLLADHFSVEYWKRDGLDWRHHI

RQECLYAGHLQGQRLSSKVAISNCHGLHGLIVADEEEYFIEPLNGRGSGVPEGEGSPHVV

YKRSSLQRPHLDAACGVLAPAMVGHPSQSFGLLADEKPWKGRPWWLRPLKTAPTKPLGNQ

TQRGQLALKRSVSQERYVETLVVADRMMVAYHGRRDVEQYVLAIMNIVAKLFQDSSLGNI

VNILVTRLILLTEDQPTLEINHHAGKSLDSFCKWQKSIVNRNGHGNAIPENGIANHDTAV

LITRYDICIYKNKPCGTLGLAPVGGMCERERSCSINEDIGLATAFTIAHEIGHTFGMNHD

GVGNGCGARGHETAKLMAAHITMKTNPFVWSSCSRDYITSFLEHRLFPRNSRALTHASEK

QRGSVERARALESEAVGSNPGSATCQLCDFGSGLGLCLNNAPPKQDFVYPTMAPGQAYDA

DEQCRFQYGVKSRQCKYGEVCSELWCLSKSNRCITNSIPAAEGTICQSSTIDKGWCYKRV

CVPFGSRPEGVDGAWGLWAPWAECSRTCGGGVSSSTRHCDSPRPTIGGKYCLGERKRYRS

CNTDDCPPGSQDFRELQCSEFDSVPFRGKYYTWRTYRGGGVKSCSLNCLAEGFNFYTERA

AAVVDGTPCRPDTIDICVNGECKHVGCDRILGSDLREDKCRVCGGDGSSCETIEGVFAPT

LTEGGYEEVIWIPKGSVHISIRNLNLSLSHLALKGENDAFLLEGKPGPSPQLRLPLAGTI

FHLRRGPDQPECLEALGPTNATLIVMVLVRSELQGIRYRFNAPITHEALPPTYTWHYAPW

TKCSALCAGGSQVQAAECRKQPDGSPVPLHHCKAHAKLPERQRSCNTEPCPPSWAVGNWS

GCSRSCNMGARTRSVVCQRRMSPNEEKTLDDSACAQPRPHVLEPCSSQSCPPEWAALDWS

ECNPSCGPGLRHRVVLCKSGDHSATLPTSQCSAATKPPTSMRCNLRRCPPPRWVAGEWGE

CSAQCGFGQQLRSVQCTTHTGQPSGDCTAALQPPATQQCESKCESSPTESPEECRDVNKV

AYCPLVLKFKFCSRSYFRQMCCKTCLGR

>MYO1F

MGSKERFHWQSHNVKQSGVDDMVLLPRVSEEAIVENLKKRFLDDYIFASSNEGWGREGWS

EFPQTDSHWLNVKATQEKVNQLGGGVHTGGERDSPEIVALDPAAQTYIGSVLISVNPFKQ

MPYFTDREIELYQGAAQYENPPHIYALTDNMYRNMLIDGENQCVIISGESGAGKTVAAKY

IMGYISKVSGGGDKVQHVKDIILQSNPLLEAFGNAKTVRNNNSSRFGKYFEIQFSRGGEP

DGGKISNFLLEKSRVVTQNESERGFHIYYQLIEGASQDQRQNLGIMTPDYYYYLNQSETY

KVDDTDDRSDFHETLNAMQVIGIPTEVQQLVLQIVAGILHLGNISFCEQGNYAQVESADL

LAFPAYLLGVDSGRLNEKLTSRKMDSKWGGRSESIDVTLNVEQAAYSRDALAKGLYARLF

DFLVEAINRAMQKPHQEYSIGVLDIYGFEIFQRNGFEQFCINFVNEKLQQIFIELTLKAE

QEEYVQEGIKWTPIEYFNNKVVCDLIENKLNPPGIMSVLDDVCATMHATGGGADQTLLQK

LQAAVGCHEHFNSWSSGFVIHHYAGKVSYDINGFCERNRDVLFSDIIELMQSSEHAFIRM

LFPEKLDADKKGRPTTAGSKIKRQANELVSTLMKCTPHYIRCIKPNETKRPRDWEESRVK

HQVEYLGLKENIRVRRAGFAYRRPFQKFLQRYAILTPETWPHWRGDERQGVQHLLHSVHM

EPDQYQMGRTKVFVKNPESLFLLEEMRERKFDGFARTIQKAWRRHVAVRKYEQMREEASN

ILLNKKERRRNSLNRNFVGDYLGLEERPELRRFLGKRERVDFADSVTKYDRRFKSIKRDL

ILTPKRLYVIGREKVKKGPEKGQVQEVLKKHLDIQVLRSVSLSTRQDDFFILHEEAADIL

LESIFKTELLSLLCKRFEEVTQRALPLTFNDTLQFRVKKEGWGGGGSRNVNFSRGSGEMA

TLKISGKTLMVSIGDGLPKSSKPTKKGAPRSQSRGRRPAPARSAPGPPRGTCRNGAPPAM

APGSSQQRLEQMYAGHQKQSRGPPAAMLPKQGASRRMRARPPSEQNLEFLNVPDQGMAGM

QRKRSIGPRPPPGVGRPKPQPRAPGPRCRALYQYVGQDVDELSFNVNEVIDILLEDEVIE

AQRSENGAVEDGADVSAAPRRRGSRIRTKAPGGGEGRSGMDQQEEKPSPGSGGQAGPAPV

PFPDLYTSSSRNQGPKLEKDQLAPTFPLGDCSSPCLSGSGRTNVVPSTSSSRHTQDLRLQ

KRRPLPGKQYPCSSYGCKLVCSSSQELAHHLRSHYLPTQSMGGKLFHCSTLGCADTFPSM

QELVTHMKVHYKPNRYFKCENCLLRFRTHRSLFKHLHVCSDPSRSPTAGPPPPALEKEPP

EPEPSAGPSPEAAPLLSPLPLGPSHTQPFPLLEPSLFDPASLPRFPAQASSPMPGAFLPY

LPPSPYSLPPGSGQQRLRPFLPAQALPISNAIWKKSQGVSGSPRRPPGGSEVLAVFLKLQ

SCRRRHYCCFCSWETGAQQGLACQVLCGRRKQGHSSNSRIVWEHTRGRYTCMQCPFSTAS

RPAMTLHLEDHRKTPPPPARLDAHMDFGVGLAAFPSKLPAEMESSLYSQL

>TMEM131

MAAFIQSENIMEVLRFDDGGLLQTDAPIGLGSYQQKSVSLYRGNCRPIRFEPPMLDFHEQ

PVGMPKMEKVYLHNPSSEETITLISISATTSHFHASFFQNRKIPPGGNTSFDVVFLARVV

GNVENTLFINTSNHGIFTYQVVEMFSSGGDLHLELPTGQQSGTSKLWVSERQSSSPGEGG

DQKPGVESGQRKMRGSGMPAPWGRRAPLPVWGKTDNFCPGVFWIAVIIRCLSRYSVGAQY

SVRPQEIPPYETKGVMRASFSSREADNHTAFIRIKTNASDSTEFIILPVEVEVTTEMLDF

GTLRSQGKVLYIDTLGKFLGPLLFSIYTHSLGEFIRSHVFNYHLYADDTQIYISAPALSP

SLQARISSCLQDISIWMSARHLKLNMSKTELLIFPPKPCPLPDFPITVDGTTILPISQAC

NLGVILDSALSFTPHIQSVTKTCRSHLRNIAKILPFLSVQTTTLLVQSLILSRLDYCIRL

LSDLPSSCLSPLQSTLHAAARIVFVQKRSGHVTPLLKNLQWLPFNLRIRQKLLTLSFKAL

HHLASSYLSFSSPARTLCSSATNLLTVPRSCLSRRRPPAHILPLAWSALPLHIRQASSLP

AFKALLRAHLLQEAFPD

>CLEC_like1

MDNEVTYADLKFQDSFKPQRIQEFDNSREIEHPAPSPAWRWSALGLLTLCLLLLIGLASL

GILYHACKPCPETWFWHEKSCYGASIVKQTWEDSREACAAVNSSLVKIDNKEEWDFIASL

QNRQYHWVGLYQNPNGRQWEWEDGSALSRDLNSLVSGDRTGGKMCAYTYGSYFYIDPCTD

KHYYICEKVAGLVKKLIAG

>CLEC_like2

MPREHLESAIEVALTPFSQIQALDAQRQIELLFQAEFLFPAFLLWPTANPWATQTSLKAA

DFEEVPGAFLSFGLEEPEATTGSWSSMGVSHNQWGLLGYPGYRKRANRAGGLQSLLMFQL

TSPDSPGSGRSPCGQGSGGLKGGGGEEEMANEVTYANLKFQDSSMAQRIQKFDTIQEIAE

AMSFSERDINRPLGFIEKQRSSVERARALESECLEKCFARSKRLINAIIIIIKGRTLIET

ITGIPTKKPWATDFLTAACEFHCGLPGTARGAAVIRPFLWTRVFQEKKNIQQLNGVKENL

SLQLDISANISKEKDLIQSDLSDALKKMATKLCRELTRNKQAPDCLSNLVSEHACKPCPE

KWQWHRDSCYWIARKLNLEKSRKVCAENNSSLVKIDNKEELVFVASKLNPYYWVGLSRNI

SSGQWVWEDGSTLSPDLQTGRTCFRAGLQALPGEVVFTQGQLCEDFEGKQTRDESRDSCA

TQNSSLLNIDNKKEWMEQRAALAIHATTIGERHPTLGYRSLLQGYQVFWGEAPASSDLPV

HFHVWRLLSLLLLTLCVLQLIGVVGFGIRYSILFRGICRDCFKQKEFLLSQNSNLSADLR

EVTTKLCQELIMKQPDHACRPCPEGWHWHGYNCFKILMDKRTWYESREACAFQNSSLMKI

DNKEEWNLLTPKIQSYHWVGLSRNASDLSLKWEDGSEINPKVLLLLSDAKTKGRLCAVVY

RNDLSLDFCNNTYPFICEKAAEPVKKELLT

>CLEC_like3

MKEEVTYADLSFQRTDDVEKVPQMDLIEKPVSSLPQLGSSSKGSRGNWPIFRCFGVWAGG

LKPGGLDLGNTAYSEFCLSVLDRTICSPVCEEICSPGSADTVFAATAAAVAAAERTRGPR

DPLCFNGFLFPESKCSPCEKGWQWSGDSCYRKFDIWNTWPKGKKFCHDNNSTLVKVDSRE

ELLIGLVVLGIKFSQARSQEIGGTSSPNDKPENLDINGSCSKQQEFLLSQNSKLSAGLRK

MAIELCREVTRNKPGLRREEESANLVDVQKKGVPGQGEDVGWWSTAGQVRTKHSEEFSSR

GAEGAGWAVEGKKGESKCSPCEKGWQWSGDSCYRKFDIWNTWPKGKKFCHDNNSTLVKVD

SREELLIGLVVLGIKFSQARSQEIGGTSSPNDKPENLDINGSCSKQQEFLLSQNSKLSAG

LRKMAIELCREVTRNKPESKCSPCGKGWQWSGDSCYRKFDIWTTWPKGKKICHDNNSTLV

KVDSREELVRSISMELTGLKAENVEDEGGYTMMSIYRQTSIRGLAGSEPARSKASMEDED

GYTMLNLKSSRIHDFSKGLSGHKCNPCSSSKYHQGNCYHLYLRNRTWEENRIYCASKNYT

LVKVDNQEELAYLTRSTHKIRWIGLSRTANDAPWTWEDGSVPAVDLFQLSGDEEAKHCAL

FHNGKIEAAGCQENYPSLCESVGGNIKIDLLL

>CLEC_like4

MQDEDGYIMLDFKSRIHATSKGPSESGCPGLDRHGNLPALKGRGLSADPPGLFPSWCWMT

LALLILCLGMLIGLIALGSMWLSTVLCTHLTKGPRKGGVVEGKESSNKRSDQLYDVRGMV

DIHSPASTNTDSRQQALLQQITQKYCQELSSKPGGHKCSSCDNNWRFHGGKCYGTFKNNK

TWEESKKYCDDRNSTLLKIDTQEAWNFIQGKPDFTRWIGLSRPSSGGRWMWMDNSALTDN

LFELSGDGDEGKHCAYIQKKQISTTFCRELHYYICEKLPPRPSADSAILPEPPAPLIILE

LRLSGQSHYLEVMQDQGAYDSLCWVTPDPPPVKRPSLRTEHQDTQQQEQPLQQQQQQLM

CLSLPRRPDACSTTRTNTLRQGPRDRNLVAVSYGDVLALLPAFAAFHCHPEPQVLLRCHV

FMSKIELFVFPPKPCPLPDFPISVDGTTILPVSQARNLGVILDSARSFTPHIQAITKTCW

SQLRNIAKIHPFLSIQTATLLVQALILSHLDYCISLLSDLPSSCLSPLQSILHAAAQIVF

VQKRSGHVTPLLKNLQWLPINLRIRQKLLTLGFKAVHHLAPSYLTSLLSFSSPARTLHSS

AANLLTVPRSRLSRRQPPAHVIPLAWNALPPHICQASSLSPFKALLRAHLLQEAFPD

>CLEC_like5

MAYVQSDQLVSTSALSTVPGNVIQASELIGKQEEILANLSHQQHVCSESLQMCQVRIRMF

TSPESNCSPCLEPWVKNGKSCYLFFDQWKNWTSSSEFCLQEKSELLKIGSKEELNFINQN

IEKKKMGSSWSYWVGLMQDGCFGDWRWRDSTVPSSDLWPKQGSWSAGETCGHLTHGVLSS

ASCSKWKYLICEKCTSSAVDFQLD

>CLEC_like6

MRRWRKRHCYCFTSYETETVPDLVILYLGQCFAYRHVAPTRTWRPVALTLLILCFVLLFG

LGALGFEFFQVFRLSNTQRTAISQQEERLGNLSQQLQDLQAQNRKLTGTLQVAQKLCWEL

YNKTGARNLGVILDSALSFTPHIQAVTETSRSQLRNIAKIRPFLSIQTATLLVQALILSR

LDYCISLLSDLPSSCLSPLQSILHATARIVFVQKRSGHVTPLLKNLQWLPINLRIRQKLL

TLGFKAVHHLAPSYLTSLLSSSSPASTLPPPPPSSATNFLTVPHSRLSRRRPPAHVVPLA

WNALPLHIRQASSLPPFKALLRAHLLQEAFPD

>CLEC_like7

MQDEDGYTILPPYVRLNPCHPAASDKGQSCLDSRMEDEDGYTILNPRTRAFARDPAASDK

GLPAVSPRWRPAAVTLGIVCLGLLGATGILASPQTTTPRDLEQQQREPASWQLGEETTGV

GVKDFVFLRIPNSCRSITKQSYDSVDTSLPERPASICNHSGRVFSVKIQKWSFVAFFHTV

NLSLCLRLTPTLLLLPRTASHLPPFVPDDHAKQLQGISSVAQWKEPGLWSQRGLNAGESH

AVGNGNSQRLAEGLSPPQRRRPRECRPSPPLRAGGGQRWRRSSLGNQSVVQMRAKHSSTQ

DMLEDDGDTTLSLHSRTSTAAGSPKPAGPDCEPAVG

>CLEC_like8

MGIKSVSPMWDNLITLYPPQRLEQCFAHMCISAGVSVWACAYLQDMEDSVQYAELKFKMP

EEKPKQKLPEEKAKDSSSLSPWWFPAAIVLGIFSFGLLGAVAVLGLEINQAHGLMNQQVK

NYTHLEGETAQFTDQKEVAKATPRDQEHVEDLLGKLENITEERNALLLQNEHLQEALKKV

KNYTGPCPQDWLWHTESCYSFPSRSTNWKESQENCASQGAQLLKVDNQDELEFIYQATVH

SRNPFWLGLRRSEAISRWLWADGSAPFVGLLQAWRYISHTYPSGTCAYIFQDNIFAENCI

ITAFSICERKANLLKLQ

>CLEC_like9

MASEIVYAEVKFKMDTQKLGAPPAPAPQKATPPPLRPWLPGLLMALLLLLLLLLLLSFLV

AFVVFYRRSHLCREDRSSKSTRILSEFVCAKQSSKTPVFSCSYIETHQENEDPPSPHQFP

KNQTTLDCITEGSEVEGRSWDCCSIGWRPFRSRCYFISTDKMPWAESQQNCSQMGAHLVV

ISSDTEQKFLESILDNHDVYFLGLTDLEGSKDWRWVDQTPYNKSVLFWHPGEPNYSWERC

ASLHWINYRGWGWNNIRCNEKQNRISRQWSKSPAASRPVECPPLSCSIVIPSDLLGIEVT

NGQRGVLQPAAE

>CLEC_like10

MVQTVCKHSVNVIDWLYERNSQAQTDRGGQCPRGAHIGLTDIFERRDSQTMVQSTFASPT

LPASTIIVTGISILLLTSCFTASCLVTQNGFAQLCKDSNGLAPLNHTQLSCLRERPQVEG

RMWSCCPQDWIPFQTSCYLFPTDVLPWDEAQSKCLKQQAHLVVINTEAEKNFITQGKPFN

FSIHLGLKKWKMESQWRWEDKTPYNPADTFGLEGDLAEGDGARRCAGLTLGQPSIRWRWS
STDCAAAAHRICEMPRRSF

>CLEC_like11
MVSILLLSACFIARCLVTQRAFSRFCKEEGTLLRLNDIFTEISCYSSGSGSIQDCCPRGW
KHFQSHCYFFSSDTLTWSSSLLNCTGMGAQLVVINSLEEQEFIFHSKPSGREFYLGLTDQ
QVEGKWTWVDGTAYDPSWSFWDLGEPNNILGLEDCATTRDSPSAKETWNDVTCFSFHYRI
CEMPAQTLSAGKKGM

>NECAP
MAAEAEYESILCVKPDISVYRIPPRASNRAYRNFPSPSPPPGERYPPPSGDRAPILIIHC
WVGTVSICCQLVLPKRLVQCSAHKHCTTRLGEYNITDAFPDHSQLSGKNYGPGNPLLTHT
ALMASDWKLDQPDWTGRLRITSKGKVAYIKLEDKVSGELFAQAPVDQYPGIAVETVTDSS
RYFVIRIQDGTGRSAFIGIGFSDRGDAFDFNVSLQDHFKWVKQESEISREAEKPDTHPKL
DLGFKEGETIKLSIGNITTKKGGPTKPRPVGAGGLSLLPPPPGGKISIPPPASSVAICNH
VTPPPVQTSSQGGGESDILLDLDAPAPSTKASASVPAAPDLWGGFSTAARMLTNKFLFSA
HSAASDTGSVPGGPCKSPHPQKHYSQKLPVPLIPLCSVFKSMSPR

>C3AR1
MGTDCEPHVGQPDHVVSPQRLEQCFAHNKRLTNAIIIIIIMGTCRGHRMGRMSVPWGNTS
SPDSPLQQSHSLPELLSMVILSFTLVLGLLGNGLVMWVAGWKMQRTVTTVWFLHLTAADL
LCCLSLPFSLAHLALGGHWPHGQLLCKLVPATIVLNMFASVFLLTVISLDRCLLVTRPVW
CQNHRGVRLATAVCGAAWVLAFVLCLPILLYRETYVDGPLWRCGYNFGHHASLDGLEDTG
LMDIGLVGFSDSSPPTPEMDDLGTLDVPLRARDALLTGSDWLLGSSLPWDPLDTKAGHPS
SIPNPDLSPRGDLLPTEMPSQPPGELDLDTDFWDDLIKIYAAESQVPGPLVAMTLTRLGL
GFLVPLAIMVACYSVTVVRVQSSRFAVGQTRTFWLATRVVAAFFTCWAPYHLVGVLSLLA
TPDSPLDHAVAALDPLTQALASANSCINPLLYALAARDFRVRARLSLRSILEAAFSEGLS
HSSTCPPSQAVALSIDPLGTEV

>BIRC2

MPRGSFIELTRAGGTLLQEPRPLSSGQDQPPHRGRLVPYPPSPTKYLIMNIVANSTFLSD

LMNGNNGYELKYDFSCELFRMSTYSTFPTNVPVSERSLARAGFYYTGASDKVKCFSCGLM

LDNWKPGDSAIEKHKQLYPSCSFIQNLLQTHNPGASSYSAFCSPPLGSLSPTATISPSLE

PSGYFSGSFSSFPLAPVTSRTIEDLSQLRPHVDNSAMSSEEARFRTFQSWPLSFLSPSAL

AKAGFYYTGPGDRVTCFTCGGKLSNWEPKDDAMSEHRRHFPGCPFLERQTRDASRFNVSN

ASMQTHAARVKTFLNWPARIPVQPEQLASAGFYYVVLPSRSGAADCTSSLTNFLSGAILD

FLGRNDDVKCFCCDGGLRCWESGDDPWIEHAKWFPRCEYMIRMKGQEFVNQIQARYPHLL

EQLLSTSDTPVDESANPPIIHFGPGENPSEDAIMMSNPVVKAALEMGFSRRLVKQTVQSK

ILTTGENYKTVNDLVSDLLNAEDETREEEKERQNEEMASDDLSLIRKNRMALFQHLTSVL

PILDSLLSASVISEQERDVIKQKTQTSLQARELIDTILVKGNAAANIFKNCLKENDCVLY

KNLFVEKNMKYIPTEDVSGLAFWSQVVTDAVEGALKGVAVLELQENFLKRNQGSHGEVFL

HMHPAAHCCCVPFPRCTSNQDFSLAYDFCVKP

>CARD9

MATNCTVLFQVLNTVLCTQRSGAGHGQTQTASPSQALSGGGPGRMMGSERGGGFFLESCH

PNCPLLPPRAFWRHLTDWRSAGDDVEAKKTEVKGRLGVSRSGSEVIAAKKRNGGAPGGRG

PERDRRSEEPSEMFGPRFLPYRGVGEPVSWLRLRVVVIVFVTTPTRGDPNGSPGGRDGAL

APCCKLCIWLPSRKTYYQGTTLQASISSPVASGRMIPGILKPRHGDKMQRHLSVAGIGSA

PTTGIFQARIVLREASSGCHLKRSPSREVHWAPAMSDYENDEECWNTLEGFRVKLISTID

PSRITPYLRQCKVINPDDEEQVLNDPSLVIRKRKVGVLLDILQRTGHKGYVAFLESLELY

YPHLYKKITGKEPTRVFSMIIDASGESGLTQLLMNEVLKLQKKVQELNLLLNSKEDFIKE

TRVKNSMLRKHQERAEKMKEECQAFSQELKKCKDENYNLAMSFAKQSEEKNAALMKNRDL

QLEIAQLKHNLMKAEDDCKVERKHTTKLKHAIEQRPSQEVMWEMQQEKDLLLAKIQELEN

SIREGKREKNSLYITVLEEDWRQSLMEHQEKETTIFHLRKDLRQAEALRNKCMEEKEMFE

LQCTTLRKDSKMYKDRIEAILQQMEEVAIERDQAIMTREQFHLQYSQSLIEKDGHRKQIR

ELGEKCDELQLQLFTREGQLLSMESKLKRLQLETPNLSSDLEETSPRNSQELTLPRSLDE

DAQLSDKSEPAILPSLAIIEKSPGEGSDL

>ARAF

MGAGSRAEGAGPARAAAPAGSSPQNRSQDGGDGGGGVVTEHCTKRLGSTSRQHIETVPTQ

QRAYSLEDYQASDKGPVCLGTSPGTHAGARLIIHGMQPWARAASLNPPCLPPGAPLMKRS

LGAAGLISNGNHLASPEPPMSPAPGPMEPARRPAPDGGEAGRTGGTVKVYLPNKQRTVVR

GPRGWGGEKVTVRAGMSVYDSLDKALKVRGLNQDCCVVYRLIRGRKTVTDWGTAIAPLDG

EELIVEVLEDVPLTMHNFVRKTFFNLAYCDFCHKFLFNGFRCQTCGYKFHQHCSSKVPTV

CVDMATARHSIHAYASVQDLSGTGVQEPLSSQSLREPLTPQPISSQQHLSPEAFPFLEAP

AQALQRIRSTSTPNVHMVSTTGPVDSSLIKRAARNFNAEGTGNGEEGQPQGTGGGSSGPP

LVSPGRKYSHPKSPSEPRERKASADDKKKVKALGYRDSSYYWEVPKSEVQLLKRIGAGSF

GTVYKGKWHGDVAVKILKVTSPTSEQVQAFKNEMQVLRKTRHVNILLFMGFMTRPKFAII

TQWCEGSSLYHHLHVAETRFDMVQLIDVARQTAQGMDYLHAKNIIHRDLKSNSIHSMGPQ

EMRDIFLHEGLTVKIGDFGLATVKTRWSGAQQVEQPSGSVLWMAPEVIRMQDSNPYSFQS

DVYAYGVVLYELMTGSLPYSHINNRDQIIFMVGRGYLSPDLSKISSNCPKAMRRLLPDCL

KFKREERPLFPQILAAIEQLQRALPKIERSASEPSLHRAQADELPPFLFSTLRLVP

>RAF1

MGPLGVVPSSGKWEVGPRSPLCPRGETGDRPRQAPEVIRMQDNNPFSFQSDVYSYGIVLY

ELMTGELPYSHINNRDQIIFMVGRGYASPDLSKLYKNCPKAMKRLVADCVKKVKEERPLF

PQILSSIELLQHSLPKINRSASEPSLHRATHAEDINSCTLTAATRLPVF
