## Supplementary material for "Large-Scale Restructuring of the Caspase-1 Gene Cluster Region in Mammals": Online Resources 4, 6-10

Journal of Molecular Evolution

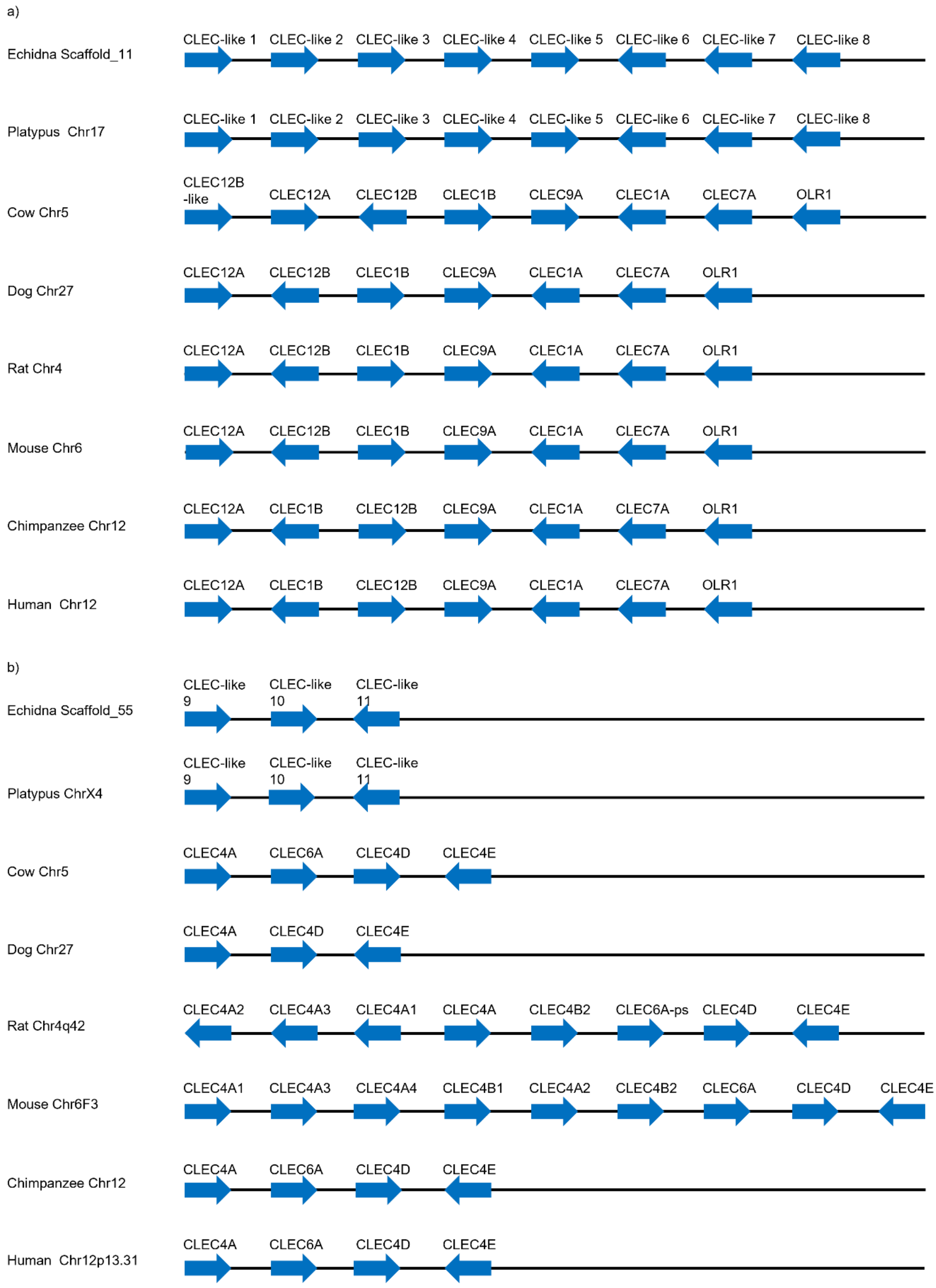


Online Resource 4 a) Chromosomal arrangements of the Dectin-1 subfamily in multiple mammalian species. The arrows represent the orientation of the genes. b) Chromosomal arrangements of the Dectin-2 subfamily in multiple mammalian species. The arrows represent the orientation of the genes. Distances not to scale

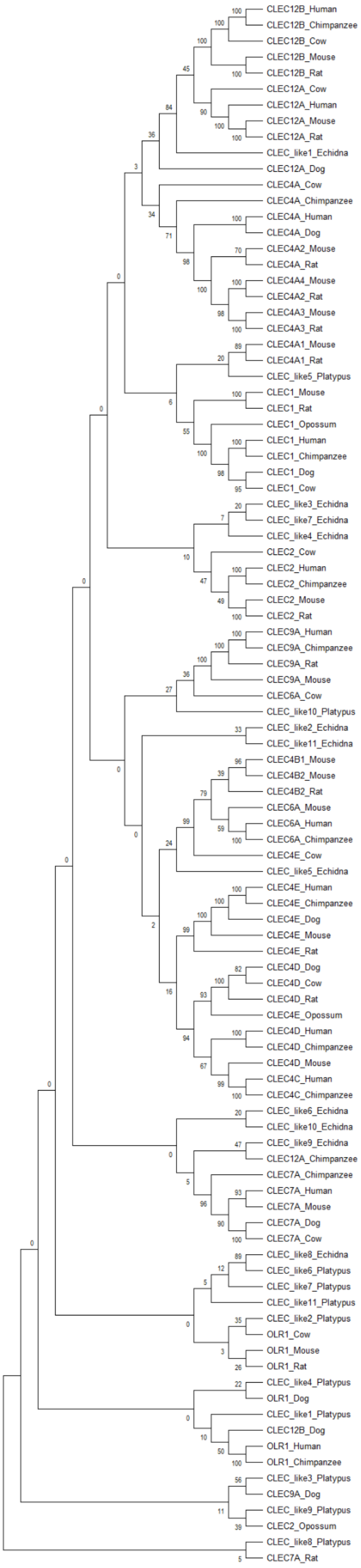
CLEC12B_Human
CLEC12B_Chimpanzee
CLEC12B_Cow
CLEC12B_Mouse
CLEC12B_Rat
CLEC12A_Cow
CLEC12A_Human
CLEC12A_Mouse
CLEC12A_Rat
CLEC_like1_Echidna
CLEC12A_Dog
CLEC4A_Cow
CLEC4A_Chimpanzee
CLEC4A_Human
CLEC4A_Dog
CLEC4A2_Mouse
CLEC4A_Rat
CLEC4A4_Mouse
CLEC4A2_Rat
CLEC4A3_Mouse
CLEC4A3_Rat
CLEC4A1_Mouse
CLEC4A1_Rat
CLEC_like5_Platypus
CLEC1_Mouse
CLEC1_Rat
CLEC1_Opossum
CLEC1_Human
CLEC1_Chimpanzee
CLEC1_Dog
CLEC1_Cow
CLEC_like3_Echidna
CLEC_like7_Echidna
CLEC_like4_Echidna
CLEC2_Cow
CLEC2_Human
CLEC2_Chimpanzee
CLEC2_Mouse
CLEC2_Rat
CLEC9A_Human
CLEC9A_Chimpanzee
CLEC9A_Rat
CLEC9A_Mouse
CLEC6A_Cow
CLEC_like10_Platypus
CLEC_like2_Echidna
CLEC_like11_Echidna
CLEC4B1_Mouse
CLEC4B2_Mouse
CLEC4B2_Rat
CLEC6A_Mouse
CLEC6A_Human
CLEC6A_Chimpanzee
CLEC4E_Cow
CLEC_like5_Echidna
CLEC4E_Human
CLEC4E_Chimpanzee
CLEC4E_Dog
CLEC4E_Mouse
CLEC4E_Rat
CLEC4D_Dog
CLEC4D_Cow
CLEC4D_Rat
CLEC4E_Opossum
CLEC4D_Human
CLEC4D_Chimpanzee
CLEC4D_Mouse
CLEC4C_Human
CLEC4C_Chimpanzee
CLEC_like6_Echidna
CLEC_like10_Echidna
CLEC_like9_Echidna
CLEC12A_Chimpanzee
CLEC7A_Chimpanzee
CLEC7A_Human
CLEC7A_Mouse
CLEC7A_Dog
CLEC7A_Cow
CLEC_like8_Echidna
CLEC_like6_Platypus
CLEC_like7_Platypus
CLEC_like11_Platypus
CLEC_like2_Platypus
OLR1_Cow
OLR1_Mouse
OLR1_Rat
CLEC_like4_Platypus
OLR1_Dog
CLEC_like1_Platypus
CLEC12B_Dog
OLR1_Human
OLR1_Chimpanzee
CLEC_like3_Platypus
CLEC9A_Dog
CLEC_like9_Platypus
CLEC2_Opossum
CLEC_like8_Platypus
CLEC7A_Rat

Online Resource 6 Phylogenetic analysis of Dectin family genes. Predicted protein sequences from multiple mammalian species were aligned in MEGAX using MUSCLE and Neighbour-joining phylogenetic trees were generated using the p-distance method with 1000 bootstrap repeats and pairwise deletion

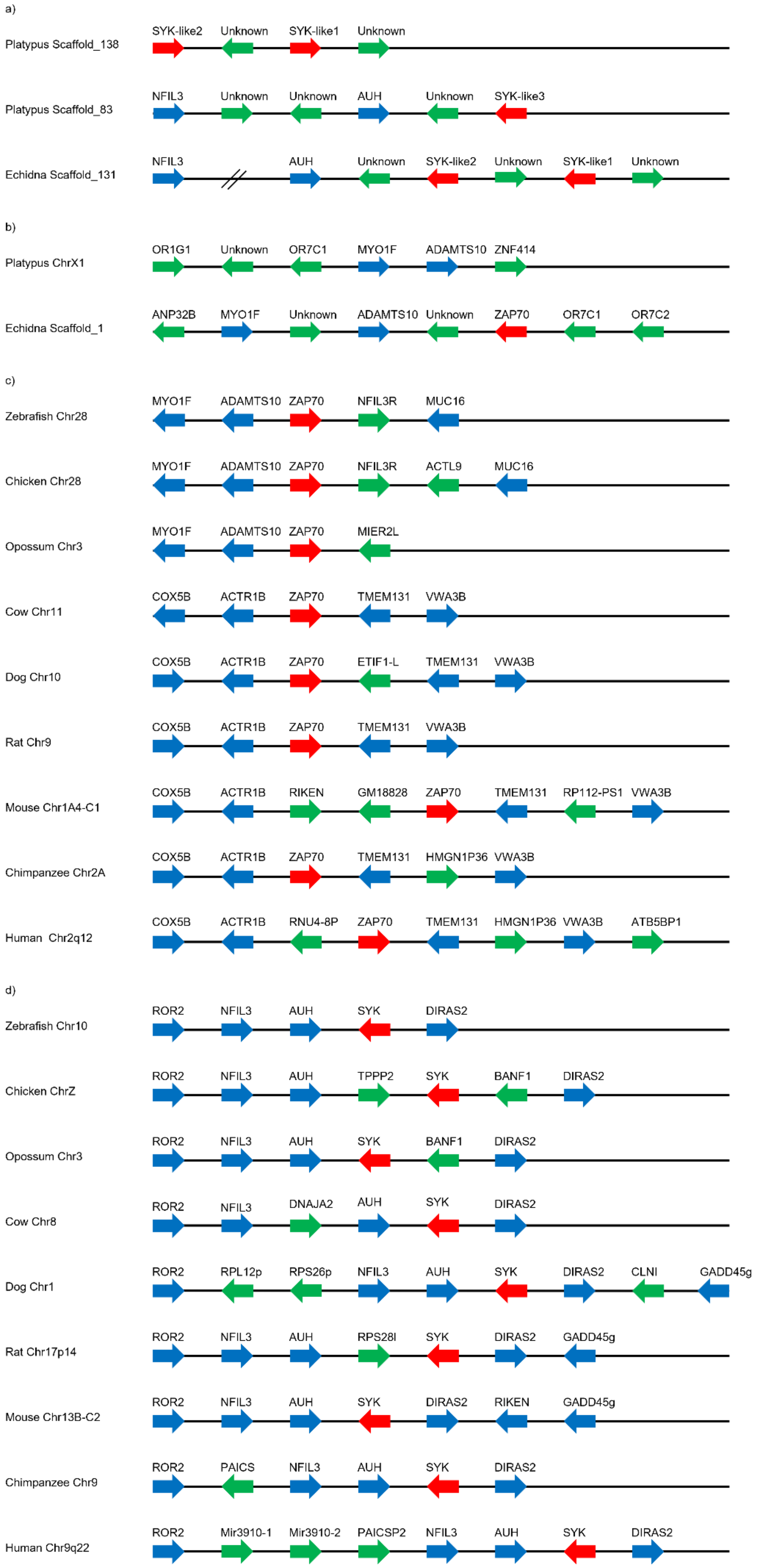

a)
Platypus Scaffold_138
SYK-like2 Unknown SYK-like1 Unknown
Platypus Scaffold_83
NFIL3 Unknown Unknown AUH Unknown SYK-like3
Echidna Scaffold_131
NFIL3 AUH Unknown SYK-like2 Unknown SYK-like1 Unknown
b)
Platypus ChrX1
OR1G1 Unknown OR7C1 MYO1F ADAMTS10 ZNF414
Echidna Scaffold_1
ANP32B MYO1F Unknown ADAMTS10 Unknown ZAP70 OR7C1 OR7C2
c)
Zebrafish Chr28
MYO1F ADAMTS10 ZAP70 NFIL3R MUC16
Chicken Chr28
MYO1F ADAMTS10 ZAP70 NFIL3R ACTL9 MUC16
Opossum Chr3
MYO1F ADAMTS10 ZAP70 MIER2L
Cow Chr11
COX5B ACTR1B ZAP70 TMEM131 VWA3B
Dog Chr10
COX5B ACTR1B ZAP70 ETIF1-L TMEM131 VWA3B
Rat Chr9
COX5B ACTR1B ZAP70 TMEM131 VWA3B
Mouse Chr1A4-C1
COX5B ACTR1B RIKEN GM18828 ZAP70 TMEM131 RP112-PS1 VWA3B
Chimpanzee Chr2A
COX5B ACTR1B ZAP70 TMEM131 HMGN1P36 VWA3B
Human Chr2q12
COX5B ACTR1B RNU4-8P ZAP70 TMEM131 HMGN1P36 VWA3B ATB5BP1
d)
Zebrafish Chr10
ROR2 NFIL3 AUH SYK DIRAS2
Chicken ChrZ
ROR2 NFIL3 AUH TPPP2 SYK BANF1 DIRAS2
Opossum Chr3
ROR2 NFIL3 AUH SYK BANF1 DIRAS2
Cow Chr8
ROR2 NFIL3 DNAJA2 AUH SYK DIRAS2
Dog Chr1
ROR2 RPL12p RPS26p NFIL3 AUH SYK DIRAS2 CLNI GADD45g
Rat Chr17p14
ROR2 NFIL3 AUH RPS28l SYK DIRAS2 GADD45g
Mouse Chr13B-C2
ROR2 NFIL3 AUH SYK DIRAS2 RIKEN GADD45g
Chimpanzee Chr9
ROR2 PAICS NFIL3 AUH SYK DIRAS2
Human Chr9q22
ROR2 Mir3910-1 Mir3910-2 PAICSP2 NFIL3 AUH SYK DIRAS2

Online Resource 7 Chromosomal arrangements for *SYK* (a and d) and *ZAP70* (b and c) for multiple mammalian species with chicken and zebrafish as non-mammalian outgroups. Arrows indicate orientation. Red arrows are genes of interest, blue arrows are conserved between species and green arrows are species specific. The genes shown as unknown represent genes identified by GENSCAN that lack any apparent homologue in human. Distances not to scale

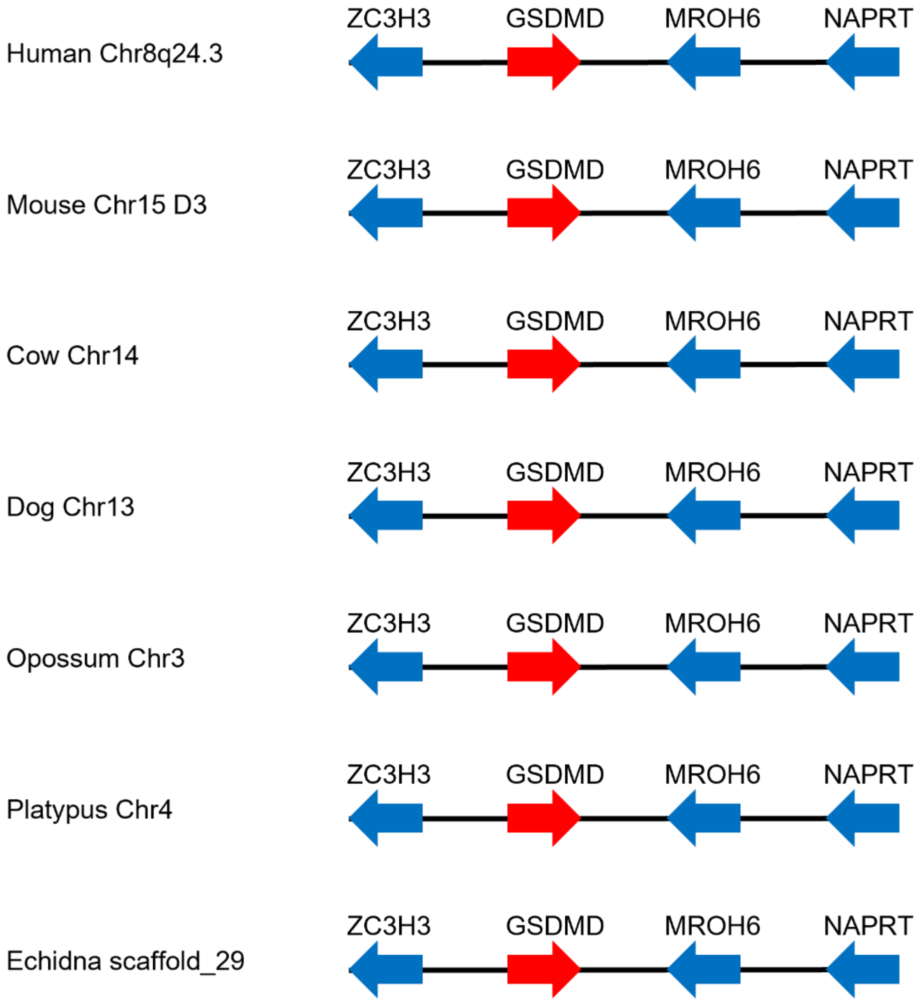


Online Resource 8: *Gsdmd* and flanking genes in multiple mammalian species. *Zc3h3*, *Mroh6* and *Naprt* flank *Gsdmd* in all mammalian species examined. The red arrow represents the gene of interest, and the blue arrows represent conserved gene synteny. Distances not to scale

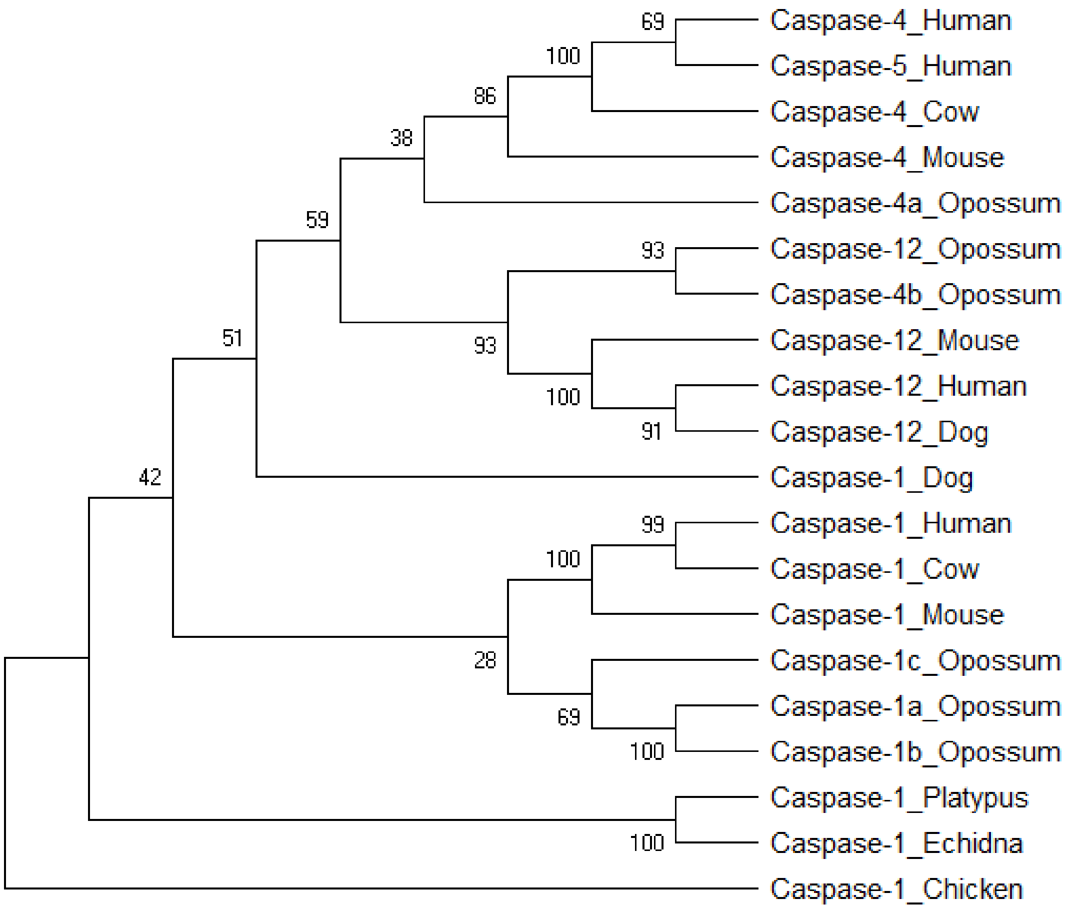


Online Resource 9: Phylogenetic analysis of inflammatory caspases in multiple mammalian species with chicken as an outgroup. Monotreme inflammatory caspases form their own cluster while dog *Caspase-1* is located separate to the therian *Caspase-1* cluster. Opossum *Casapse-4b* is located within the therian *Caspase-12* cluster. Predicted protein sequences from multiple mammalian species and chicken as an outgroup were aligned in MEGAX using MUSCLE and Neighbour-joining phylogenetic trees were generated using the p-distance method with 1000 bootstrap repeats and pairwise deletion

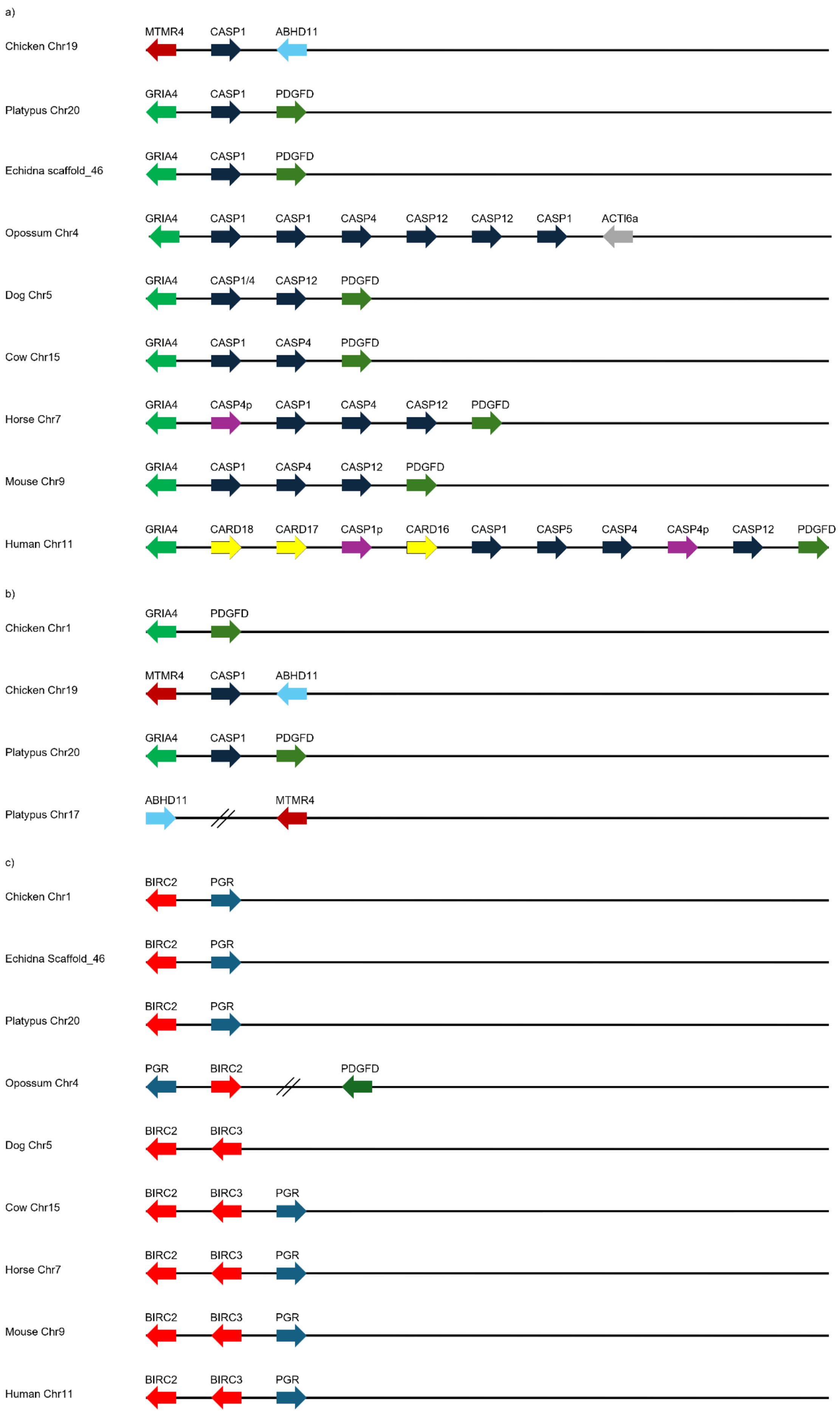

a)
Chicken Chr19
MTMR4 CASP1 ABHD11
Platypus Chr20
GRIA4 CASP1 PDGFD
Echidna scaffold_46
GRIA4 CASP1 PDGFD
Opossum Chr4
GRIA4 CASP1 CASP1 CASP4 CASP12 CASP12 CASP1 ACTl6a
Dog Chr5
GRIA4 CASP1/4 CASP12 PDGFD
Cow Chr15
GRIA4 CASP1 CASP4 PDGFD
Horse Chr7
GRIA4 CASP4p CASP1 CASP4 CASP12 PDGFD
Mouse Chr9
GRIA4 CASP1 CASP4 CASP12 PDGFD
Human Chr11
GRIA4 CARD18 CARD17 CASP1p CARD16 CASP1 CASP5 CASP4 CASP4p CASP12 PDGFD
b)
Chicken Chr1
GRIA4 PDGFD
Chicken Chr19
MTMR4 CASP1 ABHD11
Platypus Chr20
GRIA4 CASP1 PDGFD
Platypus Chr17
ABHD11 MTMR4
c)
Chicken Chr1
BIRC2 PGR
Echidna Scaffold_46
BIRC2 PGR
Platypus Chr20
BIRC2 PGR
Opossum Chr4
PGR BIRC2 PDGFD
Dog Chr5
BIRC2 BIRC3
Cow Chr15
BIRC2 BIRC3 PGR
Horse Chr7
BIRC2 BIRC3 PGR
Mouse Chr9
BIRC2 BIRC3 PGR
Human Chr11
BIRC2 BIRC3 PGR

Online Resource 10 a) Expansion of the Caspase-1 subfamily cluster. The mammalian *Caspase-1* is located in a different gene environment to chicken *Caspase-1*. Opossum has a large expansion of the gene cluster with duplications of *Caspase-1* and *Caspase-12*. There is also an inversion of the region containing the *Birc* genes and an insertion following the caspase cluster. Dog has a fusion of *Caspases -1* and *-4* while cow has a *Caspase-12* deletion. Both human and horse have a *Caspase-4-like* pseudogene. Human also has a *Caspase-1* pseudogene which is potentially a Card gene. b) Caspase-1 relocation following mammalian divergence. The *Caspase-1* gene has been identified on chicken chr19 and platypus chr20. The genes flanking chicken *Caspase-1* are different to those found flanking platypus *Caspase-1* which can be found in chicken in the same order but without *Caspase-1* on chr1. This shows that following avian divergence *Caspase-1* moved to a different chromosomal region in mammals. c) Chromosomal location of *Birc* genes. Chicken, monotremes and opossum genomes contain only a single Birc gene: *Birc2*. All eutherian species examined contained 2 genes: *Birc2* and *Birc3*. Distances not to scale
